# Identification of Microorganisms by Liquid Chromatography-Mass Spectrometry (LC-MS^1^) and *in silico* Peptide Mass Data

**DOI:** 10.1101/870089

**Authors:** Peter Lasch, Andy Schneider, Christian Blumenscheit, Joerg Doellinger

**Affiliations:** Robert Koch-Institute, ZBS6 - Proteomics and Spectroscopy, Seestraße 10, Berlin, D-13353, Germany

**Keywords:** Identification of Microorganisms, Mass Spectrometry, Proteomics, LC-MS^1^

## Abstract

Over the past decade, modern methods of mass spectrometry (MS) have emerged that allow reliable, fast and cost-effective identification of pathogenic microorganisms. While MALDI-TOF MS has already revolutionized the way microorganisms are identified, recent years have witnessed also substantial progress in the development of liquid chromatography (LC)-MS based proteomics for microbiological applications. For example, LC-tandem mass spectrometry (LC-MS^2^) has been proposed for microbial characterization by means of multiple discriminative peptides that enable identification at the species, or sometimes at the strain level. However, such investigations can be very time-consuming, especially if the experimental LC-MS^2^ data are tested against sequence databases covering a broad panel of different microbiological taxa.

In this proof of concept study, we present an alternative bottom-up proteomics method for microbial identification. The proposed approach involves efficient extraction of proteins from cultivated microbial cells, digestion by trypsin and LC-MS measurements. MS^1^ data are then extracted and systematically tested against an *in silico* library of peptide mass data compiled in house. The library has been computed from the UniProt Knowledgebase Swiss-Prot and TrEMBL databases and comprises more than 12,000 strain-specific *in silico* profiles, each containing tens of thousands of peptide mass entries. Identification analysis involves computation of score values derived from spectral distances between experimental and *in silico* peptide mass data and compilation of score ranking lists. The taxonomic positions of the microbial samples are then determined by using the best-matching database entries. The suggested method is computationally efficient – less than two minutes per sample - and has been successfully tested by a set of 19 different microbial pathogens. The approach is rapid, accurate and automatable and holds great potential for future microbiological applications.

## 2. INTRODUCTION

Rapid and reliable identification of pathogenic bacteria is of vital importance in many areas of public health and is relevant also in the food industry and for biodefense. In the context of clinical microbiology, a large variety of very different techniques, among them biochemical, serological, chemotaxonomic, and more recently spectroscopic, spectrometric and genomic tools are routinely utilized. For example, mass spectrometry-based techniques, such as matrix-assisted laser desorption/ionization time-of-flight mass spectrometry (MALDI-TOF MS) have emerged as invaluable tools for accurate and cost-effective identification of microorganisms in today’s routine clinical microbiology (Seng et al., 2009; Nomura, 2015; Schubert and Kostrzewa, 2017; Welker et al., 2019). The MALDI-TOF MS approach allows obtaining the genus and species identity of unknown samples by matching microbial mass spectra against spectral libraries collected from microorganisms with a known taxonomic identity. While identification is most reliably achieved at the species level, the question of whether MALDI-TOF MS is suitable for identification and discrimination below the species level is still controversially discussed by the scientific community (Sandrin et al., 2013; Demirev et al., 2016).

While a large number of studies convincingly demonstrate successful discrimination and identification of pathogenic bacteria by MALDI-TOF MS at the species level, there is also ample evidence for limitations of the taxonomic resolution, particularly at the infraspecies level and when dealing with differentiation of genetically closely related species (Lasch et al., 2014; Sousa et al., 2015; Rodrigues et al., 2016; Grenga et al., 2019). For example, differentiation between *Escherichia coli* and *Shigella* (He et al., 2010; Paauw et al., 2015) or of *Bacillus cereus* and *Bacillus anthracis* (Dybwad et al., 2013; Lasch et al., 2015) required additional measures beyond the standard microbial identification workflow, such as custom reference libraries, higher levels of standardization, and/or sophisticated data analysis concepts (machine learning, etc.). In these studies the reduced discriminatory power of MALDI-TOF MS has been attributed to the restricted m/z range (m/z 2000-20,000) and the dependence from mass patterns produced by a sub-proteome of small, abundant and basic proteins, mainly ribosomal subunit proteins (Rodrigues et al., 2016). Since the molecular evolution of these proteins is rather slow, ribosomal proteins are supposed to offer only limited taxonomic specificity. Further limiting factors of the MALDI-TOF MS method are the relatively low resolution, resulting in decreased selectivity and a reduced dynamic sensitivity, i.e. a lowered detectability of protein signals over a wide concentration range (Grenga et al., 2019).

In contrast to MALDI-TOF MS, liquid chromatography-tandem MS (LC-MS^2^) generally detects a large number of signals at very high resolution with very high mass precision in a single run (Grenga et al., 2019). Shotgun proteomic methods observe proteolytic cleavage products, often tryptic peptides, instead of intact proteins. This enables MS data collection with high analytical sensitivity. Moreover, coupling of mass spectrometry with chromatographic separation (LC) has shown to increase the dynamic sensitivity and allows sensitive detection also of low abundance peptides. Finally, LC-MS^2^ is much less restricted to classes of proteins with specific physicochemical properties. Even though proteomic techniques are still complex, rather cost-intensive and limited for use by well-equipped laboratories, the many advantages of LC-MS have led to an increasing number of activities aiming at evaluating potential applications of LC-MS in microbiology (Jabbour et al., 2010; Jabbour et al., 2011; Tracz et al., 2013; Berendsen et al., 2017).

Various groups have used shotgun proteomics for the classification and identification of pathogenic microorganisms. For example, a proteomics-based workflow for bacterial identification has been suggested by Dworzanski which involved construction of a bacterial proteome database from bacterial genomes, LC-MS^2^ data acquisition from digested bacterial cell extracts, identification of tryptic peptides and sequence-to-bacterium assignments (Dworzanski et al., 2006). The approach has been later utilized to determine the relatedness among strains of *B. cereus* sensu stricto, *B. thuringiensis* and *B. anthracis* by estimating fractions of shared peptides derived from a prototype database (Dworzanski et al., 2010). LC-MS^2^ has been also used by Tracz and coworkers to identify Biosafety Level 3 bacteria (Tracz et al., 2013). Sequence data from tryptic microbial peptides were obtained and employed for Mascot searches against a database containing concatenated protein sequences derived from microbial genomes. Identification of bacterial species was carried out by summing up matches from unique and degenerated (shared) peptides found per concatenated sequence; a post-culture analysis time of less than 8 hours has been reported.

Another alternative based on LC-MS has been proposed by Jabbour. Bacterial samples were lysed and subjected to tryptic digestion followed by LC-MS^2^ (Jabbour et al., 2010). Subsequently, peptides were identified and matched against databases. Bacteria were then identified based on the assessment of unique peptides obtained by an algorithm called BACid.

Comparison between microbial protein sequence data obtained by bottom-up tandem MS and reference databases was also performed by Boulund and colleagues. The proposed analysis pipeline (TCUP) not only returned specific genes of reference genomes that matched with peptide sequences determined by LC-MS^2^, but provided also the relative abundances of individual bacteria identified in a given mixed culture (Boulund et al., 2017). In this way, TCUP allowed typing and characterizing pathogenic bacteria from pure cultures and to estimate the relative abundances of individual microbial species from mixed microbial samples (Boulund et al., 2017). In the same year Berendsen and co-workers suggested a generic LC-MS^2^ method for the identification of microorganisms from positive blood cultures (Berendsen et al., 2017). A LC-MS compatible sample preparation method was developed that enabled accurate identification of bacteria grown in blood culture flasks to species level based on LC-MS^2^ bottom-up proteomics, data base searches and matching with taxon-specific discriminative peptides.

Advantages of the LC-MS^2^-based approaches for bacterial identification outlined above are the excellent accuracy of identification, high taxonomic resolution, universal applicability to the ever-growing numbers of known microbes and the ability to identify bacteria from mixtures, e.g. in polymicrobial infections. At the other hand the comparatively high computational requirements have to be mentioned. Since the time required for peptide identification correlates with the number of entries contained in sequence databases, computation time can be saved by restricting the size of the database, for example by using genus-specific databases. However, database restrictions contradict the use of shotgun proteomics as an unbiased approach for microbial identification. Another important limitation of LC-MS^2^-based approaches is the severely reduced accuracy and sensitivity of common search algorithms when extensive protein sequence databases are used. The large search space impedes the identification of true peptide matches within large numbers of similar sequences.

With this proof-of-concept study, we introduce an alternative, easy-to use and computational less demanding approach for microbial identification. The proposed method is based on bottom-up proteomics as the analytical technique and involves acquisition of LC-MS data from cultivated bacteria. MS^1^ data are extracted and tested against a database with more than 12,000 strain-specific synthetic mass profiles which has been compiled in-house using public protein databases (UniProtKB). We demonstrate that the MS^1^ information can be used for rapid and accurate taxonomic identification, at least at the species level, and discuss possibilities to combine the suggested analysis pipeline with known MS^2^-based analysis methods in microbiology.

## 3. MATERIALS AND METHODS

### Microbial strains

The performance and accuracy of the proposed method for microbial identification was tested using 19 well-characterized bacterial strains which were predominantly obtained from established strain collections such as DSM (Deutsche Sammlung von Mikroorganismen), ATCC (American Type Culture Collection) and NCTC (National Collection of Type Cultures). Strains E 125, E 131 and E153 of *Burkholderia thailandensis* originated from the strain collection at the Robert Koch-Institute (RKI) (Lasch et al., 2018). An overview of the microbial strains and species utilized is given in Tab. 1.

**Table 1.**

| No | Genus, Species | Strain name(s) |
| --- | --- | --- |
| 1 | *Acinetobacter baumannii* | DSM 30007 |
| 2 | *Bacillus cereus* | DSM 31 |
| 3 | *Bacillus velezensis* | DSM 23117 / FZB42 |
| 4 | *Burkholderia cepacia* | ATCC 25416 |
| 5 | *Burkholderia thailandensis* | DSM 13277 |
| 6 | *Burkholderia thailandensis* | E125* |
| 7 | *Burkholderia thailandensis* | E131* |
| 8 | *Burkholderia thailandensis* | E153 |
| 9 | *Burkholderia thailandensis* | LMG 20219 |
| 10 | *Burkholderia oklahomensis* | DSM 21774 |
| 11 | *Citrobacter freundii* | DSM 30039 |
| 12 | *Enterococcus faecalis* | DSM 20371 |
| 13 | *Escherichia coli* | DSM 3871 |
| 14 | *Mycobacteroides abscessus* | DSM 44196 |
| 15 | *Pseudomonas aeruginosa* | ATCC 27853 |
| 16 | *Staphylococcus epidermidis* | DSM 1798 |
| 17 | *Staphylococcus aureus* | DSM 20231 / NCTC 8532 |
| 18 | *Vibrio cholerae* | NIH 41 |
| 19 | *Yersinia pseudotuberculosis* | DSM 8992 |

Overview of the microbial species and strains used in this study. Abbreviations: DSM – Deutsche Sammlung von Mikroorganismen; ATCC – American Type Culture Collection; NCTC – National Collection of Type Cultures; LMG – Belgian Coordinated Collections of Microorganisms, Universiteit Gent – Laboratorium voor Microbiologie, NIH – National Institute of Health, \* (Lasch et al., 2018)

### Cultivation

Bacteria were streaked under sterile conditions on solid culture media by an inoculation loop and incubated for 48 hours. Strains prepared by the STrap protocol (see below and Tab. 1) were cultivated according to species-specific cultivation requirements on tryptic soy agar (TSA, ReadyPlate TSA ISO, Merck Life Science) or Middlebrook agar produced in-house. Cells from these cultures were harvested by scraping; approximately 10 µL of microbial material was then washed three times by 1 mL of ice-cold phosphate buffered salt solution and centrifuged at 4000 g at 4°C for 5 min. All other bacteria were cultured on Casein-Soy-Peptone (Caso, Oxoid, Wesel, Germany) agar plates under aerobic conditions for 24 h at 37°C and further processed according to the protocol *Sample Preparation by Easy Extraction and Digestion* (SPEED, see below and Tab. 1 for details).

### Sample preparation by suspension trapping (STrap)

Cells were suspended in 5 % sodium dodecyl sulfate (SDS), 20 mM dithiothreitol (DTT), 50 mM Tris/HCl buffer, pH 7.6 (sample/buffer 1:10 (v/v)), incubated at 95°C for 10 min and further sonicated for 15 cycles à 30 s at high intensity level and 4°C using Bioruptor^®^Plus (Diagenode, Liege, Belgium). Protein concentrations were determined by measuring the tryptophan fluorescence at an emission wavelength of 350 nm using 295 nm for excitation with an Infinite^®^ M1000 PRO microplate reader (Tecan, Männedorf, Switzerland) according to (Wisniewski and Gaugaz, 2015). Samples were further processed using S-Trap™ mini filters according to the STrap sample preparation protocol (Zougman et al., 2014) and the manufacturer’s instructions (Protifi, Huntington NY, USA).

### Sample preparation by the SPEED protocol

SPEED protocol-based preparation of microbial cells was carried as outlined in (Doellinger et al., 2018). Briefly, bacterial cells were suspended in trifluoroacetic acid (TFA, Uvasol^®^ for spectroscopy, Merck, Darmstadt, Germany) in a sample/TFA ratio of 1:4 (v/v) and heated at 70°C for 3 min. Acid extract samples were then neutralized with 2M TrisBase using the tenfold volume of TFA solution and further incubated at 95°C for 5 min after adding tris (2-carboxyethyl)phosphine (TCEP) to a final concentration of 10 mM and 2-chloroacetamide (CAA) to a final concentration of 40 mM. Protein concentrations were determined by turbidity measurements at 360 nm (1 AU = 0.79 µg/µL) using a GENESYS™ 10S UV-Vis Spectrophotometer (Thermo Fisher Scientific, Waltham, Massachusetts, USA) and adjusted to 0.25 µg/µL using a 10:1 (v/v) mixture of 2M TrisBase and TFA. The solution was afterwards diluted 1:5 (v/v) with water. Proteins were digested for 20 h at 37°C using Trypsin Gold, Mass Spectrometry grade (Promega, Fitchburg, USA) at an enzyme/protein ratio of 1:50 (w/w).

### Peptide desalting

Peptides were desalted using 200 µL StageTips™ packed with three Empore™ SPE Disks C18 (3M Purification, Inc., Lexington, USA) according to (Ishihama et al., 2006) and concentrated using a vacuum concentrator. Samples were resuspended in 20 µL 0.1 % formic acid (FA) and peptides were quantified by measuring the absorbance at 280 nm using a Nanodrop 1000 device (Thermo Fisher Scientific, Rockford, IL, USA).

### Nano-LC tandem MS measurements

Nano-LC tandem MS (nLC-MS^2^) measurements of pathogenic bacteria were partly carried out within the scope of other proteomics projects. Desalted digests were analyzed on an EASY-nanoLC 1200 device (Thermo Fisher Scientific, Bremen, Germany) coupled online to a Q Exactive™ Plus mass spectrometer (Thermo Fisher Scientific, Bremen, Germany). 1 µg peptides were separated on a 50 cm Acclaim™ PepMap™ column (75 μm inner diameter, i.d., 100 Å C18, 2 μm; Thermo Fisher Scientific, Bremen, Germany) using a linear 120 min gradient of 3 to 28 % acetonitrile (ACN) in 0.1 % FA at 200 nL/min flow rate. Column temperature was kept at 40°C using a butterfly heater (Phoenix S&T, Chester, PA, USA). The mass spectrometer was operated in a data-dependent manner in the m/z range of 300 – 1,650. Full scan spectra were recorded with a resolution of 70,000 using an automatic gain control (AGC) target value of 3 × 10^6^ with a maximum injection time of 20 ms. Up to the 10 most intense 2+ – 5+ charged ions were selected for higher-energy c-trap dissociation (HCD) with a normalized collision energy (NCE) of 25 %. Fragment spectra were recorded at an isolation width of 2 Th and a resolution of 17,500@200 m/z using an AGC target value of 1 × 10^5^ with a maximum injection time of 50 ms. The minimum AGC MS^2^ target value was set to 1 × 10^4^. Once fragmented, peaks were dynamically excluded from precursor selection for 30 s within a 10 ppm window. Peptides were ionized using electrospray with a stainless steel emitter, i.d. 30 µm, (Proxeon, Odense, Denmark) at a spray voltage of 2.0 kV and a heated capillary temperature of 275°C.

### Extraction of MS^1^ data from MS^2^ data sets, pre-processing of MS^1^ data

Peptide feature detection in MS^1^ data was carried out using the Minora algorithm of the Proteome Discoverer software v. 2. 2.0388 (Thermo-Fisher Scientific) with default settings. Peptide feature text files were then produced by *LCMS-Biotyping*.*vvf*, a home-written method compiled for obtaining the following parameters from the peptide features: MS^1^ peak positions (in m/z units) with the respective ion charge state, retention time, normalized abundancy and signal-to-noise ratio (SNR). These parameters were stored in tab-separated text files and imported by a custom-designed Matlab function *readlcmstxtfile* (Matlab, The Mathworks, Natick, USA). As part of the *parseuniprot* toolbox (see below) this command line tool supports import of peptide feature text files obtained from LC-MS^1^ data and performs data pre-processing, including molecular weight (MW) determination by considering charge states, detecting and removing peak entries originating from oxidized peptides (mass shift +15.99491 Da) as well as from peptides with deamidated glutamine or asparagines residues (mass shift +0.98402 Da). Spectral pre-processing involved furthermore partially removing (underweighting) of low intensity and low MW peaks; based on the principle that the relevance of a specific peak for subsequent identification analysis co-varies with its intensity and MW values (see below). As the result of pre-processing, experimental LC-MS^1^ data of a sample is collapsed into a single MS^1^ mass peak list, which contains the filtered peptide mass data. Such peak lists are in the following referred to as LC-MS^1^ peak lists.

### Compilation of the in silico peptide mass database

Assembly of the *in silico* library of microbial peptide mass data was done utilizing information from the UniProt Knowledgebase (UniProtKB). In particular, UniProtKB/Swiss-Prot and UniProtKB/TrEMBL text files, *uniprot_sprot_bacteria*.*dat*.*gz* (size: 211,465,997 byte, date: Dec 10, 2018) and *uniprot_trembl_bacteria*.*dat*.*gz* (60,987,643,246 byte, Dec 09, 2018) were downloaded from ftp://ftp.uniprot.org and unpacked. Extraction of relevant data i.e. of protein sequences and metadata was carried out by means of *parseuniprot*, a collection of specifically developed Matlab-based functions (The Mathworks). The *parseuniprot* toolbox is available from the principal author of this study on request and will be publicly available after publication of this study.

Compilation of the *in silico* database and identification analysis was carried using a dual processor Dell Precision T7500 workstation (2× Intel 3.46 GHz x5690 CPU with 6 cores each) that was equipped with 160 GB RAM, a Samsung 512 GB solid-state drive (850 Pro) and additional 20 TB hard drive space. A 64-bit version of Microsoft Windows 7 operating system and Matlab-based software developed in-house (*parseuniprot*, MicrobeMS) running under a 64-bit version of Matlab R2014a (The Mathworks, Natick, USA) were utilized. MicrobeMS and *parseuniprot* required the Matlab Statistics (v. 9.0), Bioinformatics (v. 4.4) and Parallel Computing (v. 6.4) toolboxes of Matlab. The output of the *parseuniprot* toolbox i.e. the *in silico* library of microbial peptide mass data can be stored as a so-called *pkf* file. Files of this type are compatible with MicrobeMS, the identification software utilized (Lasch, 2019b); a detailed description of the *pkf* file format is provided in the MicrobeMS wiki (Lasch, 2019a).

### Bacterial Identification analysis

For bacterial identification analysis we used the MicrobeMS toolbox which has been initially developed in the context of biotyping pathogenic microorganisms by MALDI-TOF MS (Lasch et al., 2014; Lasch et al., 2015). This Matlab-based toolbox can be freely downloaded from the MicrobeMS website (compiled version for Microsoft Windows 64-bit, registration required). Within the scope of the present study the MicrobeMS toolbox was specifically adapted in order to meet the higher memory and larger computational requirements.

For identification analyses LC-MS^1^ peak lists (see above) were first imported via the *muf* data format that is specific for MicrobeMS (Lasch, 2019a). Inter-spectral distances between LC-MS^1^ peak lists and *in silico* peptide mass profiles were then obtained utilizing the function *compare with DB* of MicrobeMS.

Bar coded MS^1^ test spectra were constructed from LC-MS^1^ peak lists using MW bins of a relative width of 1.2 ppm. As distance metrics, variance-scaled Pearson’s product momentum correlation coefficients (Pareto scaling 0.25) were selected, whereby data between 2000 and 5500 Da served as inputs. The calibration range factor was set to a value of 5 giving the total number of calibration factor variations of 125, see MicrobeMS wiki for details (Lasch, 2019a). In MicrobeMS correlation coefficient-based inter-spectral distances are converted to score values between 0 and 1000. This is achieved on the basis of linear scaling, whereby a score of 1000 can only be achieved if the LC-MS^1^ test and the database spectrum match entirely, i.e. in cases of identical spectra. A score of zero, on the other hand, is obtained only when any correlation is absent.

In MicrobeMS the score values for each test spectrum are arranged in a ranking list and the best matching data-base entries are used to determine the taxonomic identity of the strain investigated. This approach is not new and has been used for many years in infrared, Raman, or MALDI-TOF MS identification software solutions such as Bio-Rad’s KnowItAll, Bruker’s MALDI Biotyper, or Biomerieux’s Saramis / VITEK MS. In the current implementation of MicrobeMS, the score ranking list is provided in a HTML format where the top 30 best-matching database records are displayed for each LC-MS^1^ test spectrum analyzed (Lasch, 2019a), see also supporting information, SI).

## 4. RESULTS

### Overview of the identification analysis workflow

In this paper we present a computational pipeline which is suitable for identification of pathogenic microorganisms from bottom-up mass spectrometry (MS) data. An overview of the proposed sequence of analysis steps is presented in Fig. 1. First steps of the proposed approach include cultivation of pathogenic bacteria under standardized conditions followed by sample preparation using validated protocols. Microbial materials are subsequently characterized by means of nanoLC-tandem MS (nLC-MS^2^). First steps of analysis involve extraction of MS^1^ and MS^2^ data. After pre-processing, the MS^1^ data are systematically compared against entries of an *in silico* database, which has been compiled beforehand from the totality of microbial protein entries present in the UniProt Knowledgebase, i.e. in UniProtKB/Swiss-Prot and UniProtKB/TrEMBL databases. The output of this comparison is a ranking list of score values derived from inter-spectral correlation, or distance values. Score lists ideally provide information on the taxonomic identity of the microorganism under investigation. The MS^1^-derived genus and species information is thought to be useful for further characterization of the bacteria. In cases where the results of MS^1^ analyses are incomplete, the species assignments could be useful to guide and assist subsequent analyses of MS^2^ data, for example to identify bacteria at the infraspecies level (strain typing).

**Figure 1.**
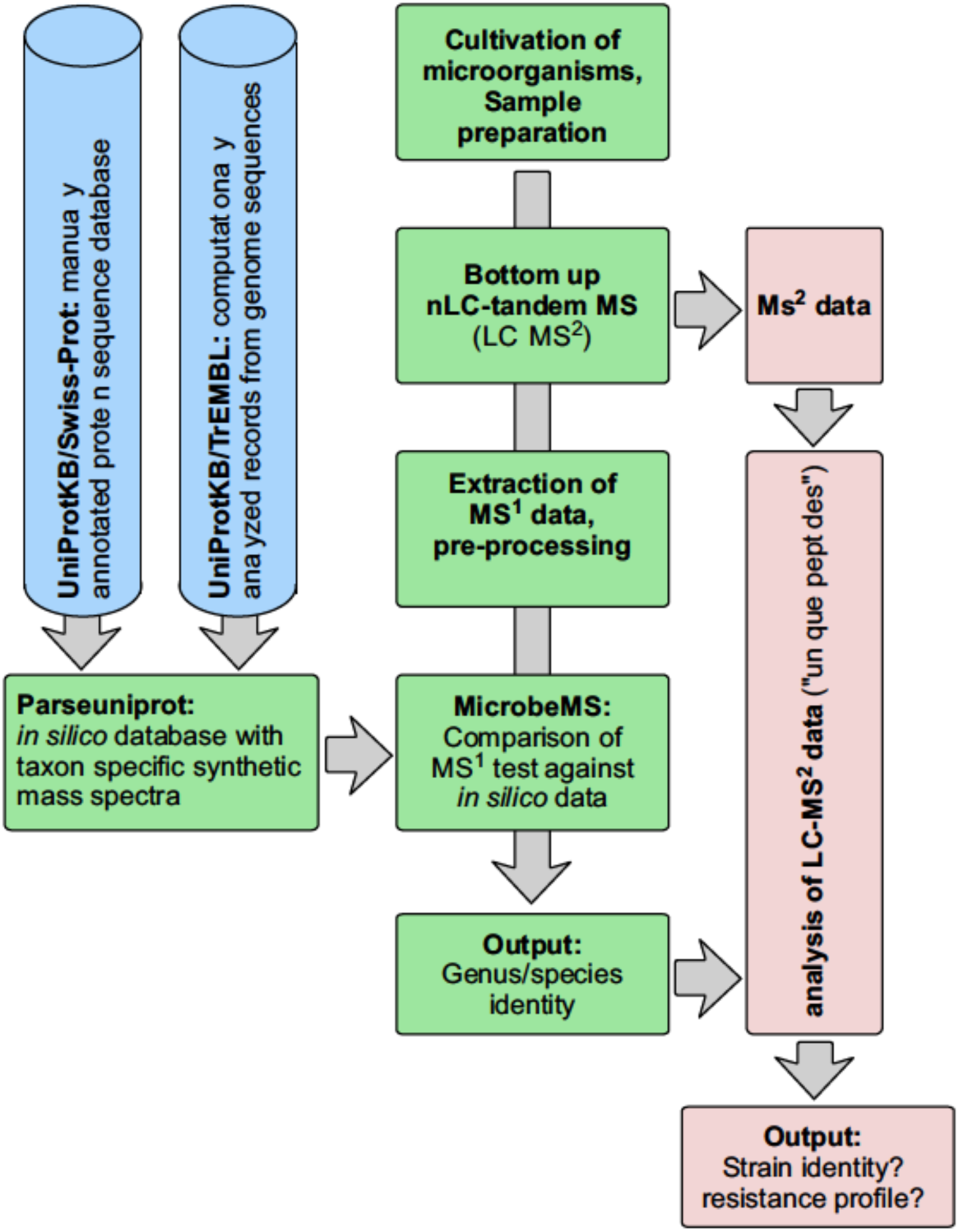
Overview of the proposed LC-MS based microbial identification workflow. Microorganisms of unknown identity are cultivated under standardized conditions and colony material is prepared using established sample preparation protocols for shotgun proteomics. Mass spectrometry data are then acquired using tandem MS (MS^2^). MS^1^ data are extracted and pre-processed for subsequent comparison against a library of strain-specific synthetic mass profiles obtained from UniProtKB/Swiss-Prot and UniProtKB/TrEMBL protein sequence data. A ranking list of correlation or inter-spectral distance values (i.e. of scores) is established. These lists provide information on the taxonomic identity of the studied microorganism. The genus and/or species level information can be used for more detailed characterization by alternative methods, for example from MS^2^ data (see red boxes to the right). The latter approaches are, however, no subjects of the present study.

### Compilation of an in silico database of microbial peptide mass data

Compilation of a database of taxon-specific peptide mass lists from microbial proteome data constituted the main challenge in this proof-of-concept study. Development of such a database required downloading the complete bacterial UniProtKB/Swiss-Prot library with reviewed and manually annotated proteins and the UniProtKB/TrEMBL proteome data (unreviewed, computationally analyzed proteins) from ftp://ftp.uniprot.org. Both mutually exclusive databases were merged and the protein sequence information and scattered metadata were extracted, further processed, re-sorted and stored in a format the microbial identification software can read. The specified data analysis steps were carried out by means of the *parseuniprot* toolbox, a Matlab-based analysis pipeline specifically developed at RKI. In addition to supplementary command-line Matlab functions, e.g. for merging the UniProtKB/Swiss-Prot and UniProtKB/TrEMBL databases or for creating and modifying taxonomy white lists (see below), the *parseuniprot* toolbox comes with three major functions, (i) *readdat*, (ii) *resort* und (iii) *modfeat* which are consecutively executed and whose analysis steps build upon each other.

Fig.2 schematically illustrates the sequence of the analysis steps *readdat, resort* and *modfeat*. The first function *parseuniprot/readdat* has been employed for data import from UniProtKB files uniprot_sprot_bacteria.dat (UniProtKB/Swiss-Prot) and uniprot_trembl_bacteria.dat (UniProtKB/TrEMBL) and for generating a first Matlab structure array. In this array each protein is mapped to an array element whereby each element comprises specific fields that contain protein sequence and metadata as well as protein, proteome and taxonomic identifiers (IDs). In order to reduce the overall data amount and to exclude entries with unclear taxonomic assignment from further analyses *parseuniprot/readdat* has also support for “whitelisting” organisms. The white list was obtained from UniProtKB and initially contained 22,529 taxon entries with genus, species and strain designation, protein numbers, and the corresponding UniProtKB taxonomic and proteome IDs. The raw white list was scanned for keywords like ‘*unidentified*’, ‘*candidatus*’,’ *sp*.’, ‘*uncultured*’, etc. for subsequent removal of respective entries.

**Figure 2.**
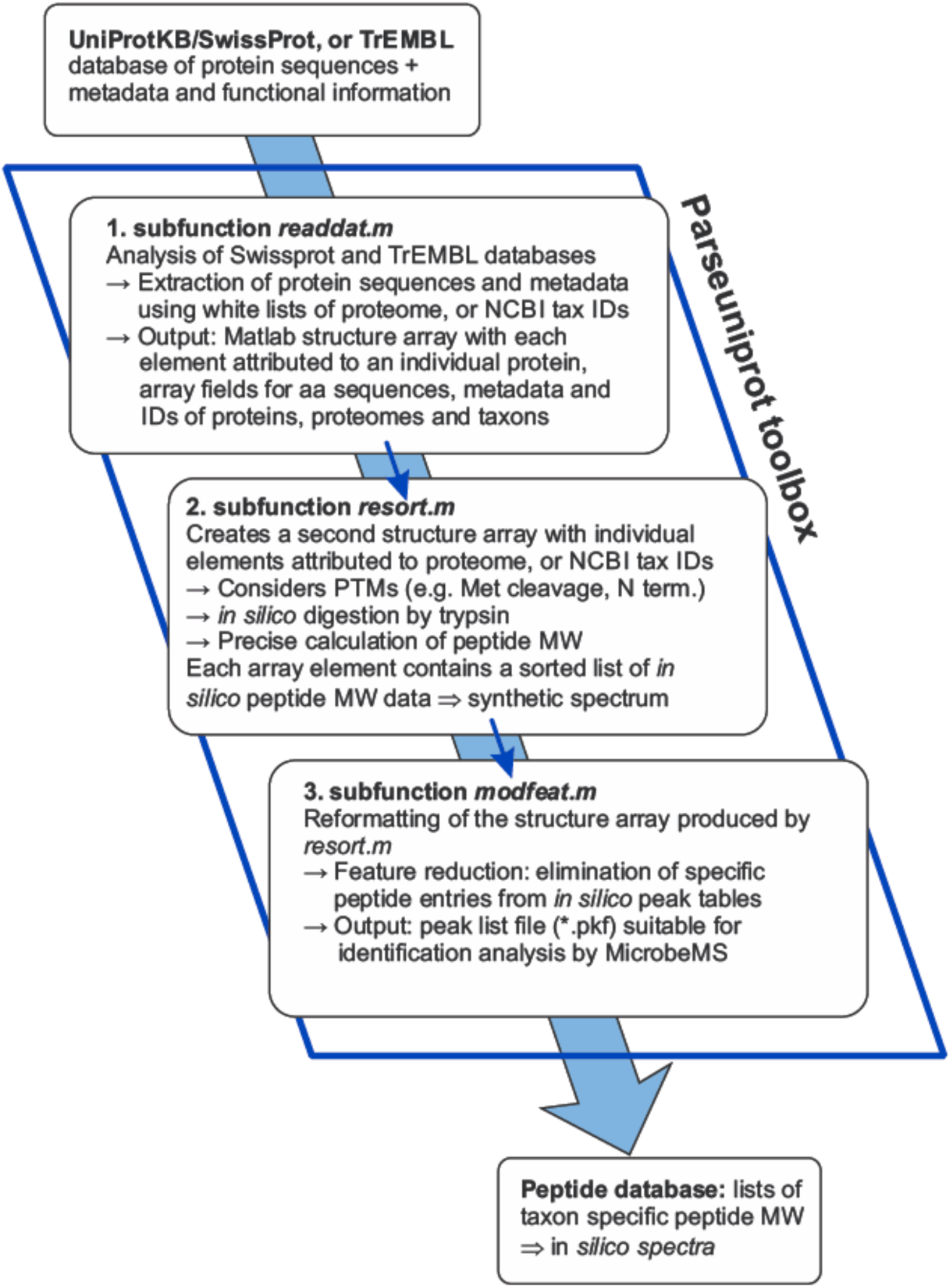
Schematic workflow for generating *in silico* databases from UniProtKB/Swiss-Prot and/or UniProtKB/TrEMBL protein sequence data. The Matlab toolbox *parseuniprot* represents a proteomic pipeline in which three main internal functions, *readdat, resort* and *modfeat* are consecutively executed. The function *readdat* converts the content from structured text files available from ftp://ftp.uniprot.org into Matlab structure arrays that contain the complete information needed to compile *in silico* MS^1^ databases. These structure arrays are subsequently processed by the functions *resort* and *modfeat*; the output of the *parseuniprot* pipeline is a peak list file comprising strain-specific synthetic (*in silico*) peptide mass profiles suitable for computer-based comparison with LC-MS^1^ test spectra.

The data structure produced by *parseuniprot/readdat* served as input for the second function, *parseuniprot/resort* and contained altogether sequences and metadata from 47,537,746 microbial proteins. Subsequent analyses involved disregarding strain entries with less than 1200 proteins, identification/assignment of posttranslational modification (PTM) sites such as cleavage of N-terminal methionine and *in silico* digestion by trypsin using the rule “cleave at the carboxyl termini of lysine or arginine, except when either is followed by proline”. No missed cleavages were allowed. Afterwards, exact peptide molecular weight (MW) determination was conducted from amino-acid sequences of tryptic peptides whereas it was considered that carbamidomethylcysteines (160.030647 Da) are present in CAA treated samples instead of cysteine (103.009185 Da). Although available through Matlab’s Bioinformatics Toolbox, PTM profiling and MW determination routines had to be re-written for performance reasons and, after careful testing, integrated into the *parseuniprot* toolbox. All calculated MW values of tryptic peptides were then re-indexed according to proteome or UniProtKB IDs. As a result, each proteome or taxon endpoint in question was associated with the respective peptide entries which permitted compiling strain-specific mass lists after sorting, filtering and cleansing. These mass lists are in the following referred to as synthetic peptide mass spectra, or microbial *in silico* (MS^1^) spectra. These data were allowed for use in further analyses in cases where the peptide MW was larger than 780 Da and if spectra contained more than 15,000 and less than 280,000 peptide entries. After application of these criteria, the *in silico* database for LC-MS^1^ based identification contained 723,820,940 individual peptide entries sorted in altogether 12,044 taxon-specific synthetic spectra which corresponds to approximately 60,000 peptide peaks per spectrum.

The third function of the analysis pipeline, *parseuniprot/modfeat*, was designed for feature selection, specifically to allow filtering of presumptive non-specific peptide mass entries with the objective to increase the accuracy of subsequent identification analysis. In the present implementation of the *parseuniprot/modfeat* function selected peptide entries are removed on the basis of the following considerations: Peaks of a given synthetic MS^1^ spectrum should ideally be taxon-specific, i.e. should preferentially be entirely absent, or at least rarely present in the remaining database spectra. To this end, an algorithm has been tested which served the purpose to determine throughout the whole database the relative peak frequency at each MW interval. Subsequently, peak frequency values are extracted for all MW intervals at which the first database spectrum exhibits a peak. Peak frequencies are then ranked in descending order and MW intervals above a certain frequency threshold are removed. The procedure has to be repeated for each of the 12,044 database spectra. Tests revealed that the overall accuracy of identification markedly increased if peaks above the 90th frequency percentile are disregarded from further analyses. In this way, 10% of the peaks carrying presumably non-specific information are removed from the *in silico* database. Peak lists and selected metadata were subsequently stored in the *pkf* file format, see above and (Lasch, 2019a).

For the future, it is planned to test and implement more advanced feature selection approaches. Genetic algorithms, for example, could be advantageously applied to identify combinations of taxon-specific peptide markers and could thus contribute to improve the accuracy of identification. Fig. S1 shows a screenshot with the graphical user interface of the current version of the *parseuniprot* toolbox.

### Spectral pre-processing

Pre-processing of raw experimental data aims at increasing the robustness and accuracy of subsequent quantitative or classification analysis (Lasch, 2012; Tsai et al., 2016). In the context of the present study, the strategy of pre-processing LC-MS^1^ spectra was inspired by the following ideas:

Firstly, the number of experimentally determined MS^1^ peaks varied between 60,000 and 90,000. It is reasonable to assume that relevant fractions of these peaks carry non-specific information with some of them arising from chemically modified peptides. For the sake of simplicity, we have proposed that the intensity of such peaks is lower on average compared to intensities from unmodified peptides. Thus, underweighting low intensity features from MS^1^ spectra was assumed to have a positive impact on the accuracy of identification. Secondly, high MW peptides are thought to be somewhat more specific with regard to pathogen identification than short peptides with a lower MW. An important objective of data pre-processing was therefore to eliminate low intensity peaks in low MW regions at a higher rate than in high MW regions. Thirdly, peaks from chemically modified peptides, i.e. from oxidized or deamidated species, are not represented in the synthetic database and should be thus identified and removed.

The results of pre-processing MS^1^ data are exemplarily illustrated in Fig. 3. As shown by the LC-MS^1^ test spectrum of *Enterococcus faecalis* DSM 20371, pre-processing preferentially removed low intensity peaks in the low MW region. The upper row of Fig. 3 illustrates MS^1^ peak density functions of raw (blue bars) and of pre-processed data (red bars) whereas the x-axes encode the log10-scaled intensity (left), or the MW value (right). The blue shaded area indicates the lower and upper bounds of the MW region utilized for identification analysis by MicrobeMS (see below). More data exemplifying the strategy of spectral pre-processing are provided in the supporting information (see Figs. S2 and S3)

**Figure 3.**
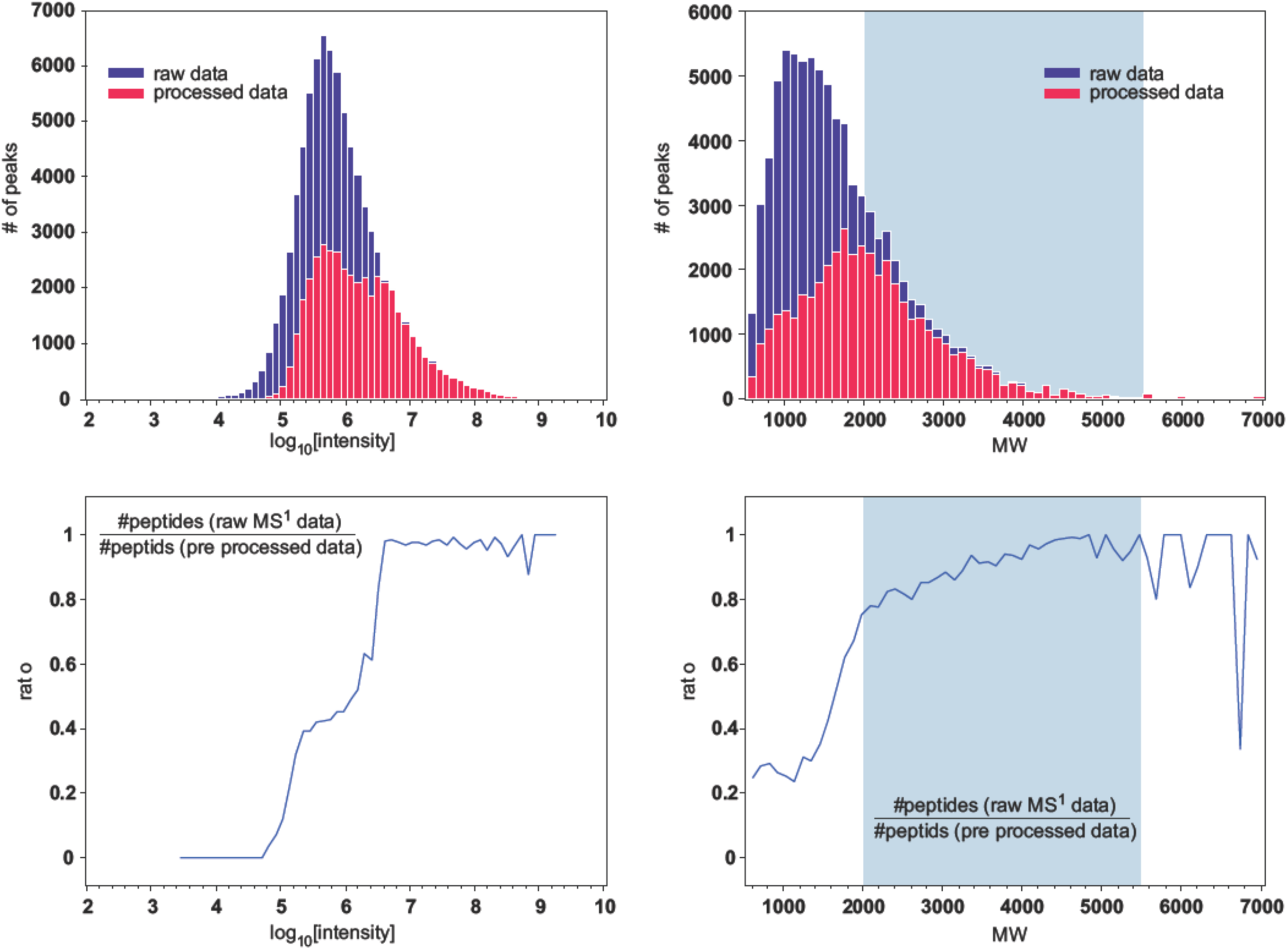
Pre-processing and feature selection of LC-MS^1^ data. MS^1^ peak data were acquired from a culture of *Enterococcus faecalis* DSM 20371; sample preparation has been carried according to the SPEED sample preparation protocol. (Doellinger et al., 2018) **Top row:** histogram bar chart of log_10_ scaled MS^1^ peak intensities (left) and the molecular weight (MW) distribution (right) of peaks after feature detection by the Minora algorithm (=original* data, blue bars) and after pre-processing and feature selection by *readlcmstxtfile* (processed data, red bars). Total number of peaks in original / processed MS^1^ data: 82843 / 42559 Number of oxidized / deamidated peptides found and removed: 389 / 329 **Lower row**: ratio between the number of peaks present in processed and original MS^1^ data as a function of peak intensity (log_10_ scaled, left), or of the MW (right). Pre-processing was carried out by *readlcmstxtfile*, a Matlab function developed in house. This function has been designed to preferentially remove low intensity signals in the low MW region (< 2000 Da). The blue shaded area indicates the MW range used for identification analysis by MicrobeMS (2000 – 5500 Da).

### MS^1^ identification analysis by using the silico database

The results of identification analysis by means of LC-MS^1^ test data and the *in silico* database are given in Tab. 2. The table summarizes information on the sample identity, i.e. of the genus, species and strain identities of the microorganisms tested (cf. column 2) and the sample preparation technique applied (SPEED for samples #1 - #13) and STrap for samples #14 - #21, see column 1). Furthermore, columns 3, 4 and 5 list the three top-ranked bacteria as determined by MicrobeMS: Column 3 details their genus, species and strain identities while columns 4 and 5 depict the corresponding proteome IDs and the correlation-derived score values, respectively.

**Table 2.**

| Sample no. / Protocol | Genus, species and strain identity | Top database hits | Proteome ID | Scores |
| --- | --- | --- | --- | --- |
| #1<br>SPEED | *Bacillus velezensis*<br>DSM 23117 / FZB42 | 1. *B. velezensis* ATR2<br>2. *B. velezensis* DSM 23117 / FZB42<br>3. *B. vallismortis* NBIF-001 | UP000228760<br>UP000001120<br>UP000193819 | 172.1654<br>165.3249<br>164.1915 |
| #2<br>SPEED | *Staphylococcus epidermidis*<br>DSM 1798 | 1. *S. epidermidis* DSM 1798 / ATCC 12228<br>2. *S. epidermidis* M23864:W2(grey)<br>3. *S. epidermidis* ATCC 35984 / RP62A | UP000001411<br>UP000004733<br>UP000000531 | 260.842<br>254.5784<br>250.1004 |
| #3<br>SPEED | *Staphylococcus aureus* DSM 20231 / NCTC 8532 | 1. *S. aureus* Newman<br>2. *S. aureus* USA300<br>3. *S. aureus* NCTC 8325 | UP000006386<br>UP000001939<br>UP000008816 | 213.046<br>212.0801<br>211.2254 |
| #4<br>SPEED | *Pseudomonas aeruginosa*<br>ATCC 27853 | 1. *P. aeruginosa* LIM1410<br>2. *P. aeruginosa* LIM1680<br>3. *P. aeruginosa* MH19 | UP000247072<br>UP000246728<br>UP000043988 | 197.6311<br>195.8522<br>188.7818 |
| #5<br>SPEED | *Enterococcus faecalis*<br>DSM 20371 | 1. *E. faecalis* CH116<br>2. *E. faecalis* RMC1<br>3. *E. faecalis* EnGen0426 | UP000013839<br>UP000013621<br>UP000023191 | 203.0399<br>202.8773<br>202.6593 |
| #6<br>SPEED | *Citrobacter freundii*<br>DSM 30039 | 1. *C. freundii* GED7749C<br>2. *E. coli* ISC11<br>3. *C. freundii* MRSN 12115 | UP000070673<br>UP000019194<br>UP000032554 | 178.2909<br>172.3027<br>169.3774 |
| #7<br>SPEED | *Mycobacteroides abscessus*<br>DSM 44196 | 1. *M. abscessus* ATCC 19977 / DSM 44196<br>2. *M. abscessus* subsp. *abscessus* 1135<br>3. *M. abscessus* subsp. *bolletii* 909 | UP000007137<br>UP000184667<br>UP000184881 | 144.387<br>131.1567<br>128.8442 |
| #8<br>SPEED | *Burkholderia oklahomensis*<br>DSM 21774 | 1. *B. oklahomensis* NCTC 13388<br>2. *B. pseudomallei* MSHR346<br>3. *B. pseudomallei* 1710b | UP000254484<br>UP000002031<br>UP000002700 | 217.199<br>149.3277<br>149.2699 |
| #9<br>SPEED | *Burkholderia thailandensis*<br>E153 | 1. *B. thailandensis* DSM 13276 / LMG 20219<br>2. *B. thailandensis* FDAARGOS_243<br>3. *B. pseudomallei* 1710b | UP000001930<br>UP000235972<br>UP000002700 | 214.043<br>204.8531<br>164.4795 |
| #10<br>SPEED | *Burkholderia thailandensis*<br>E131 | 1. *B. thailandensis* DSM 13276 / LMG 20219<br>2. *B. thailandensis* FDAARGOS_243<br>3. *B. pseudomallei* 1710b | UP000001930<br>UP000235972<br>UP000002700 | 204.9895<br>196.8191<br>159.7089 |
| #11<br>SPEED | *Burkholderia thailandensis*<br>E125 | 1. *B. thailandensis* DSM 13276 / LMG 20219<br>2. *B. thailandensis* FDAARGOS_243<br>3. *B. pseudomallei* 1710b | UP000001930<br>UP000235972<br>UP000002700 | 202.7592<br>195.6465<br>161.2297 |
| #12<br>SPEED | *Burkholderia thailandensis*<br>LMG 20219 | 1. *B. thailandensis* DSM 13276 / LMG 20219<br>2. *B. thailandensis* FDAARGOS_243<br>3. *B. pseudomallei* 1710b | UP000001930<br>UP000235972<br>UP000002700 | 203.342<br>192.4561<br>161.0222 |
| #13<br>SPEED | *Burkholderia thailandensis*<br>DSM 13277 | 1. *B. thailandensis* DSM 13276 / LMG 20219<br>2. *B. thailandensis* FDAARGOS_243<br>3. *B. pseudomallei* 668 | UP000001930<br>UP000235972<br>UP000002153 | 204.233<br>194.5373<br>159.5722 |
| #14<br>STrap | *Yersinia pseudotuberculosis*<br>DSM 8992 | 1. *Y. pestis* SCPM-O-DNA-17 (I-2457)<br>2. *Y. pseudotuberculosis* IP 31758<br>3. *Y. pseudotuberculosis* IP32953 | UP000239693<br>UP000002412<br>UP000001011 | 157.8063<br>140.4254<br>140.2100 |
| #15<br>STrap | *Vibrio cholerae*<br>NIH 41 | 1. *V. cholerae* ATCC 39541 / Ogawa 395<br>2. *V. cholerae* MO10<br>3. *V. cholerae* 2740-80 | UP000000249<br>UP000004687<br>UP000003017 | 158.1642<br>153.7766<br>153.4644 |
| **#16**<br>STrap | *Pseudomonas aeruginosa*<br>ATCC 27853 | 1. *P. aeruginosa* ATCC 15692 / DSM 22644<br>2. *P. aeruginosa* LIM4285<br>3. *P. aeruginosa* MH19 | UP000002438<br>UP000246712<br>UP000043988 | 114.6227<br>112.7038<br>112.6669 |
| **#17**<br>STrap | *Bacillus cereus*<br>DSM 31 | 1. *B. cereus* BDRD-Cer4<br>2. *B. cereus* FSL R5-860<br>3. *B. thuringiensis* T13001 | UP000001373<br>UP000019049<br>UP000001911 | 139.0604<br>138.2032<br>137.4324 |
| **#18**<br>STrap | *Burkholderia cepacia*<br>ATCC 25416 | 1. *B. cepacia* ATCC 25416<br>2. *B. cepacia* FDAARGOS_345<br>3. *B. reimsis* BE51* | UP000263019<br>UP000197040<br>UP000252458 | 141.5042<br>135.7973<br>134.2624 |
| **#19**<br>STrap | *Mycobacteroides abscessus*<br>DSM 44196 | 1. *M. abscessus* ATCC 19977 / DSM 44196<br>2. *M. abscessus* subsp. *abscessus* 1135<br>3. *M. abscessus* subsp. *bolletii* 1513 | UP000007137<br>UP000184667<br>UP000023351 | 129.9005<br>119.8615<br>118.8616 |
| **#20**<br>STrap | *Acinetobacter baumannii*<br>DSM 30007 | 1. *A. baumannii* SDF<br>2. *A. baumannii* ATCC 19606 / DSM 30007<br>3. *A. baumannii* 1295743 | UP000001741<br>UP000005740<br>UP000020595 | 156.7779<br>153.8487<br>147.1175 |
| **#21**<br>STrap | *Escherichia coli*<br>DSM 3871 | 1. *E. coli* ATCC 27325 / DSM 5911 / K12<br>2. *E. coli* B / BL21-DE3<br>3. *S. sonnei* Ss046 | UP000000318<br>UP000002032<br>UP000002529 | 128.7893<br>128.2228<br>127.1311 |

Identification results of a microbial LC-MS<sup>1</sup> test data set comprising 19 different strains from 11 genera and 15 species. Samples #1-#13 were prepared by the SPEED proteomic sample preparation protocol developed at RKI (Doellinger et al., 2018). The STrap (Zougman et al., 2014) method was utilized for preparing proteomic samples from strains listed at lines 14-21. Of note: two strains, *P. aeruginosa* ATCC 27853 and *M. abscessus* DSM 44196 were prepared and measured twice. \* refers to a not validly described species (Esmaeel et al., 2018) (see text for details).

Test results summarized in Tab. 2 generally suggest a high level of identification accuracy and demonstrate that LC-MS^1^ data from microbial samples can be meaningfully and successfully queried against synthetic peptide mass spectra. As an illustration, identification was accurate at the genus and species level in 20 out of 21 cases and in 5 cases there was even an agreement at the strain level. Furthermore, complete matching between the species identity of the test sample and three top scored database entries has been determined in 9 out of 21 instances. In view of the facts that only some of the characterized strains are contained in the *in silico* database and that not all microbial species are represented by three or more database entries, these first findings point out the large potential of MS^1^ data analysis for rapid microbial identification. Detailed analysis of the *Burkholderia pseudomallei* group test subset which comprised 5 different strains of *Burkholderia thailandensis* and one strain of *Burkholderia oklahomensis* revealed further excellent test results (cf. samples #8-#13): almost perfect accuracy was obtained from this sample set considering the facts that (i) *Burkholderia pseudomallei* and *Burkholderia mallei* are the phylogenetically closest relatives of the latter two species (Vandamme and Eberl, 2018) and (ii) that the current version of the *in silico* library contained only one synthetic spectrum of *B. oklahomensis* and two profiles of *B. thailandensis*.

The complete results of identification tests of the *Burkholderia pseudomallei* group subset including data from biological and technical replicates are given in the supporting information (cf. Fig S4).

A closer examination of test results of samples #1 (*Bacillus velezensis*), #17 (*Bacillus cereus*) and #18 (*Burkholderia cepacia*) showed that genus and species assignments of the two top scored bacteria were accurate whereas bacteria listed at position 3 were either close relatives thereof, or strains with unclear species assignment. For example, in case of sample #1, *Bacillus vallismortis* was found at position 3. *B. vallismortis* is like *B. velezensis* a member of the *Bacillus subtilis* group (Fan et al., 2017). In the instances of samples #17 and #18 either a close relative of *B. cereus* was ranked at position 3 - *Bacillus thuringiensis* is a member of the *B. cereus* group, cf. (Helgason et al., 2000) - or *Burkholderia reimsis*, better known as *Burkholderia* sp. BE51 a bacterium with incomplete species description was listed at position 3, see (Esmaeel et al., 2018; Vandamme and Eberl, 2018) for details.

Identification results of samples #6 (*Citrobacter freundii*) and #21 (*Escherichia coli*) reveal that two out of the three top scored bacteria were correctly identified, including both top hits. Inspection of data from sample #21 suggested, however, that differentiation between *E. coli* and strains of *Shigella* is not straightforward, an observation which is substantiated by the special taxonomy of *E. coli* and *Shigella* and additionally reflected by findings made by other techniques, such as MALDI-TOF MS (Khot and Fisher, 2013; Paauw et al., 2015), LC-MS^2^-based approaches (Berendsen et al., 2017) or whole genome sequencing, WGS (Chattaway et al., 2017). In the instance of sample #21, we found 25 strain entries of *E. coli* and 5 entries from 3 different *Shigella* species (*S. sonnei*, 3 × *S. boydii* and *S. dysenteriae*) among the top 30 entries present in the score ranking list (data not shown). This and the finding of insignificant distances among scores in the ranking list suggest that the current version of the MS^1^ analysis pipeline might not be always suitable for reliable differentiation of *E. coli* and *Shigella*. This observation is supported by taxonomy data. For example, it has been stated that *Shigella* strains can be viewed from a genetic perspective as subpopulations within *E. coli* (Pupo et al., 2000; Paauw et al., 2015; Chattaway et al., 2017) and some studies even recommend re-classification of *Shigella* and *E. coli*, e.g. (Pupo et al., 2000; Alves et al., 2018).

True misidentification was observed only in a single instance, see sample #14. In this example *Yersinia pseudotuberculosis* has been identified as *Yersinia pestis* SCPM-O-DNA-17 (I-2457, top hit); strains of Y. *pseudotuberculosis* were, however, ranked at positions 2 and 3. As in the previous example, the analysis of the taxonomy is helpful to understand the particular test result. On a genomic level *Y. pestis* is known to be highly similar to the enteric pathogen *Y. pseudotuberculosis* (Achtman et al., 1999; Demeure et al., 2019). In fact, Y. pestis can be considered a clone of *Y. pseudotuberculosis* which has evolved only recently (Achtman et al., 1999; Califf et al., 2015). Moreover, the top scored strain of *Y. pestis* is a member of the subspecies *Y. pestis* ssp. *ulegeica* (Kislichkina et al., 2018). Strains of *Y. pestis ssp. ulegeica* belong to a branch which is from a phylogenetic point of view more closely related to the ancestor *Y. pseudotuberculosis* than members from other branches of *Y. pestis*, including recent strains from *Y. pestis* ssp. *pestis* (Kutyrev et al., 2018; Demeure et al., 2019).

### Influence of sample preparation on the accuracy of identification

In this paper, two different methods of microbial sample preparation, STrap (Zougman et al., 2014) and SPEED (Doellinger et al., 2018), were utilized. Although the relatively low number of tested samples does not allow a statistically valid comparison, the tests provide some preliminary indications with regard to the influence of sample preparation on the accuracy of identification.

As given by Tab. 2, the collected data involve 13 microbial samples processed by means of the SPEED method (samples #1-#13) while 8 samples were prepared using the STrap protocol (samples #14-#21). Two of the microbial samples, *Mycobacteroides abscessus* DSM 44196 and *Pseudomonas aeruginosa* ATCC 27853 were processed by either method, samples #7 and #4 (SPEED) and samples #16 and #19 (STrap).

Results from differently processed samples revealed that top score values of STrap samples varied between 110 and 160 whereas identification tests with SPEED processed samples led in 12 of 13 instances to scores > 170, among them 9 cases with scores > 200 (identical parameters of identification). Among SPEED samples, only one case with a score below 170 was observed (#7, *M. abscessus*, score of 144). Of note, the higher scores of SPEED processed samples do not appear to result in a significantly improved accuracy of identification: Data from *M. abscessus* and *P. aeruginosa* which are both prepared by STrap and SPEED demonstrate accurate identification at the species level in both instances. Of note this has been achieved despite the higher score values found in SPEED samples - 144 (SPEED) vs. 129 (STrap) in case of *M. abscessus* and 198 vs. 115 in *P. aeruginosa*, respectively. Of note that identification of *M. abscessus* was accurate in either sample even at the strain level.

In summary, it can be stated that higher score values were determined with SPEED samples, whereby the identification accuracy, however, is thought to benefit only slightly from this increase. Nevertheless, the data set used is rather small, so a more comprehensive test set would be required to support further conclusions.

## 5. DISCUSSION

In this proof-of-concept study, we evaluated the principal applicability of shotgun nanoLC - mass spectrometry (nLC-MS^1^) for microbial identification and explored the taxonomic resolution of the proposed method. To this end, an *in silico* database of microbial peptide MW data was constructed from UniProtKB resources which was then queried by a set of LC-MS^1^ test spectra obtained from microbial cells grown in pure cultures. The results of these queries are summarized in score ranking lists which are helpful to obtain insights into the taxonomic identity of the bacteria studied. It can be stated that the suggested approach is generally suitable for identifying bacteria at the genus and species level, and sometimes even at strain level (cf. Tab. 2 and supporting information). Despite these encouraging results, it should be also noted that in certain limited instances ambiguous identification results were found. Targeted screening for combinations of taxon-specific peptide features (feature selection) and a better quality of the underlying protein sequence database, particularly of UniProtKB/TrEMBL (database curation), are suggested as potential starting points to address the remaining shortcomings and to further improve the accuracy of identification.

### Computational considerations

With a size of the final *in silico* database of 1.44 GB, disk space requirements were only moderate. At the other hand, the computational time required for compilation from UniProtKB resource data was substantial: Approximately 30 h were needed using a Dell Precision T7500 workstation (see above). It should be noted, however, that the *in silico* database only needs to be constructed at longer intervals, for example every two months, so that the high computational requirements and long database construction times would become less important in a routine setup.

Similarly, determination of score values based on Pareto-scaled correlation coefficients between pre-processed MS^1^ spectra and *in silico* peptide MW lists are computationally intensive tasks and require powerful hardware. The Matlab implementation therefore involved parallel programming with fully vectorized code and multicore support, which was helpful to reduce analyses time down to less than 2 minutes per LC-MS^1^ test spectrum (Dell workstation). Precondition for short computation times is a sufficient amount of installed memory; approx. 160 GB RAM is the minimum requirement. Furthermore, within the scope of the present study the MicrobeMS toolbox has been ported from the original Windows 7 64-bit Matlab version to Linux (Debian 8.11) and tested on one of the RKI’s bioinformatics server (8× Intel Xeon E7-4890 v2@2.8 GHz, 120 cores, 1 TB RAM). This led to shorter analysis times (∽40 s per strain) whereby tests with 12, 15, 20, 40 and 60 cores revealed only minor improvements of computation times (not shown). Obviously identification analysis did not significantly benefit from utilization of more than 12 cores.

### Comparison with other MS methods for microbial identification

The workflow of the proposed method is schematically illustrated in Fig. 4. It is to be noted that the MS^1^-based analytical method resembles in many aspects the well-known MALDI-TOF MS identification technique. Despite the many similarities with the MALDI-TOF MS approach, such as the need to cultivate, or utilization of spectral databases and score ranking lists, there is one important difference: The proposed method does not require a spectra library acquired experimentally from pathogens of known identity. The database used in this study has been ultimately computer generated from genome data. This implies that an impressive number of bacterial strains – more than 12,000 at present – is already available for identification analysis purposes. This number exceeds the number of entries contained in the current version of Bruker’s MALDI Biotyper (MBT) CE *in vitro* diagnostics (CE-IVD) database. The MBT CE-IVD database has been developed over a number of years and contains in its present version V.9 (2019) 8326 MSP entries from bacteria and yeasts (Dewaele et al., 2019). This number nicely illustrates that the suggested LC-MS^1^ method directly benefits from the ever-increasing growth of genomic data: Database coverage of both, newly discovered and already known microbiological taxa can be improved with relatively little efforts.

**Figure 4.**
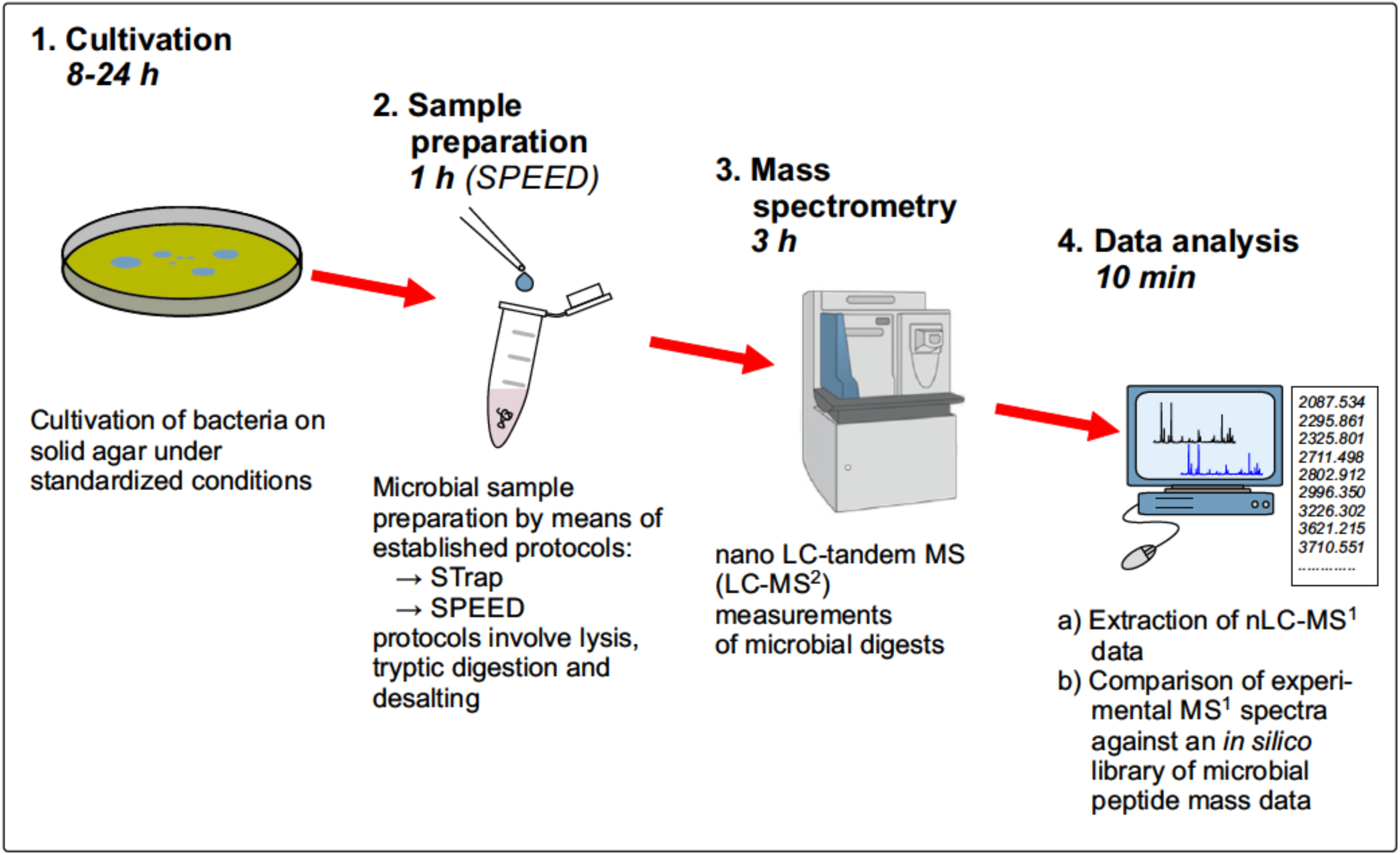
Schematic workflow for microbial identification based on nLC-MS^1^ spectra obtained from cultivated bacteria and *in silico* databases comprising strain-specific synthetic peptide mass profiles.

The significantly higher number of signals present in LC-MS data constitutes another important difference to MALDI-TOF MS. While MALDI-TOF MS usually detects limited numbers (< 150) of predominantly high-abundance proteins with housekeeping functions, such as basic ribosomal proteins, or nucleic acid-binding proteins (Ryzhov and Fenselau, 2001; Pineda et al., 2003; Dieckmann et al., 2008), usually in the m/z region between 2 and 20 kDa, LC-MS shotgun proteomics identifies usually more than 10,000 peptides from a single microbial sample. However, further studies are required to clarify whether the enormous increase of information contained in the LC-MS data actually leads to a better taxonomic resolution.

Current drawbacks of the LC-MS method are, above all, longer measurement times, higher instrument costs and a relatively low dissemination of LC-MS equipment and analysis concepts in clinical or food microbiology laboratories. However, it is to be expected that technological developments will help to reduce measurement times and it is anticipated that dedicated systems will reduce costs and thus improve dissemination of LC-MS technology.

Compared to published LC-MS^2^ methods to identify and classify microbial pathogens, the proposed approach offers a number of important advantages. Firstly, the computational requirements are significantly reduced due to the fact that identification of peptides and/or proteins is not necessary. Under our experimental conditions we have been able to obtain an identification result within less than 5 hours (excluding cultivation, cf. Fig. 4), whereby computational time requirements were negligible (less than 10 minutes). Secondly, the method does not rely on specifically identifying, i.e. discriminating (“unique”) peptides. This is important because the number of such peptides tends to diminish with the ever-growing number of peptides contained in future database versions. Thirdly, the simplicity of the proposed identification method allows to define scores in a straightforward manner and to adapt a well-established principle in the field of MS-based microbial diagnostics (score ranking lists). Internal discussions and an in-depth literature search led us to the conclusion that score definition from LC-MS^2^ data by using unique peptides constitutes a rather complex problem for which no universally accepted solution has yet been presented.

Major disadvantages compared to LC-MS^2^-based identification are the need for cultivation due to the higher requirements of sample purity. Although polymicrobial samples were not tested by us, it is reasonably clear that the presented approach is not applicable to the study of such samples. For the same reasons as MALDI-TOF MS, the proposed concept of MS^1^ data analysis does not allow reliable identification of individual microorganisms from complex microbial mixtures, such as faeces, or natural biofilm habitats. In addition, the method does not offer a direct way for detecting database errors, such as missing entries or only incomplete database coverage. While the absence of unique peptides in MS^2^–based identification analyses would indicate database gaps, MS^1^ score ranking lists do not necessarily provide direct evidences for database errors or poor (incomplete) coverage. However, such gaps are likely to be rare events, due to the large number of microbial species already contained in the presented very first database version.

The main advancement of the suggested LC-MS^1^ workflow to bacterial differentiation consists in the possibility for combination with LC-MS^2^-based analysis concepts. Such a hybrid approach would involve straightforward correlation analysis between LC-MS^1^ test spectra and synthetic peptide mass profiles and peptide identification analyses by the help of MS^2^ data, ideally from order, family, or genus specific sequence databases. It is expected that this concept is not only helpful to speed up analysis, but improves also the accuracy of identification as a whole.

## 6. CONCLUSIONS

This proof-of-concept study has demonstrated that identification analysis from LC-MS^1^ data represents a powerful technology that could drive improvements in bacterial identification. The technique utilizes *in silico* libraries which can be generated from publicly available proteome resources and does not require databases of experimental mass spectra. The proposed pipeline is easy to use, computationally efficient and freely available for both Linux and Windows operating systems. The taxonomic resolution of the method is promising, but improvements, such as well-curated databases, application of feature selection methods, better quality checks as well as rigorously conducted tests with large LC-MS^1^ data sets are needed to answer the question whether the observed inconsistencies are inherent limitations of the method or are simply due to the fact that the LC-MS^1^ approach is not yet fully developed.

## Abbreviations

ACN: acetonitrile;
AGC: automatic gain control;
DTT: dithiothreitol;
CAA: 2-chloroacetamide;
FA: formic acid;
ID: identifier;
MALDI-TOF: matrix-assisted laser desorption/ionization - time–of–flight;
MBT: MALDI Biotyper;
MS: mass spectrometry;
MW: molecular weight;
NCE: normalized collision energy;
ppm: parts per million;
RKI: Robert Koch-Institute;
SDS: sodium dodecyl sulfate;
SNR: signal-to-noise ratio;
SPEED: sample preparation by easy extraction and digestion;
STrap: suspension trapping;
TFA: trifluoroacetic acid;
TSA: tryptic soy agar;
TCEP: Tris(2-carboxyethyl)phosphine;
UniProtKB: UniProt Knowledgebase

## 7. ACKNOWLEDGEMENTS

The authors are thankful to Max Weydmann for conducting tandem LC-MS measurements of the microbial samples prepared by the STrap method.

## 8. AUTHOR CONTRIBUTION STATEMENT

PL and JD contributed conception and design of the study; AS and JD collected the data and performed the experiments and first steps of data pre-processing; PL wrote the code of the Matlab toolboxes parseuniprot and MicrobeMS and performed the data processing; PL wrote the first draft of the manuscript; AS, CB and JD wrote sections of the manuscript. All authors contributed to manuscript revision, read and approved the final version of the manuscript

## 9. CONTRIBUTION TO THE FIELD STATEMENT

Modern methods of mass spectrometry have emerged and allow reliable, fast and cost-effective identification of pathogenic microorganisms. For example, matrix-assisted laser desorption/ionization time-of-flight (MALDI-TOF) mass spectrometry (MS) has revolutionized the way pathogenic microorganisms are identified in today’s routine clinical microbiology. Furthermore, recent years have witnessed also substantial progress in the development of liquid chromatography-mass spectrometry (LC-MS) based proteomics for microbiological applications.

With this proof-of-concept study, we introduce an alternative technique for microbial identification using mass spectrometry. The proposed method is based on bottom-up proteomics and involves acquisition of LC-MS data from cultivated bacteria. Mass spectrometry data are processed and tested against an *in silico* database which can be compiled in-house using public protein sequence data. A first version of this database currently contains more than 12,000 strain-specific synthetic mass profiles. In our study we demonstrate accurate taxonomic identification by the suggested approach, at least at the species level, and discuss possibilities to combine our pipeline with existing LC-MS analysis concepts. The proposed method is rapid, simple and automatable and we foresee wide application potential for future microbiological applications.

