## Supporting Information for "Identification of Microorganisms by Liquid Chromatography-Mass Spectrometry (LC-MS^1^) and *in silico* Peptide Mass Data"

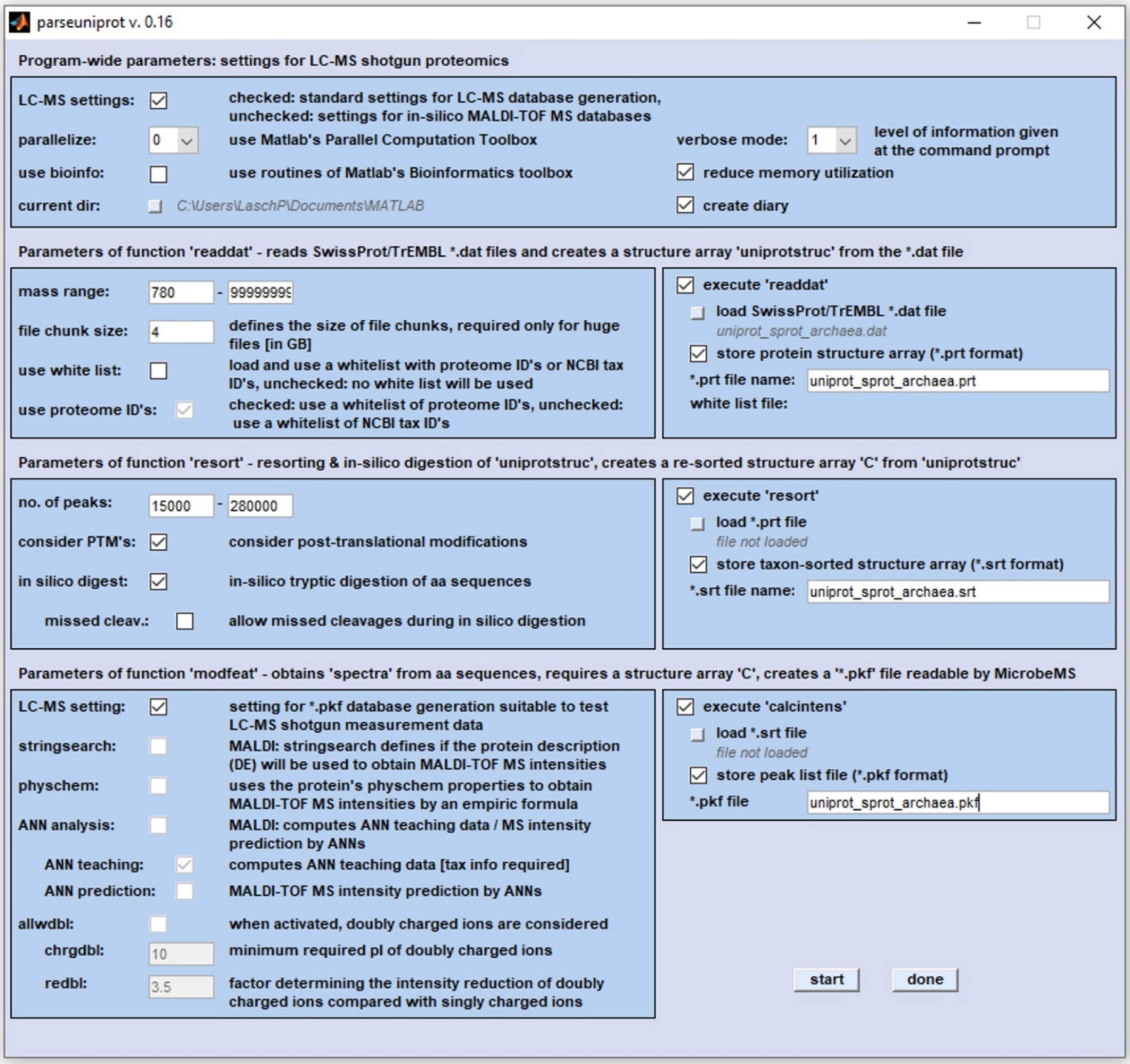


**Figure S1.**

Screenshot of the Matlab-based toolbox *parseuniprot* suitable for compilation of *in silico* databases comprising strain-specific synthetic peptide mass profiles from UniProtKB/Swiss-Prot and/or UniProtKB/TrEMBL protein sequence data.

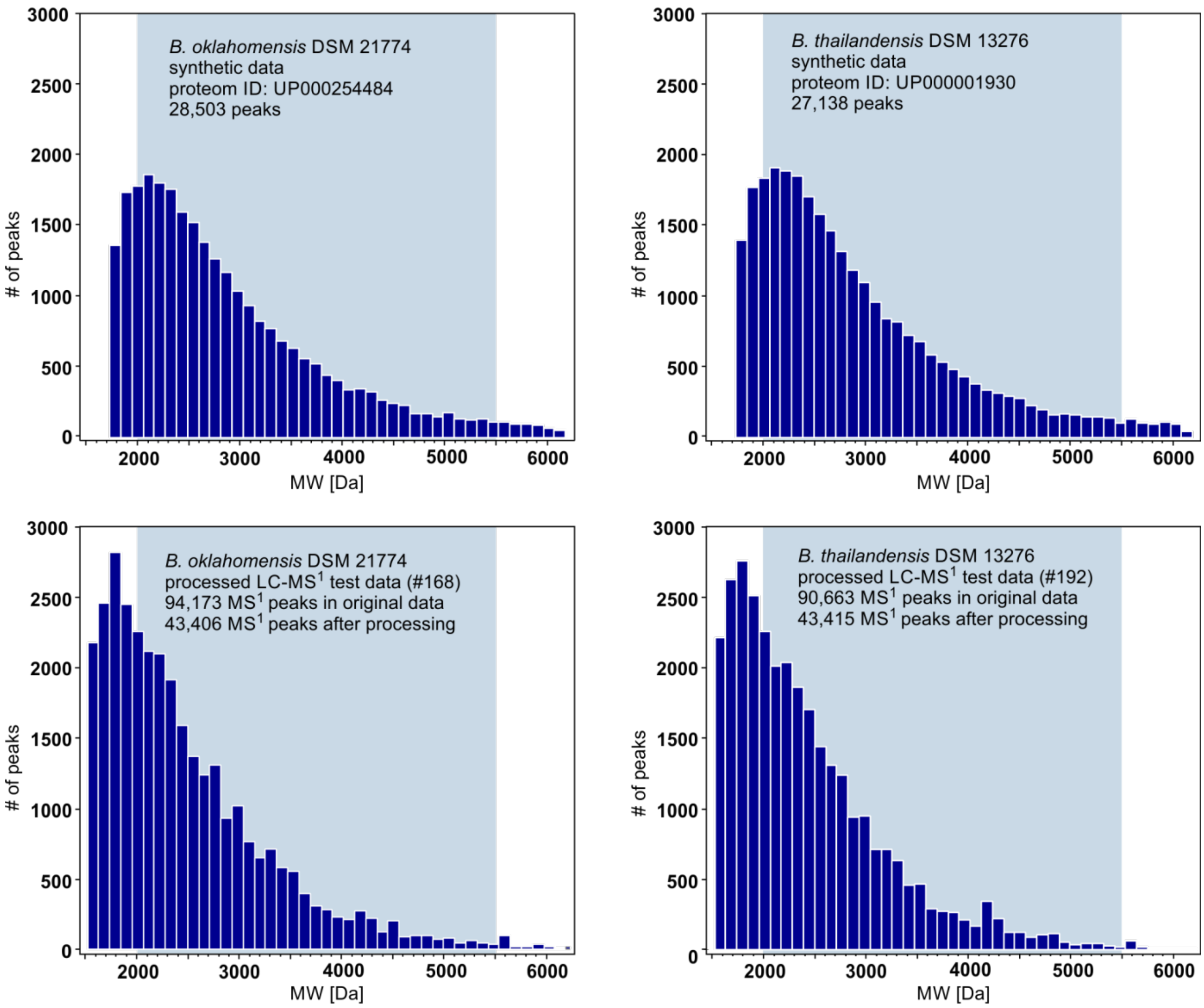


**Figure S2.**

Comparison of density functions of synthetic molecular weight (MW) and LC-MS[1] test data from preparations of *Burkholderia oklahomensis* DSM 21774 (left column) or *Burkholderia thailandensis* DSM 13276 (right).

***Top row:*** peptide MW density functions from strain-specific *in silico* profiles derived from UniProtKB data by the Matlab-based *parseuniprot* toolbox.

***Bottom row:*** peak density functions obtained from processed LC-MS[1] test data; sample preparation has been carried according to the SPEED sample preparation protocol.

Pre-processing and feature selection of original MS[1] spectra has been carried out by the function *readlcmstxtfile*. The blue shaded area indicates the MW range used for identification analysis by MicrobeMS (2000 – 5500 Da).

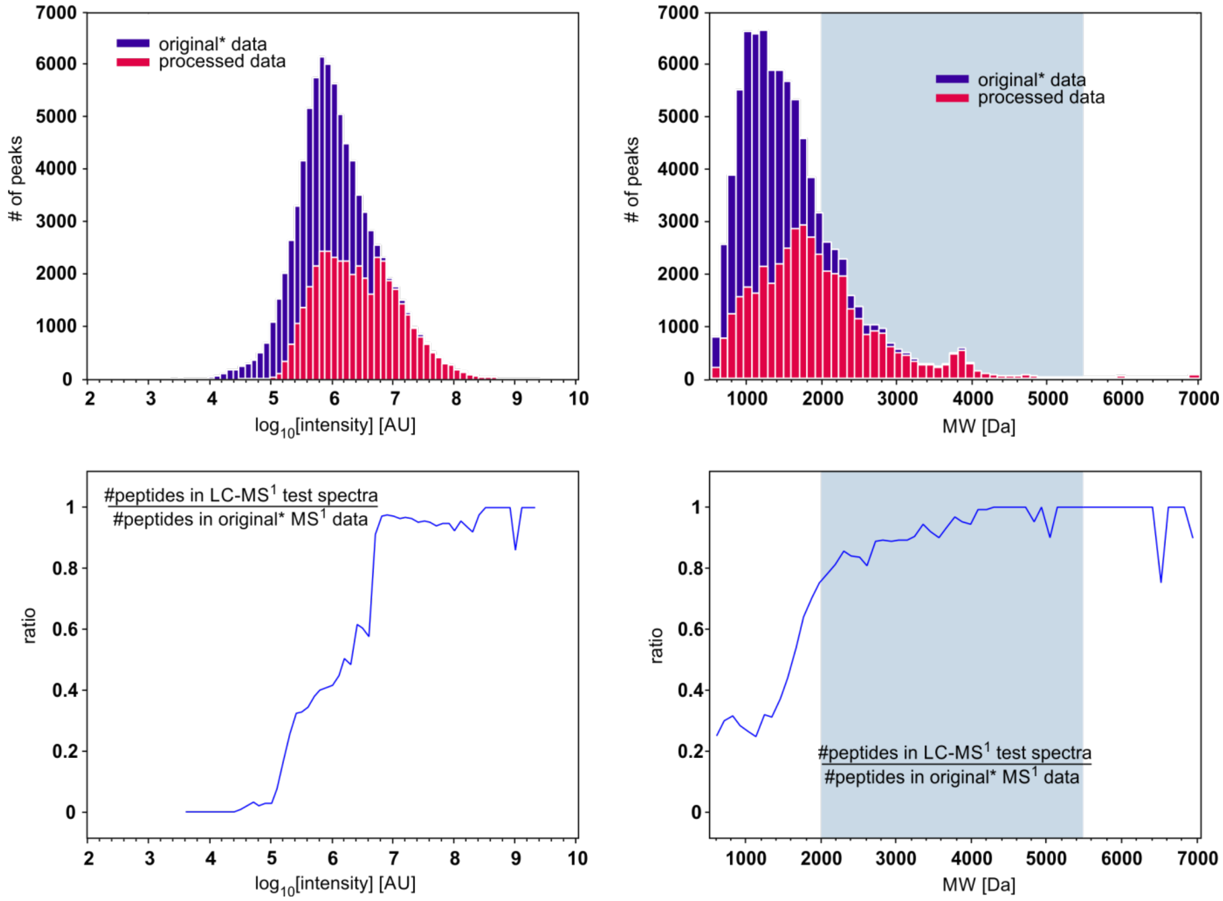


**Figure S3.**

Pre-processing and feature selection of LC-MS[1] data from a preparation of *Mycobacteroides abscessus* DSM 44196

***Top row**:* histogram bar chart of $log_{10}$ scaled MS[1] peak intensities (left) and the molecular weight (MW) distribution (right) of peaks after feature detection by the Minora algorithm (=original* data, blue bars) and after pre-processing and feature selection by the Matlab function *readlcmstxtfile* (processed data, red bars).

Total number of peaks in original / processed data: 85,457 / 43,264

Number of removed peaks from oxidized / deamidated peptides: 483 / 429

***Lower row**:* ratio between the number of peaks present in processed and in original MS[1] data as a function of peak intensity ($log_{10}$ scaled, left) or of the MW (right).

The blue shaded area indicates the MW range used for identification analysis by MicrobeMS (2000 – 5500 Da).

**Figure S4.**

Identification results of the *Burkholderia* subset of LC-MS[1] test data. This subset contained 24 LC-MS[1] test spectra from 5 different strains of *Burkholderia thailandensis* and one strain of *Burkholderia* oklahomensis (cf. samples #5-#10 of table 1). The data were obtained from two independent cultivations of each strain; from the first cultivation series three technical replicate samples per strains were characterized. For processing microbial cells, the SPEED proteomic sample preparation protocol was used. Identification analysis has been carried out by MicrobeMS using the *in silico* database of strain-specific tryptic peptide mass profiles (see text for further details).

### MicrobeMS CLASSIFICATION REPORT FOR MICROORGANISM MASS SPECTRA - OVERVIEW

#### Database search parameters

| | |
|---|---|
| **database name:** | non-redundant-proteomes-22000-processed-Trembl+Swissprot.pkf_90_s_0_time_06_05_2019_10_38_42_red.pkf |
| **date/time of analysis:** | 06-Sep-2019 / 14:12:36 |
| **# of database entries:** | 12044 |
| **distance method:** | Pareto scaling 0.25 |
| **use weightings:** | distances obtained from barcode spectra |
| **vary calibration parameters:** | the complete set of calibration parameters was applied |
| **# of variations:** | 125 ⇒ $(2\times2+1)^3$ variations of calibration parameters |
| **rel width of m/z intervals (ppm):** | 1.2 |
| **calib variation range factor:** | 4 |
| **m/z range, Mmin:** | 1950 |
| **m/z range, Mmax:** | 5125 |
| **peak number ratio corr factor:** | 2 |

#### Short classification report for test spectra

| ID's of test spectra, links to subreports | Best hits in the database | Scores | Log scores |
|---|---|---|---|
| **# 1** - 165_2019_ZBS6_Peter-Burkholderia_E131 | **1** - Burkholderia thailandensis ATCC 700388 / DSM 13276 / CIP 106301 / E264<br>**2** - Burkholderia thailandensis<br>**3** - Burkholderia pseudomallei 1710b | 204.9895<br>196.8191<br>159.7089 | 2.3117<br>2.2941<br>2.2033 |
| **# 2** - 169_2019_ZBS6_Peter-Burkholderia_E131 | **1** - Burkholderia thailandensis ATCC 700388 / DSM 13276 / CIP 106301 / E264<br>**2** - Burkholderia thailandensis<br>**3** - Burkholderia pseudomallei 668 | 201.5313<br>195.0903<br>157.5362 | 2.3043<br>2.2902<br>2.1974 |
| **# 3** - 173_2019_ZBS6_Peter-Burkholderia_E131 | **1** - Burkholderia thailandensis ATCC 700388 / DSM 13276 / CIP 106301 / E264<br>**2** - Burkholderia thailandensis<br>**3** - Burkholderia pseudomallei 1710b | 202.8462<br>195.001<br>156.4678 | 2.3072<br>2.29<br>2.1944 |
| **# 4** - 245_2019_ZBS6_Peter-Burkholderia_E131 | **1** - Burkholderia thailandensis ATCC 700388 / DSM 13276 / CIP 106301 / E264<br>**2** - Burkholderia thailandensis<br>**3** - Burkholderia pseudomallei | 186.3768<br>180.5832<br>151.6297 | 2.2704<br>2.2567<br>2.1808 |
| **# 5** - 166_2019_ZBS6_Peter-Burkholderia_E153 | **1** - Burkholderia thailandensis ATCC 700388 / DSM 13276 / CIP 106301 / E264<br>**2** - Burkholderia thailandensis<br>**3** - Burkholderia pseudomallei 1710b | 214.043<br>204.8531<br>164.4795 | 2.3305<br>2.3114<br>2.2161 |
| **# 6** - 170_2019_ZBS6_Peter-Burkholderia_E153 | **1** - Burkholderia thailandensis ATCC 700388 / DSM 13276 / CIP 106301 / E264<br>**2** - Burkholderia thailandensis<br>**3** - Burkholderia pseudomallei 1710b | 209.5739<br>201.2091<br>162.4502 | 2.3213<br>2.3036<br>2.2107 |
| **# 7** - 174_2019_ZBS6_Peter-Burkholderia_E153 | **1** - Burkholderia thailandensis ATCC 700388 / DSM 13276 / CIP 106301 / E264<br>**2** - Burkholderia thailandensis<br>**3** - Burkholderia pseudomallei | 210.9993<br>201.4565<br>161.5364 | 2.3243<br>2.3042<br>2.2083 |
| **# 8** - 244_2019_ZBS6_Peter-Burkholderia_E153 | **1** - Burkholderia thailandensis ATCC 700388 / DSM 13276 / CIP 106301 / E264<br>**2** - Burkholderia thailandensis<br>**3** - Burkholderia pseudomallei 1710b | 194.7787<br>187.3294<br>154.5136 | 2.2895<br>2.2726<br>2.189 |
| **# 9** - 189_2019_ZBS6_Peter-Burkholderia_E125 | **1** - Burkholderia thailandensis ATCC 700388 / DSM 13276 / CIP 106301 / E264<br>**2** - Burkholderia thailandensis<br>**3** - Burkholderia pseudomallei 1710b | 202.7592<br>195.6465<br>161.2297 | 2.307<br>2.2915<br>2.2074 |
| **# 10** - 190_2019_ZBS6_Peter-Burkholderia_E125 | **1** - Burkholderia thailandensis ATCC 700388 / DSM 13276 / CIP 106301 / E264<br>**2** - Burkholderia thailandensis<br>**3** - Burkholderia pseudomallei 1710b | 198.8781<br>191.8065<br>157.03 | 2.2986<br>2.2829<br>2.196 |
| **# 11** - 191_2019_ZBS6_Peter-Burkholderia_E125 | **1** - Burkholderia thailandensis ATCC 700388 / DSM 13276 / CIP 106301 / E264<br>**2** - Burkholderia thailandensis<br>**3** - Burkholderia pseudomallei 1710b | 202.6297<br>194.8451<br>164.1784 | 2.3067<br>2.2897<br>2.2153 |
| **# 12** - 246_2019_ZBS6_Peter-Burkholderia_E125 | **1** - Burkholderia thailandensis ATCC 700388 / DSM 13276 / CIP 106301 / E264<br>**2** - Burkholderia thailandensis<br>**3** - Burkholderia pseudomallei 1710b | 190.1885<br>185.4785<br>154.1379 | 2.2792<br>2.2683<br>2.1879 |
| **# 13** - 167_2019_ZBS6_Peter-Burkholderia_20219 | **1** - Burkholderia thailandensis ATCC 700388 / DSM 13276 / CIP 106301 / E264<br>**2** - Burkholderia thailandensis<br>**3** - Burkholderia pseudomallei 1710b | 220.4719<br>207.1905<br>167.539 | 2.3434<br>2.3164<br>2.2241 |
| **# 14** - 171_2019_ZBS6_Peter-Burkholderia_20219 | **1** - Burkholderia thailandensis ATCC 700388 / DSM 13276 / CIP 106301 / E264<br>**2** - Burkholderia thailandensis<br>**3** - Burkholderia pseudomallei 1710b | 212.2679<br>199.9794<br>164.3614 | 2.3269<br>2.301<br>2.2158 |
| **# 15** - 175_2019_ZBS6_Peter-Burkholderia_20219 | **1** - Burkholderia thailandensis ATCC 700388 / DSM 13276 / CIP 106301 / E264<br>**2** - Burkholderia thailandensis<br>**3** - Burkholderia pseudomallei 1710b | 221.6617<br>206.7336<br>167.497 | 2.3457<br>2.3154<br>2.224 |
| **# 16** - 247_2019_ZBS6_Peter-Burkholderia_LMG20219 | **1** - Burkholderia thailandensis ATCC 700388 / DSM 13276 / CIP 106301 / E264<br>**2** - Burkholderia thailandensis<br>**3** - Burkholderia pseudomallei 1710b | 203.342<br>192.4561<br>161.0222 | 2.3082<br>2.2843<br>2.2069 |

| | | | |
|---|---|---|---|
| **# 17** - 192_2019_ZBS6_Peter-Burkholderia_DSM13277 | **1** - Burkholderia thailandensis ATCC 700388 / DSM 13276 / CIP 106301 / E264 | 204.233 | 2.3101 |
| | **2** - Burkholderia thailandensis | 194.5373 | 2.289 |
| | **3** - Burkholderia pseudomallei 668 | 159.5722 | 2.203 |
| **# 18** - 193_2019_ZBS6_Peter-Burkholderia_DSM13277 | **1** - Burkholderia thailandensis ATCC 700388 / DSM 13276 / CIP 106301 / E264 | 206.5838 | 2.3151 |
| | **2** - Burkholderia thailandensis | 197.0168 | 2.2945 |
| | **3** - Burkholderia pseudomallei 668 | 162.9761 | 2.2121 |
| **# 19** - 194_2019_ZBS6_Peter-Burkholderia_DSM13277 | **1** - Burkholderia thailandensis ATCC 700388 / DSM 13276 / CIP 106301 / E264 | 208.7549 | 2.3196 |
| | **2** - Burkholderia thailandensis | 196.7919 | 2.294 |
| | **3** - Burkholderia pseudomallei 1710b | 162.134 | 2.2099 |
| **# 20** - 248_2019_ZBS6_Peter-Burkholderia_DSM13277 | **1** - Burkholderia thailandensis ATCC 700388 / DSM 13276 / CIP 106301 / E264 | 192.1228 | 2.2836 |
| | **2** - Burkholderia thailandensis | 183.0681 | 2.2626 |
| | **3** - Burkholderia pseudomallei Pseudomonas pseudomallei | 153.6698 | 2.1866 |
| **# 21** - 168_2019_ZBS6_Peter-Burkholderia_21774 | **1** - Burkholderia oklahomensis | 217.199 | 2.3369 |
| | **2** - Burkholderia pseudomallei | 149.3277 | 2.1741 |
| | **3** - Burkholderia pseudomallei 1710b | 149.2699 | 2.174 |
| **# 22** - 172_2019_ZBS6_Peter-Burkholderia_21774 | **1** - Burkholderia oklahomensis | 211.9611 | 2.3263 |
| | **2** - Burkholderia pseudomallei 1710b | 146.1807 | 2.1649 |
| | **3** - Burkholderia pseudomallei | 145.7201 | 2.1635 |
| **# 23** - 176_2019_ZBS6_Peter-Burkholderia_21774 | **1** - Burkholderia oklahomensis | 207.9029 | 2.3179 |
| | **2** - Burkholderia pseudomallei | 147.1922 | 2.1679 |
| | **3** - Burkholderia pseudomallei 1710b | 147.169 | 2.1678 |
| **# 24** - 243_2019_ZBS6_Peter-Burkholderia_DSM21774 | **1** - Burkholderia oklahomensis | 202.1129 | 2.3056 |
| | **2** - Burkholderia pseudomallei 1710b | 148.8684 | 2.1728 |
| | **3** - Burkholderia pseudomallei 668 | 147.8435 | 2.1698 |

---

#### # 1 - CLASSIFICATION SUBREPORT FOR SPECTRUM *'165_2019_ZBS6_Peter-Burkholderia_E131'*

##### Actual test mass spectrum

| | |
|---|---|
| **genus / species / strain:** | Burkholderia thailandensis E131 |
| **file id:** | 165_2019_ZBS6_Peter-Burkholderia_E131 |
| **type:** | 165 |
| **NCBI id (species level):** | NaN |
| **NCBI id (strain level):** | NaN |
| **growth time:** | 24 h |
| **growth temperature:** | 37Â°C |
| **growth conditions:** | no CO2 |
| **growth medium:** | Caso (Oxo d) |
| **sample treatment:** | SPEED, operator: A. Schneider |
| **spores:** | no |
| **concentration:** | not available |
| **extra info:** | biological replicate #1, technical replicate #1 |
| **calibration standard:** | not available |
| **measurement method:** | LC-MS1 spectrum, method not available |
| **customer:** | RKI/ZBS 6 |
| **measurement date/time:** | 2019-01_11:11:32.906+02:00 |
| **path to MS file:** | E:\Matlab |

##### List of best matches with test spectrum

| No | Genus/Species/Strain | Score | Log Score | Peak Numbers | Spectral ID | Proteome ID |
|---|---|---|---|---|---|---|
| 1 | Burkholderia thailandensis ATCC 700388 / DSM 13276 / CIP 106301 / E264 | 204.9895 | 2.3117 | 16465 / 23226 | dbspec-15-Mar-2019-3-2-18.411 | UP000001930 |
| 2 | Burkholderia thailandensis | 196.8191 | 2.2941 | 16465 / 22799 | dbspec-15-Mar-2019-3-9-35.914 | UP000235972 |
| 3 | Burkholderia pseudomallei 1710b | 159.7089 | 2.2033 | 16465 / 24945 | dbspec-15-Mar-2019-3-2-33.574 | UP000002700 |
| 4 | Burkholderia pseudomallei | 159.0635 | 2.2016 | 16465 / 24544 | dbspec-15-Mar-2019-3-2-15.743 | UP000001812 |
| 5 | Burkholderia pseudomallei | 158.8731 | 2.2011 | 16465 / 24832 | dbspec-15-Mar-2019-3-5-56.499 | UP000030449 |
| 6 | Burkholderia pseudomallei | 158.737 | 2.2007 | 16465 / 24836 | dbspec-15-Mar-2019-3-2-21.734 | UP000002031 |
| 7 | Burkholderia pseudomallei 668 | 158.4655 | 2.1999 | 16465 / 24129 | dbspec-15-Mar-2019-3-2-23.122 | UP000002153 |
| 8 | Burkholderia pseudomallei 1106a | 158.192 | 2.1992 | 16465 / 24105 | dbspec-15-Mar-2019-3-3-42.854 | UP000006738 |
| 9 | Burkholderia pseudomallei Pseudomonas pseudomallei | 158.0812 | 2.1989 | 16465 / 24887 | dbspec-15-Mar-2019-3-5-52.272 | UP000029527 |
| 10 | Burkholderia pseudomallei | 157.8695 | 2.1983 | 16465 / 24032 | dbspec-15-Mar-2019-3-3-23.479 | UP000005700 |
| 11 | Burkholderia pseudomallei | 157.687 | 2.1978 | 16465 / 24475 | dbspec-15-Mar-2019-3-3-10.094 | UP000004875 |
| 12 | Burkholderia pseudomallei | 157.6827 | 2.1978 | 16465 / 24630 | dbspec-15-Mar-2019-3-6-4.424 | UP000032603 |
| 13 | Burkholderia pseudomallei K96243 | 157.6149 | 2.1976 | 16465 / 23961 | dbspec-15-Mar-2019-3-1-58.302 | UP000000605 |
| 14 | Burkholderia pseudomallei 1026b | 157.4712 | 2.1972 | 16465 / 23916 | dbspec-15-Mar-2019-3-4-15.005 | UP000010087 |
| 15 | Burkholderia pseudomallei | 156.1092 | 2.1934 | 16465 / 23842 | dbspec-15-Mar-2019-3-3-0.858 | UP000004448 |
| 16 | Burkholderia thailandensis | 155.1077 | 2.1906 | 16465 / 23446 | dbspec-15-Mar-2019-3-8-58.458 | UP000218714 |
| 17 | Burkholderia pseudomallei | 154.7006 | 2.1895 | 16465 / 22928 | dbspec-15-Mar-2019-3-2-35.82 | UP000002781 |
| 18 | Burkholderia pseudomallei | 152.2044 | 2.1824 | 16465 / 21894 | dbspec-15-Mar-2019-3-3-7.02 | UP000004700 |
| 19 | Burkholderia mallei Pseudomonas mallei | 150.8386 | 2.1785 | 16465 / 19381 | dbspec-15-Mar-2019-3-10-40.934 | UP000254505 |
| 20 | Burkholderia mallei | 149.5895 | 2.1749 | 16465 / 19432 | dbspec-15-Mar-2019-3-2-54.15 | UP000003995 |
| 21 | Burkholderia mallei ATCC 23344 | 149.1502 | 2.1736 | 16465 / 18442 | dbspec-15-Mar-2019-3-3-41.84 | UP000006693 |
| 22 | Burkholderia mallei NCTC 10229 | 147.7165 | 2.1694 | 16465 / 18362 | dbspec-15-Mar-2019-3-2-26.164 | UP000002283 |
| 23 | Burkholderia oklahomensis | 146.3237 | 2.1653 | 16465 / 24424 | dbspec-15-Mar-2019-3-10-40.31 | UP000254484 |

| | | | | | | |
|---|---|---|---|---|---|---|
| 24 | Hel cobacter pylori | 134.6405 | 2.1292 | 16465 / 6078 | dbspec-15-Mar-2019-3-2-38.067 | UP000002997 |
| 25 | Hel cobacter pylori | 134.331 | 2.1282 | 16465 / 6107 | dbspec-15-Mar-2019-3-2-56.974 | UP000004225 |
| 26 | Porphyromonas macacae | 134.0234 | 2.1272 | 16465 / 8458 | dbspec-15-Mar-2019-3-5-55.329 | UP000030103 |
| 27 | Porphyromonas macacae | 133.9486 | 2.1269 | 16465 / 7416 | dbspec-15-Mar-2019-3-10-38.392 | UP000254156 |
| 28 | Porphyromonas macacae | 133.2957 | 2.1248 | 16465 / 8196 | dbspec-15-Mar-2019-3-10-38.782 | UP000254263 |
| 29 | Streptobacillus moniliformis ATCC 14647 / DSM 12112 / NCTC 10651 / 9901 | 133.0254 | 2.1239 | 16465 / 5488 | dbspec-15-Mar-2019-3-2-22.685 | UP000002072 |
| 30 | Hel cobacter pylori | 132.4187 | 2.1219 | 16465 / 6070 | dbspec-15-Mar-2019-3-3-43.306 | UP000006748 |

---

#### # 2 - CLASSIFICATION SUBREPORT FOR SPECTRUM *'169_2019_ZBS6_Peter-Burkholderia_E131'*

##### Actual test mass spectrum

| | |
|---|---|
| **genus / species / strain:** | Burkholderia thailandensis E131 |
| **file id:** | 169_2019_ZBS6_Peter-Burkholderia_E131 |
| **type:** | 169 |
| **NCBI id (species level):** | NaN |
| **NCBI id (strain level):** | NaN |
| **growth time:** | 24 h |
| **growth temperature:** | 37Â°C |
| **growth conditions:** | no CO2 |
| **growth medium:** | Caso (Oxo d) |
| **sample treatment:** | SPEED, operator: A. Schneider |
| **spores:** | no |
| **concentration:** | not available |
| **extra info:** | biological replicate #1, technical replicate #2 |
| **calibration standard:** | not available |
| **measurement method:** | LC-MS1 spectrum, method not available |
| **customer:** | RKI/ZBS 6 |
| **measurement date/time:** | 2019-01_11:11:32.906+02:00 |
| **path to MS file:** | E:\Matlab |

##### List of best matches with test spectrum

| No | Genus/Species/Strain | Score | Log Score | Peak Numbers | Spectral ID | Proteome ID |
|---|---|---|---|---|---|---|
| 1 | Burkholderia thailandensis ATCC 700388 / DSM 13276 / CIP 106301 / E264 | 201.5313 | 2.3043 | 16417 / 23226 | dbspec-15-Mar-2019-3-2-18.411 | UP000001930 |
| 2 | Burkholderia thailandensis | 195.0903 | 2.2902 | 16417 / 22799 | dbspec-15-Mar-2019-3-9-35.914 | UP000235972 |
| 3 | Burkholderia pseudomallei 668 | 157.5362 | 2.1974 | 16417 / 24129 | dbspec-15-Mar-2019-3-2-23.122 | UP000002153 |
| 4 | Burkholderia pseudomallei 1710b | 157.4994 | 2.1973 | 16417 / 24945 | dbspec-15-Mar-2019-3-2-33.574 | UP000002700 |
| 5 | Burkholderia pseudomallei 1106a | 156.6792 | 2.195 | 16417 / 24105 | dbspec-15-Mar-2019-3-3-42.854 | UP000006738 |
| 6 | Burkholderia pseudomallei | 156.1648 | 2.1936 | 16417 / 24544 | dbspec-15-Mar-2019-3-2-15.743 | UP000001812 |
| 7 | Burkholderia pseudomallei | 156.1151 | 2.1934 | 16417 / 24836 | dbspec-15-Mar-2019-3-2-21.734 | UP000002031 |
| 8 | Burkholderia pseudomallei | 155.9862 | 2.1931 | 16417 / 24032 | dbspec-15-Mar-2019-3-3-23.479 | UP000005700 |
| 9 | Burkholderia pseudomallei | 155.9696 | 2.193 | 16417 / 24630 | dbspec-15-Mar-2019-3-6-4.424 | UP000032603 |
| 10 | Burkholderia pseudomallei Pseudomonas pseudomallei | 155.4675 | 2.1916 | 16417 / 24887 | dbspec-15-Mar-2019-3-5-52.272 | UP000029527 |
| 11 | Burkholderia pseudomallei | 155.3294 | 2.1913 | 16417 / 24832 | dbspec-15-Mar-2019-3-5-56.499 | UP000030449 |
| 12 | Burkholderia pseudomallei K96243 | 155.2974 | 2.1912 | 16417 / 23961 | dbspec-15-Mar-2019-3-1-58.302 | UP000000605 |
| 13 | Burkholderia pseudomallei | 155.065 | 2.1905 | 16417 / 24475 | dbspec-15-Mar-2019-3-3-10.094 | UP000004875 |
| 14 | Burkholderia pseudomallei 1026b | 154.5238 | 2.189 | 16417 / 23916 | dbspec-15-Mar-2019-3-4-15.005 | UP000010087 |
| 15 | Burkholderia pseudomallei | 154.5231 | 2.189 | 16417 / 23842 | dbspec-15-Mar-2019-3-3-0.858 | UP000004448 |
| 16 | Burkholderia pseudomallei | 152.4687 | 2.1832 | 16417 / 22928 | dbspec-15-Mar-2019-3-2-35.82 | UP000002781 |
| 17 | Burkholderia thailandensis | 151.5999 | 2.1807 | 16417 / 23446 | dbspec-15-Mar-2019-3-8-58.458 | UP000218714 |
| 18 | Burkholderia pseudomallei | 150.126 | 2.1765 | 16417 / 21894 | dbspec-15-Mar-2019-3-3-7.02 | UP000004700 |
| 19 | Burkholderia mallei Pseudomonas mallei | 148.5723 | 2.1719 | 16417 / 19381 | dbspec-15-Mar-2019-3-10-40.934 | UP000254505 |
| 20 | Burkholderia mallei | 147.7319 | 2.1695 | 16417 / 19432 | dbspec-15-Mar-2019-3-2-54.15 | UP000003995 |
| 21 | Burkholderia mallei ATCC 23344 | 146.9034 | 2.167 | 16417 / 18442 | dbspec-15-Mar-2019-3-3-41.84 | UP000006693 |
| 22 | Burkholderia oklahomensis | 146.0295 | 2.1644 | 16417 / 24424 | dbspec-15-Mar-2019-3-10-40.31 | UP000254484 |
| 23 | Burkholderia mallei NCTC 10229 | 145.9025 | 2.1641 | 16417 / 18362 | dbspec-15-Mar-2019-3-2-26.164 | UP000002283 |
| 24 | Hel cobacter pylori | 133.9445 | 2.1269 | 16417 / 5873 | dbspec-15-Mar-2019-3-4-51.104 | UP000015602 |
| 25 | Hel cobacter pylori | 133.9101 | 2.1268 | 16417 / 6083 | dbspec-15-Mar-2019-3-3-11.919 | UP000004974 |
| 26 | Porphyromonas macacae | 133.2653 | 2.1247 | 16417 / 8196 | dbspec-15-Mar-2019-3-10-38.782 | UP000254263 |
| 27 | Hel cobacter pylori | 133.1289 | 2.1243 | 16417 / 6100 | dbspec-15-Mar-2019-3-2-42.185 | UP000003215 |
| 28 | Hel cobacter cetorum ATCC BAA-540 / MIT 99-5656 | 132.8923 | 2.1235 | 16417 / 6477 | dbspec-15-Mar-2019-3-3-12.153 | UP000005013 |
| 29 | Hel cobacter pylori | 132.1804 | 2.1212 | 16417 / 5871 | dbspec-15-Mar-2019-3-2-56.116 | UP000004153 |
| 30 | Hel cobacter pylori | 132.0928 | 2.1209 | 16417 / 6070 | dbspec-15-Mar-2019-3-3-43.306 | UP000006748 |

---

#### # 3 - CLASSIFICATION SUBREPORT FOR SPECTRUM *'173_2019_ZBS6_Peter-Burkholderia_E131'*

##### Actual test mass spectrum

| | |
|---|---|
| **genus / species / strain:** | Burkholderia thailandensis E131 |
| **file id:** | 173_2019_ZBS6_Peter-Burkholderia_E131 |
| **type:** | 173 |
| **NCBI id (species level):** | NaN |
| **NCBI id (strain level):** | NaN |

| | |
|---|---|
| **growth time:** | 24 h |
| **growth temperature:** | 37Â°C |
| **growth conditions:** | no CO2 |
| **growth medium:** | Caso (Oxo d) |
| **sample treatment:** | SPEED, operator: A. Schneider |
| **spores:** | no |
| **concentration:** | not available |
| **extra info:** | biological replicate #1, technical replicate #3 |
| **calibration standard:** | not available |
| **measurement method:** | LC-MS1 spectrum, method not available |
| **customer:** | RKI/ZBS 6 |
| **measurement date/time:** | 2019-01_11:11:32.906+02:00 |
| **path to MS file:** | E:\Matlab |

#### List of best matches with test spectrum

| No | Genus/Species/Strain | Score | Log Score | Peak Numbers | Spectral ID | Proteome ID |
|---|---|---|---|---|---|---|
| 1 | Burkholderia thailandensis ATCC 700388 / DSM 13276 / CIP 106301 / E264 | 202.8462 | 2.3072 | 16475 / 23226 | dbspec-15-Mar-2019-3-2-18.411 | UP000001930 |
| 2 | Burkholderia thailandensis | 195.001 | 2.29 | 16475 / 22799 | dbspec-15-Mar-2019-3-9-35.914 | UP000235972 |
| 3 | Burkholderia pseudomallei 1710b | 156.4678 | 2.1944 | 16475 / 24945 | dbspec-15-Mar-2019-3-2-33.574 | UP000002700 |
| 4 | Burkholderia pseudomallei | 156.4422 | 2.1944 | 16475 / 24630 | dbspec-15-Mar-2019-3-6-4.424 | UP000032603 |
| 5 | Burkholderia pseudomallei | 156.3864 | 2.1942 | 16475 / 24544 | dbspec-15-Mar-2019-3-2-15.743 | UP000001812 |
| 6 | Burkholderia pseudomallei 668 | 156.0589 | 2.1933 | 16475 / 24129 | dbspec-15-Mar-2019-3-2-23.122 | UP000002153 |
| 7 | Burkholderia pseudomallei | 155.8169 | 2.1926 | 16475 / 24836 | dbspec-15-Mar-2019-3-2-21.734 | UP000002031 |
| 8 | Burkholderia pseudomallei | 155.7595 | 2.1925 | 16475 / 24832 | dbspec-15-Mar-2019-3-5-56.499 | UP000030449 |
| 9 | Burkholderia pseudomallei | 154.9537 | 2.1902 | 16475 / 24475 | dbspec-15-Mar-2019-3-3-10.094 | UP000004875 |
| 10 | Burkholderia pseudomallei K96243 | 154.9487 | 2.1902 | 16475 / 23961 | dbspec-15-Mar-2019-3-1-58.302 | UP000000605 |
| 11 | Burkholderia pseudomallei 1106a | 154.7047 | 2.1895 | 16475 / 24105 | dbspec-15-Mar-2019-3-3-42.854 | UP000006738 |
| 12 | Burkholderia pseudomallei | 154.3294 | 2.1884 | 16475 / 24032 | dbspec-15-Mar-2019-3-3-23.479 | UP000005700 |
| 13 | Burkholderia pseudomallei 1026b | 154.2357 | 2.1882 | 16475 / 23916 | dbspec-15-Mar-2019-3-4-15.005 | UP000010087 |
| 14 | Burkholderia pseudomallei Pseudomonas pseudomallei | 153.7438 | 2.1868 | 16475 / 24887 | dbspec-15-Mar-2019-3-5-52.272 | UP000029527 |
| 15 | Burkholderia pseudomallei | 152.9291 | 2.1845 | 16475 / 23842 | dbspec-15-Mar-2019-3-3-0.858 | UP000004448 |
| 16 | Burkholderia pseudomallei | 151.5465 | 2.1805 | 16475 / 22928 | dbspec-15-Mar-2019-3-2-35.82 | UP000002781 |
| 17 | Burkholderia thailandensis | 149.9965 | 2.1761 | 16475 / 23446 | dbspec-15-Mar-2019-3-8-58.458 | UP000218714 |
| 18 | Burkholderia pseudomallei | 149.342 | 2.1742 | 16475 / 21894 | dbspec-15-Mar-2019-3-3-7.02 | UP000004700 |
| 19 | Burkholderia mallei Pseudomonas mallei | 147.1693 | 2.1678 | 16475 / 19381 | dbspec-15-Mar-2019-3-10-40.934 | UP000254505 |
| 20 | Burkholderia mallei | 146.4545 | 2.1657 | 16475 / 19432 | dbspec-15-Mar-2019-3-2-54.15 | UP000003995 |
| 21 | Burkholderia mallei ATCC 23344 | 146.2168 | 2.165 | 16475 / 18442 | dbspec-15-Mar-2019-3-3-41.84 | UP000006693 |
| 22 | Burkholderia mallei NCTC 10229 | 144.5904 | 2.1601 | 16475 / 18362 | dbspec-15-Mar-2019-3-2-26.164 | UP000002283 |
| 23 | Burkholderia oklahomensis | 142.8176 | 2.1548 | 16475 / 24424 | dbspec-15-Mar-2019-3-10-40.31 | UP000254484 |
| 24 | Hel cobacter pylori | 133.5188 | 2.1255 | 16475 / 5871 | dbspec-15-Mar-2019-3-2-56.116 | UP000004153 |
| 25 | Hel cobacter pylori ATCC 700392 / 26695 | 132.8919 | 2.1235 | 16475 / 6120 | dbspec-15-Mar-2019-3-1-54.87 | UP000000429 |
| 26 | Hel cobacter pylori | 132.6766 | 2.1228 | 16475 / 6170 | dbspec-15-Mar-2019-3-2-56.241 | UP000004177 |
| 27 | Hel cobacter pylori | 132.6362 | 2.1227 | 16475 / 6245 | dbspec-15-Mar-2019-3-4-25.208 | UP000011870 |
| 28 | Riemerella anatipestifer | 132.3355 | 2.1217 | 16475 / 8892 | dbspec-15-Mar-2019-3-3-33.868 | UP000006276 |
| 29 | Hel cobacter pylori | 132.2575 | 2.1214 | 16475 / 6310 | dbspec-15-Mar-2019-3-4-27.142 | UP000012012 |
| 30 | Hel cobacter pylori | 132.202 | 2.1212 | 16475 / 6070 | dbspec-15-Mar-2019-3-3-43.306 | UP000006748 |

---

### # 4 - CLASSIFICATION SUBREPORT FOR SPECTRUM *'245_2019_ZBS6_Peter-Burkholderia_E131'*

#### Actual test mass spectrum

| | |
|---|---|
| **genus / species / strain:** | Burkholderia thailandensis E131 |
| **file id:** | 245_2019_ZBS6_Peter-Burkholderia_E131 |
| **type:** | 245 |
| **NCBI id (species level):** | NaN |
| **NCBI id (strain level):** | NaN |
| **growth time:** | 24 h |
| **growth temperature:** | 37Â°C |
| **growth conditions:** | no CO2 |
| **growth medium:** | Caso (Oxo d) |
| **sample treatment:** | SPEED, operator: A. Schneider |
| **spores:** | no |
| **concentration:** | not available |
| **extra info:** | biological replicate #2, technical replicate #1 |
| **calibration standard:** | not available |
| **measurement method:** | LC-MS1 spectrum, method not available |
| **customer:** | RKI/ZBS 6 |
| **measurement date/time:** | 2019-01_11:11:32.906+02:00 |
| **path to MS file:** | E:\Matlab |

#### List of best matches with test spectrum

| No | Genus/Species/Strain | Score | Log Score | Peak Numbers | Spectral ID | Proteome ID |
|---|---|---|---|---|---|---|
| 1 | Burkholderia thailandensis ATCC 700388 / DSM 13276 / CIP 106301 / E264 | 186.3768 | 2.2704 | 15032 / 23226 | dbspec-15-Mar-2019-3-2-18.411 | UP000001930 |

| | | | | | | |
|---|---|---|---|---|---|---|
| 2 | Burkholderia thailandensis | 180.5832 | 2.2567 | 15032 / 22799 | dbspec-15-Mar-2019-3-9-35.914 | UP000235972 |
| 3 | Burkholderia pseudomallei | 151.6297 | 2.1808 | 15032 / 24630 | dbspec-15-Mar-2019-3-6-4.424 | UP000032603 |
| 4 | Burkholderia pseudomallei 668 | 151.5052 | 2.1804 | 15032 / 24129 | dbspec-15-Mar-2019-3-2-23.122 | UP000002153 |
| 5 | Burkholderia pseudomallei 1710b | 151.3751 | 2.1801 | 15032 / 24945 | dbspec-15-Mar-2019-3-2-33.574 | UP000002700 |
| 6 | Burkholderia pseudomallei | 151.0921 | 2.1792 | 15032 / 24032 | dbspec-15-Mar-2019-3-3-23.479 | UP000005700 |
| 7 | Burkholderia pseudomallei | 150.9367 | 2.1788 | 15032 / 24475 | dbspec-15-Mar-2019-3-3-10.094 | UP000004875 |
| 8 | Burkholderia pseudomallei | 150.3872 | 2.1772 | 15032 / 24544 | dbspec-15-Mar-2019-3-2-15.743 | UP000001812 |
| 9 | Burkholderia pseudomallei Pseudomonas pseudomallei | 150.3303 | 2.177 | 15032 / 24887 | dbspec-15-Mar-2019-3-5-52.272 | UP000029527 |
| 10 | Burkholderia pseudomallei K96243 | 150.2091 | 2.1767 | 15032 / 23961 | dbspec-15-Mar-2019-3-1-58.302 | UP000000605 |
| 11 | Burkholderia pseudomallei | 149.9692 | 2.176 | 15032 / 24836 | dbspec-15-Mar-2019-3-2-21.734 | UP000002031 |
| 12 | Burkholderia pseudomallei 1106a | 149.9354 | 2.1759 | 15032 / 24105 | dbspec-15-Mar-2019-3-3-42.854 | UP000006738 |
| 13 | Burkholderia pseudomallei 1026b | 149.4664 | 2.1745 | 15032 / 23916 | dbspec-15-Mar-2019-3-4-15.005 | UP000010087 |
| 14 | Burkholderia pseudomallei | 149.3842 | 2.1743 | 15032 / 23842 | dbspec-15-Mar-2019-3-3-0.858 | UP000004448 |
| 15 | Burkholderia pseudomallei | 148.9053 | 2.1729 | 15032 / 24832 | dbspec-15-Mar-2019-3-5-56.499 | UP000030449 |
| 16 | Burkholderia pseudomallei | 146.9434 | 2.1671 | 15032 / 22928 | dbspec-15-Mar-2019-3-2-35.82 | UP000002781 |
| 17 | Burkholderia thailandensis | 146.4459 | 2.1657 | 15032 / 23446 | dbspec-15-Mar-2019-3-8-58.458 | UP000218714 |
| 18 | Burkholderia pseudomallei | 145.1343 | 2.1618 | 15032 / 21894 | dbspec-15-Mar-2019-3-3-7.02 | UP000004700 |
| 19 | Burkholderia mallei | 143.3865 | 2.1565 | 15032 / 19432 | dbspec-15-Mar-2019-3-2-54.15 | UP000003995 |
| 20 | Burkholderia mallei Pseudomonas mallei | 142.5067 | 2.1538 | 15032 / 19381 | dbspec-15-Mar-2019-3-10-40.934 | UP000254505 |
| 21 | Burkholderia mallei NCTC 10229 | 140.3604 | 2.1472 | 15032 / 18362 | dbspec-15-Mar-2019-3-2-26.164 | UP000002283 |
| 22 | Burkholderia mallei ATCC 23344 | 139.723 | 2.1453 | 15032 / 18442 | dbspec-15-Mar-2019-3-3-41.84 | UP000006693 |
| 23 | Burkholderia oklahomensis | 138.3041 | 2.1408 | 15032 / 24424 | dbspec-15-Mar-2019-3-10-40.31 | UP000254484 |
| 24 | Hel cobacter pylori | 131.6856 | 2.1195 | 15032 / 5968 | dbspec-15-Mar-2019-3-3-19.251 | UP000005411 |
| 25 | Burkholderia vietnamiensis G4 / LMG 22486 | 131.3273 | 2.1184 | 15032 / 27924 | dbspec-15-Mar-2019-3-2-26.18 | UP000002287 |
| 26 | Burkholderia pseudomultivorans | 130.6877 | 2.1162 | 15032 / 27165 | dbspec-15-Mar-2019-3-6-36.232 | UP000061512 |
| 27 | Hel cobacter pylori | 130.6545 | 2.1161 | 15032 / 6078 | dbspec-15-Mar-2019-3-2-38.067 | UP000002997 |
| 28 | Burkholderia cenocepacia | 130.4963 | 2.1156 | 15032 / 28696 | dbspec-15-Mar-2019-3-5-52.022 | UP000029413 |
| 29 | Leptotr chia wadei F0279 | 130.4006 | 2.1153 | 15032 / 7438 | dbspec-15-Mar-2019-3-4-56.907 | UP000016626 |
| 30 | Burkholderia puraquae | 130.1786 | 2.1145 | 15032 / 28790 | dbspec-15-Mar-2019-3-8-4.997 | UP000193146 |

#### # 5 - CLASSIFICATION SUBREPORT FOR SPECTRUM *'166_2019_ZBS6_Peter-Burkholderia_E153'*

##### Actual test mass spectrum

| | |
|---|---|
| **genus / species / strain:** | Burkholderia thailandensis E153 |
| **file id:** | 166_2019_ZBS6_Peter-Burkholderia_E153 |
| **type:** | 166 |
| **NCBI id (species level):** | NaN |
| **NCBI id (strain level):** | NaN |
| **growth time:** | 24 h |
| **growth temperature:** | 37Â°C |
| **growth conditions:** | no CO2 |
| **growth medium:** | Caso (Oxo d) |
| **sample treatment:** | SPEED, operator: A. Schneider |
| **spores:** | no |
| **concentration:** | not available |
| **extra info:** | biological replicate #1, technical replicate #1 |
| **calibration standard:** | not available |
| **measurement method:** | LC-MS1 spectrum, method not available |
| **customer:** | RKI/ZBS 6 |
| **measurement date/time:** | 2019-01_11:11:32.906+02:00 |
| **path to MS file:** | E:\Matlab |

##### List of best matches with test spectrum

| No | Genus/Species/Strain | Score | Log Score | Peak Numbers | Spectral ID | Proteome ID |
|---|---|---|---|---|---|---|
| 1 | Burkholderia thailandensis ATCC 700388 / DSM 13276 / CIP 106301 / E264 | 214.043 | 2.3305 | 16640 / 23226 | dbspec-15-Mar-2019-3-2-18.411 | UP000001930 |
| 2 | Burkholderia thailandensis | 204.8531 | 2.3114 | 16640 / 22799 | dbspec-15-Mar-2019-3-9-35.914 | UP000235972 |
| 3 | Burkholderia pseudomallei 1710b | 164.4795 | 2.2161 | 16640 / 24945 | dbspec-15-Mar-2019-3-2-33.574 | UP000002700 |
| 4 | Burkholderia pseudomallei | 163.3285 | 2.2131 | 16640 / 24544 | dbspec-15-Mar-2019-3-2-15.743 | UP000001812 |
| 5 | Burkholderia pseudomallei | 162.8408 | 2.2118 | 16640 / 24630 | dbspec-15-Mar-2019-3-6-4.424 | UP000032603 |
| 6 | Burkholderia pseudomallei | 162.8326 | 2.2117 | 16640 / 24032 | dbspec-15-Mar-2019-3-3-23.479 | UP000005700 |
| 7 | Burkholderia pseudomallei | 162.2659 | 2.2102 | 16640 / 24832 | dbspec-15-Mar-2019-3-5-56.499 | UP000030449 |
| 8 | Burkholderia pseudomallei | 162.1738 | 2.21 | 16640 / 24836 | dbspec-15-Mar-2019-3-2-21.734 | UP000002031 |
| 9 | Burkholderia pseudomallei 1106a | 162.0457 | 2.2096 | 16640 / 24105 | dbspec-15-Mar-2019-3-3-42.854 | UP000006738 |
| 10 | Burkholderia pseudomallei 668 | 161.8398 | 2.2091 | 16640 / 24129 | dbspec-15-Mar-2019-3-2-23.122 | UP000002153 |
| 11 | Burkholderia pseudomallei 1026b | 161.6791 | 2.2087 | 16640 / 23916 | dbspec-15-Mar-2019-3-4-15.005 | UP000010087 |
| 12 | Burkholderia pseudomallei K96243 | 161.5111 | 2.2082 | 16640 / 23961 | dbspec-15-Mar-2019-3-1-58.302 | UP000000605 |
| 13 | Burkholderia pseudomallei Pseudomonas pseudomallei | 161.1092 | 2.2071 | 16640 / 24887 | dbspec-15-Mar-2019-3-5-52.272 | UP000029527 |
| 14 | Burkholderia pseudomallei | 160.8697 | 2.2065 | 16640 / 23842 | dbspec-15-Mar-2019-3-3-0.858 | UP000004448 |
| 15 | Burkholderia pseudomallei | 160.8405 | 2.2064 | 16640 / 24475 | dbspec-15-Mar-2019-3-3-10.094 | UP000004875 |
| 16 | Burkholderia thailandensis | 159.8711 | 2.2038 | 16640 / 23446 | dbspec-15-Mar-2019-3-8-58.458 | UP000218714 |
| 17 | Burkholderia pseudomallei | 159.0081 | 2.2014 | 16640 / 22928 | dbspec-15-Mar-2019-3-2-35.82 | UP000002781 |
| 18 | Burkholderia pseudomallei | 156.7144 | 2.1951 | 16640 / 21894 | dbspec-15-Mar-2019-3-3-7.02 | UP000004700 |
| 19 | Burkholderia mallei Pseudomonas mallei | 152.5469 | 2.1834 | 16640 / 19381 | dbspec-15-Mar-2019-3-10-40.934 | UP000254505 |
| 20 | Burkholderia mallei | 151.8871 | 2.1815 | 16640 / 19432 | dbspec-15-Mar-2019-3-2-54.15 | UP000003995 |
| 21 | Burkholderia oklahomensis | 151.0042 | 2.179 | 16640 / 24424 | dbspec-15-Mar-2019-3-10-40.31 | UP000254484 |
| 22 | Burkholderia mallei NCTC 10229 | 150.4921 | 2.1775 | 16640 / 18362 | dbspec-15-Mar-2019-3-2-26.164 | UP000002283 |
| 23 | Burkholderia mallei ATCC 23344 | 150.3305 | 2.177 | 16640 / 18442 | dbspec-15-Mar-2019-3-3-41.84 | UP000006693 |

| | | | | | | |
|---|---|---|---|---|---|---|
| 24 | Riemerella anatipestifer ATCC 11845 / DSM 15868 / JCM 9532 / NCTC 11014 | 139.5022 | 2.1446 | 16640 / 8046 | dbspec-15-Mar-2019-3-4-15.021 | UP000010093 |
| 25 | Hel cobacter pylori | 139.4446 | 2.1444 | 16640 / 6119 | dbspec-15-Mar-2019-3-4-27.173 | UP000012016 |
| 26 | Hel cobacter pylori | 139.0954 | 2.1433 | 16640 / 6036 | dbspec-15-Mar-2019-3-3-22.246 | UP000005535 |
| 27 | Arcobacter cryaerophilus | 138.9837 | 2.143 | 16640 / 6493 | dbspec-15-Mar-2019-3-9-48.44 | UP000239151 |
| 28 | Hel cobacter pylori | 138.7229 | 2.1421 | 16640 / 6175 | dbspec-15-Mar-2019-3-4-26.284 | UP000011920 |
| 29 | Hel cobacter pylori | 138.6284 | 2.1419 | 16640 / 6090 | dbspec-15-Mar-2019-3-4-26.799 | UP000011929 |
| 30 | Hel cobacter pylori | 138.5863 | 2.1417 | 16640 / 6070 | dbspec-15-Mar-2019-3-3-43.306 | UP000006748 |

#### # 6 - CLASSIFICATION SUBREPORT FOR SPECTRUM *'170_2019_ZBS6_Peter-Burkholderia_E153'*

##### Actual test mass spectrum

| | |
|---|---|
| **genus / species / strain:** | Burkholderia thailandensis E153 |
| **file id:** | 170_2019_ZBS6_Peter-Burkholderia_E153 |
| **type:** | 170 |
| **NCBI id (species level):** | NaN |
| **NCBI id (strain level):** | NaN |
| **growth time:** | 24 h |
| **growth temperature:** | 37Â°C |
| **growth conditions:** | no CO2 |
| **growth medium:** | Caso (Oxo d) |
| **sample treatment:** | SPEED, operator: A. Schneider |
| **spores:** | no |
| **concentration:** | not available |
| **extra info:** | biological replicate #1, technical replicate #2 |
| **calibration standard:** | not available |
| **measurement method:** | LC-MS1 spectrum, method not available |
| **customer:** | RKI/ZBS 6 |
| **measurement date/time:** | 2019-01_11:11:32.906+02:00 |
| **path to MS file:** | E:\Matlab |

##### List of best matches with test spectrum

| No | Genus/Species/Strain | Score | Log Score | Peak Numbers | Spectral ID | Proteome ID |
|---|---|---|---|---|---|---|
| 1 | Burkholderia thailandensis ATCC 700388 / DSM 13276 / CIP 106301 / E264 | 209.5739 | 2.3213 | 16064 / 23226 | dbspec-15-Mar-2019-3-2-18.411 | UP000001930 |
| 2 | Burkholderia thailandensis | 201.2091 | 2.3036 | 16064 / 22799 | dbspec-15-Mar-2019-3-9-35.914 | UP000235972 |
| 3 | Burkholderia pseudomallei 1710b | 162.4502 | 2.2107 | 16064 / 24945 | dbspec-15-Mar-2019-3-2-33.574 | UP000002700 |
| 4 | Burkholderia pseudomallei K96243 | 161.8706 | 2.2092 | 16064 / 23961 | dbspec-15-Mar-2019-3-1-58.302 | UP000000605 |
| 5 | Burkholderia pseudomallei | 161.84 | 2.2091 | 16064 / 24630 | dbspec-15-Mar-2019-3-6-4.424 | UP000032603 |
| 6 | Burkholderia pseudomallei | 161.6998 | 2.2087 | 16064 / 24475 | dbspec-15-Mar-2019-3-3-10.094 | UP000004875 |
| 7 | Burkholderia pseudomallei | 161.6436 | 2.2086 | 16064 / 24836 | dbspec-15-Mar-2019-3-2-21.734 | UP000002031 |
| 8 | Burkholderia pseudomallei | 161.2374 | 2.2075 | 16064 / 24544 | dbspec-15-Mar-2019-3-2-15.743 | UP000001812 |
| 9 | Burkholderia pseudomallei 1106a | 161.2322 | 2.2075 | 16064 / 24105 | dbspec-15-Mar-2019-3-3-42.854 | UP000006738 |
| 10 | Burkholderia pseudomallei | 161.1863 | 2.2073 | 16064 / 24832 | dbspec-15-Mar-2019-3-5-56.499 | UP000030449 |
| 11 | Burkholderia pseudomallei Pseudomonas pseudomallei | 161.057 | 2.207 | 16064 / 24887 | dbspec-15-Mar-2019-3-5-52.272 | UP000029527 |
| 12 | Burkholderia pseudomallei 668 | 160.5048 | 2.2055 | 16064 / 24129 | dbspec-15-Mar-2019-3-2-23.122 | UP000002153 |
| 13 | Burkholderia pseudomallei | 160.1313 | 2.2045 | 16064 / 24032 | dbspec-15-Mar-2019-3-3-23.479 | UP000005700 |
| 14 | Burkholderia pseudomallei 1026b | 159.9821 | 2.2041 | 16064 / 23916 | dbspec-15-Mar-2019-3-4-15.005 | UP000010087 |
| 15 | Burkholderia pseudomallei | 159.8261 | 2.2036 | 16064 / 23842 | dbspec-15-Mar-2019-3-3-0.858 | UP000004448 |
| 16 | Burkholderia pseudomallei | 157.9871 | 2.1986 | 16064 / 22928 | dbspec-15-Mar-2019-3-2-35.82 | UP000002781 |
| 17 | Burkholderia pseudomallei | 157.1046 | 2.1962 | 16064 / 21894 | dbspec-15-Mar-2019-3-3-7.02 | UP000004700 |
| 18 | Burkholderia thailandensis | 156.14 | 2.1935 | 16064 / 23446 | dbspec-15-Mar-2019-3-8-58.458 | UP000218714 |
| 19 | Burkholderia oklahomensis | 153.5594 | 2.1863 | 16064 / 24424 | dbspec-15-Mar-2019-3-10-40.31 | UP000254484 |
| 20 | Burkholderia mallei Pseudomonas mallei | 152.3437 | 2.1828 | 16064 / 19381 | dbspec-15-Mar-2019-3-10-40.934 | UP000254505 |
| 21 | Burkholderia mallei | 152.0545 | 2.182 | 16064 / 19432 | dbspec-15-Mar-2019-3-2-54.15 | UP000003995 |
| 22 | Burkholderia mallei ATCC 23344 | 151.2728 | 2.1798 | 16064 / 18442 | dbspec-15-Mar-2019-3-3-41.84 | UP000006693 |
| 23 | Burkholderia mallei NCTC 10229 | 150.1967 | 2.1767 | 16064 / 18362 | dbspec-15-Mar-2019-3-2-26.164 | UP000002283 |
| 24 | Hel cobacter pylori | 136.4052 | 2.1348 | 16064 / 6069 | dbspec-15-Mar-2019-3-4-25.333 | UP000011898 |
| 25 | Hel cobacter pylori | 136.3305 | 2.1346 | 16064 / 6180 | dbspec-15-Mar-2019-3-2-45.804 | UP000003402 |
| 26 | Hel cobacter pylori | 135.2801 | 2.1312 | 16064 / 5873 | dbspec-15-Mar-2019-3-4-51.104 | UP000015602 |
| 27 | Hel cobacter pylori B8 | 134.8856 | 2.13 | 16064 / 6271 | dbspec-15-Mar-2019-3-3-50.701 | UP000007091 |
| 28 | Peptoanaerobacter stomatis | 134.879 | 2.1299 | 16064 / 8144 | dbspec-15-Mar-2019-3-3-15.522 | UP000005244 |
| 29 | Hel cobacter pylori | 134.5519 | 2.1289 | 16064 / 6217 | dbspec-15-Mar-2019-3-3-21.154 | UP000005483 |
| 30 | Burkholderia lata ATCC 17760 / DSM 23089 / LMG 22485 / NCIMB 9086 / R18194 / 383 | 133.7329 | 2.1262 | 16064 / 31172 | dbspec-15-Mar-2019-3-2-33.948 | UP000002705 |

#### # 7 - CLASSIFICATION SUBREPORT FOR SPECTRUM *'174_2019_ZBS6_Peter-Burkholderia_E153'*

##### Actual test mass spectrum

| | |
|---|---|
| **genus / species / strain:** | Burkholderia thailandensis E153 |
| **file id:** | 174_2019_ZBS6_Peter-Burkholderia_E153 |
| **type:** | 174 |
| **NCBI id (species level):** | NaN |

| | |
|---|---|
| **NCBI id (strain level):** | NaN |
| **growth time:** | 24 h |
| **growth temperature:** | 37Â°C |
| **growth conditions:** | no CO2 |
| **growth medium:** | Caso (Oxo d) |
| **sample treatment:** | SPEED, operator: A. Schneider |
| **spores:** | no |
| **concentration:** | not available |
| **extra info:** | biological replicate #1, technical replicate #3 |
| **calibration standard:** | not available |
| **measurement method:** | LC-MS1 spectrum, method not available |
| **customer:** | RKI/ZBS 6 |
| **measurement date/time:** | 2019-01_11:11:32.906+02:00 |
| **path to MS file:** | E:\Matlab |

#### List of best matches with test spectrum

| No | Genus/Species/Strain | Score | Log Score | Peak Numbers | Spectral ID | Proteome ID |
|---|---|---|---|---|---|---|
| 1 | Burkholderia thailandensis ATCC 700388 / DSM 13276 / CIP 106301 / E264 | 210.9993 | 2.3243 | 16014 / 23226 | dbspec-15-Mar-2019-3-2-18.411 | UP000001930 |
| 2 | Burkholderia thailandensis | 201.4565 | 2.3042 | 16014 / 22799 | dbspec-15-Mar-2019-3-9-35.914 | UP000235972 |
| 3 | Burkholderia pseudomallei | 161.5364 | 2.2083 | 16014 / 24832 | dbspec-15-Mar-2019-3-5-56.499 | UP000030449 |
| 4 | Burkholderia pseudomallei 1710b | 161.1751 | 2.2073 | 16014 / 24945 | dbspec-15-Mar-2019-3-2-33.574 | UP000002700 |
| 5 | Burkholderia pseudomallei Pseudomonas pseudomallei | 161.1083 | 2.2071 | 16014 / 24887 | dbspec-15-Mar-2019-3-5-52.272 | UP000029527 |
| 6 | Burkholderia pseudomallei | 161.1075 | 2.2071 | 16014 / 24032 | dbspec-15-Mar-2019-3-3-23.479 | UP000005700 |
| 7 | Burkholderia pseudomallei | 160.9021 | 2.2066 | 16014 / 24544 | dbspec-15-Mar-2019-3-2-15.743 | UP000001812 |
| 8 | Burkholderia pseudomallei K96243 | 160.7014 | 2.206 | 16014 / 23961 | dbspec-15-Mar-2019-3-1-58.302 | UP000000605 |
| 9 | Burkholderia pseudomallei | 160.5717 | 2.2057 | 16014 / 24836 | dbspec-15-Mar-2019-3-2-21.734 | UP000002031 |
| 10 | Burkholderia pseudomallei | 160.537 | 2.2056 | 16014 / 24630 | dbspec-15-Mar-2019-3-6-4.424 | UP000032603 |
| 11 | Burkholderia pseudomallei | 160.2404 | 2.2048 | 16014 / 24475 | dbspec-15-Mar-2019-3-3-10.094 | UP000004875 |
| 12 | Burkholderia pseudomallei 1106a | 160.194 | 2.2046 | 16014 / 24105 | dbspec-15-Mar-2019-3-3-42.854 | UP000006738 |
| 13 | Burkholderia pseudomallei 668 | 160.0427 | 2.2042 | 16014 / 24129 | dbspec-15-Mar-2019-3-2-23.122 | UP000002153 |
| 14 | Burkholderia pseudomallei 1026b | 158.8803 | 2.2011 | 16014 / 23916 | dbspec-15-Mar-2019-3-4-15.005 | UP000010087 |
| 15 | Burkholderia pseudomallei | 158.6509 | 2.2004 | 16014 / 23842 | dbspec-15-Mar-2019-3-3-0.858 | UP000004448 |
| 16 | Burkholderia pseudomallei | 158.02 | 2.1987 | 16014 / 22928 | dbspec-15-Mar-2019-3-2-35.82 | UP000002781 |
| 17 | Burkholderia pseudomallei | 155.3304 | 2.1913 | 16014 / 21894 | dbspec-15-Mar-2019-3-3-7.02 | UP000004700 |
| 18 | Burkholderia thailandensis | 154.0315 | 2.1876 | 16014 / 23446 | dbspec-15-Mar-2019-3-8-58.458 | UP000218714 |
| 19 | Burkholderia mallei | 151.9185 | 2.1816 | 16014 / 19432 | dbspec-15-Mar-2019-3-2-54.15 | UP000003995 |
| 20 | Burkholderia mallei Pseudomonas mallei | 151.7057 | 2.181 | 16014 / 19381 | dbspec-15-Mar-2019-3-10-40.934 | UP000254505 |
| 21 | Burkholderia mallei ATCC 23344 | 150.353 | 2.1771 | 16014 / 18442 | dbspec-15-Mar-2019-3-3-41.84 | UP000006693 |
| 22 | Burkholderia mallei NCTC 10229 | 150.235 | 2.1768 | 16014 / 18362 | dbspec-15-Mar-2019-3-2-26.164 | UP000002283 |
| 23 | Burkholderia oklahomensis | 149.0914 | 2.1735 | 16014 / 24424 | dbspec-15-Mar-2019-3-10-40.31 | UP000254484 |
| 24 | Peptoanaerobacter stomatis | 133.3649 | 2.125 | 16014 / 8144 | dbspec-15-Mar-2019-3-3-15.522 | UP000005244 |
| 25 | Burkholderia reimsis | 133.3519 | 2.125 | 16014 / 30986 | dbspec-15-Mar-2019-3-10-32.495 | UP000252458 |
| 26 | Riemerella anatipestifer Moraxella anatipestifer | 133.1498 | 2.1243 | 16014 / 8629 | dbspec-15-Mar-2019-3-7-10.163 | UP000093661 |
| 27 | Arcobacter cryaerophilus | 132.7981 | 2.1232 | 16014 / 7684 | dbspec-15-Mar-2019-3-9-46.459 | UP000238649 |
| 28 | Neisseria lactam ca | 131.9494 | 2.1204 | 16014 / 7050 | dbspec-15-Mar-2019-3-10-38.532 | UP000254193 |
| 29 | Burkholderia cepacia | 131.8332 | 2.12 | 16014 / 29972 | dbspec-15-Mar-2019-3-10-52.322 | UP000263019 |
| 30 | Burkholderia contaminans | 131.6369 | 2.1194 | 16014 / 28983 | dbspec-15-Mar-2019-3-6-11.491 | UP000035664 |

---

### # 8 - CLASSIFICATION SUBREPORT FOR SPECTRUM *'244_2019_ZBS6_Peter-Burkholderia_E153'*

#### Actual test mass spectrum

| | |
|---|---|
| **genus / species / strain:** | Burkholderia thailandensis E153 |
| **file id:** | 244_2019_ZBS6_Peter-Burkholderia_E153 |
| **type:** | 244 |
| **NCBI id (species level):** | NaN |
| **NCBI id (strain level):** | NaN |
| **growth time:** | 24 h |
| **growth temperature:** | 37Â°C |
| **growth conditions:** | no CO2 |
| **growth medium:** | Caso (Oxo d) |
| **sample treatment:** | SPEED, operator: A. Schneider |
| **spores:** | no |
| **concentration:** | not available |
| **extra info:** | biological replicate #2, technical replicate #1 |
| **calibration standard:** | not available |
| **measurement method:** | LC-MS1 spectrum, method not available |
| **customer:** | RKI/ZBS 6 |
| **measurement date/time:** | 2019-01_11:11:32.906+02:00 |
| **path to MS file:** | E:\Matlab |

#### List of best matches with test spectrum

| No | Genus/Species/Strain | Score | Log Score | Peak Numbers | Spectral ID | Proteome ID |
|---|---|---|---|---|---|---|

| | | | | | | |
|---|---|---|---|---|---|---|
| 1 | Burkholderia thailandensis ATCC 700388 / DSM 13276 / CIP 106301 / E264 | 194.7787 | 2.2895 | 15121 / 23226 | dbspec-15-Mar-2019-3-2-18.411 | UP000001930 |
| 2 | Burkholderia thailandensis | 187.3294 | 2.2726 | 15121 / 22799 | dbspec-15-Mar-2019-3-9-35.914 | UP000235972 |
| 3 | Burkholderia pseudomallei 1710b | 154.5136 | 2.189 | 15121 / 24945 | dbspec-15-Mar-2019-3-2-33.574 | UP000002700 |
| 4 | Burkholderia pseudomallei 668 | 154.0598 | 2.1877 | 15121 / 24129 | dbspec-15-Mar-2019-3-2-23.122 | UP000002153 |
| 5 | Burkholderia pseudomallei | 153.6093 | 2.1864 | 15121 / 24544 | dbspec-15-Mar-2019-3-2-15.743 | UP000001812 |
| 6 | Burkholderia pseudomallei | 153.0988 | 2.185 | 15121 / 24475 | dbspec-15-Mar-2019-3-3-10.094 | UP000004875 |
| 7 | Burkholderia pseudomallei | 152.879 | 2.1843 | 15121 / 24032 | dbspec-15-Mar-2019-3-3-23.479 | UP000005700 |
| 8 | Burkholderia pseudomallei 1106a | 152.7693 | 2.184 | 15121 / 24105 | dbspec-15-Mar-2019-3-3-42.854 | UP000006738 |
| 9 | Burkholderia pseudomallei | 152.7641 | 2.184 | 15121 / 24836 | dbspec-15-Mar-2019-3-2-21.734 | UP000002031 |
| 10 | Burkholderia pseudomallei | 152.544 | 2.1834 | 15121 / 24832 | dbspec-15-Mar-2019-3-5-56.499 | UP000030449 |
| 11 | Burkholderia pseudomallei | 152.5281 | 2.1833 | 15121 / 23842 | dbspec-15-Mar-2019-3-3-0.858 | UP000004448 |
| 12 | Burkholderia pseudomallei Pseudomonas pseudomallei | 152.2114 | 2.1824 | 15121 / 24887 | dbspec-15-Mar-2019-3-5-52.272 | UP000029527 |
| 13 | Burkholderia thailandensis | 152.169 | 2.1823 | 15121 / 23446 | dbspec-15-Mar-2019-3-8-58.458 | UP000218714 |
| 14 | Burkholderia pseudomallei | 151.8682 | 2.1815 | 15121 / 24630 | dbspec-15-Mar-2019-3-6-4.424 | UP000032603 |
| 15 | Burkholderia pseudomallei K96243 | 151.7286 | 2.1811 | 15121 / 23961 | dbspec-15-Mar-2019-3-1-58.302 | UP000000605 |
| 16 | Burkholderia pseudomallei 1026b | 151.4772 | 2.1803 | 15121 / 23916 | dbspec-15-Mar-2019-3-4-15.005 | UP000010087 |
| 17 | Burkholderia pseudomallei | 150.2717 | 2.1769 | 15121 / 22928 | dbspec-15-Mar-2019-3-2-35.82 | UP000002781 |
| 18 | Burkholderia pseudomallei | 146.5738 | 2.1661 | 15121 / 21894 | dbspec-15-Mar-2019-3-3-7.02 | UP000004700 |
| 19 | Burkholderia oklahomensis | 145.308 | 2.1623 | 15121 / 24424 | dbspec-15-Mar-2019-3-10-40.31 | UP000254484 |
| 20 | Burkholderia mallei Pseudomonas mallei | 145.2536 | 2.1621 | 15121 / 19381 | dbspec-15-Mar-2019-3-10-40.934 | UP000254505 |
| 21 | Burkholderia mallei | 144.6409 | 2.1603 | 15121 / 19432 | dbspec-15-Mar-2019-3-2-54.15 | UP000003995 |
| 22 | Burkholderia mallei ATCC 23344 | 143.9866 | 2.1583 | 15121 / 18442 | dbspec-15-Mar-2019-3-3-41.84 | UP000006693 |
| 23 | Burkholderia mallei NCTC 10229 | 142.9792 | 2.1553 | 15121 / 18362 | dbspec-15-Mar-2019-3-2-26.164 | UP000002283 |
| 24 | Leptotr chia wadei F0279 | 134.525 | 2.1288 | 15121 / 7438 | dbspec-15-Mar-2019-3-4-56.907 | UP000016626 |
| 25 | Burkholderia pyrrocinia Pseudomonas pyrrocinia | 131.595 | 2.1192 | 15121 / 28460 | dbspec-15-Mar-2019-3-6-38.026 | UP000064483 |
| 26 | Burkholderia gladioli BSR3 | 131.4363 | 2.1187 | 15121 / 30592 | dbspec-15-Mar-2019-3-4-0.872 | UP000008316 |
| 27 | Burkholderia cenocepacia | 131.1849 | 2.1179 | 15121 / 30167 | dbspec-15-Mar-2019-3-6-38.822 | UP000065643 |
| 28 | Thermocrinis ruber | 130.7982 | 2.1166 | 15121 / 5633 | dbspec-15-Mar-2019-3-5-14.941 | UP000018914 |
| 29 | Hel cobacter pylori | 130.7857 | 2.1166 | 15121 / 5988 | dbspec-15-Mar-2019-3-4-51.353 | UP000015920 |
| 30 | Burkholderia cenocepacia | 130.6924 | 2.1163 | 15121 / 28696 | dbspec-15-Mar-2019-3-5-52.022 | UP000029413 |

---

#### # 9 - CLASSIFICATION SUBREPORT FOR SPECTRUM *'189_2019_ZBS6_Peter-Burkholderia_E125'*

##### Actual test mass spectrum

| | |
|---|---|
| **genus / species / strain:** | Burkholderia thailandensis E125 |
| **file id:** | 189_2019_ZBS6_Peter-Burkholderia_E125 |
| **type:** | 189 |
| **NCBI id (species level):** | NaN |
| **NCBI id (strain level):** | NaN |
| **growth time:** | 24 h |
| **growth temperature:** | 37Â°C |
| **growth conditions:** | no CO2 |
| **growth medium:** | Caso (Oxo d) |
| **sample treatment:** | SPEED, operator: A. Schneider |
| **spores:** | no |
| **concentration:** | not available |
| **extra info:** | biological replicate #1, technical replicate #1 |
| **calibration standard:** | not available |
| **measurement method:** | LC-MS1 spectrum, method not available |
| **customer:** | RKI/ZBS 6 |
| **measurement date/time:** | 2019-01_11:11:32.906+02:00 |
| **path to MS file:** | E:\Matlab |

##### List of best matches with test spectrum

| No | Genus/Species/Strain | Score | Log Score | Peak Numbers | Spectral ID | Proteome ID |
|---|---|---|---|---|---|---|
| 1 | Burkholderia thailandensis ATCC 700388 / DSM 13276 / CIP 106301 / E264 | 202.7592 | 2.307 | 15581 / 23226 | dbspec-15-Mar-2019-3-2-18.411 | UP000001930 |
| 2 | Burkholderia thailandensis | 195.6465 | 2.2915 | 15581 / 22799 | dbspec-15-Mar-2019-3-9-35.914 | UP000235972 |
| 3 | Burkholderia pseudomallei 1710b | 161.2297 | 2.2074 | 15581 / 24945 | dbspec-15-Mar-2019-3-2-33.574 | UP000002700 |
| 4 | Burkholderia pseudomallei 668 | 161.0323 | 2.2069 | 15581 / 24129 | dbspec-15-Mar-2019-3-2-23.122 | UP000002153 |
| 5 | Burkholderia pseudomallei | 160.0027 | 2.2041 | 15581 / 24544 | dbspec-15-Mar-2019-3-2-15.743 | UP000001812 |
| 6 | Burkholderia pseudomallei | 159.8997 | 2.2038 | 15581 / 24475 | dbspec-15-Mar-2019-3-3-10.094 | UP000004875 |
| 7 | Burkholderia pseudomallei Pseudomonas pseudomallei | 159.8932 | 2.2038 | 15581 / 24887 | dbspec-15-Mar-2019-3-5-52.272 | UP000029527 |
| 8 | Burkholderia pseudomallei | 159.8529 | 2.2037 | 15581 / 24032 | dbspec-15-Mar-2019-3-3-23.479 | UP000005700 |
| 9 | Burkholderia pseudomallei | 159.7486 | 2.2034 | 15581 / 24630 | dbspec-15-Mar-2019-3-6-4.424 | UP000032603 |
| 10 | Burkholderia pseudomallei K96243 | 159.6414 | 2.2031 | 15581 / 23961 | dbspec-15-Mar-2019-3-1-58.302 | UP000000605 |
| 11 | Burkholderia pseudomallei 1026b | 159.3121 | 2.2022 | 15581 / 23916 | dbspec-15-Mar-2019-3-4-15.005 | UP000010087 |
| 12 | Burkholderia pseudomallei 1106a | 159.2015 | 2.2019 | 15581 / 24105 | dbspec-15-Mar-2019-3-3-42.854 | UP000006738 |
| 13 | Burkholderia pseudomallei | 159.118 | 2.2017 | 15581 / 24832 | dbspec-15-Mar-2019-3-5-56.499 | UP000030449 |
| 14 | Burkholderia pseudomallei | 159.0503 | 2.2015 | 15581 / 24836 | dbspec-15-Mar-2019-3-2-21.734 | UP000002031 |
| 15 | Burkholderia pseudomallei | 158.7934 | 2.2008 | 15581 / 23842 | dbspec-15-Mar-2019-3-3-0.858 | UP000004448 |
| 16 | Burkholderia pseudomallei | 156.1078 | 2.1934 | 15581 / 22928 | dbspec-15-Mar-2019-3-2-35.82 | UP000002781 |
| 17 | Burkholderia thailandensis | 155.2498 | 2.191 | 15581 / 23446 | dbspec-15-Mar-2019-3-8-58.458 | UP000218714 |
| 18 | Burkholderia pseudomallei | 153.7177 | 2.1867 | 15581 / 21894 | dbspec-15-Mar-2019-3-3-7.02 | UP000004700 |
| 19 | Burkholderia mallei Pseudomonas mallei | 148.9483 | 2.173 | 15581 / 19381 | dbspec-15-Mar-2019-3-10-40.934 | UP000254505 |
| 20 | Burkholderia mallei | 148.5309 | 2.1718 | 15581 / 19432 | dbspec-15-Mar-2019-3-2-54.15 | UP000003995 |
| 21 | Burkholderia mallei ATCC 23344 | 147.5558 | 2.169 | 15581 / 18442 | dbspec-15-Mar-2019-3-3-41.84 | UP000006693 |

| | | | | | | |
|---|---|---|---|---|---|---|
| 22 | Burkholderia oklahomensis | 147.5208 | 2.1689 | 15581 / 24424 | dbspec-15-Mar-2019-3-10-40.31 | UP000254484 |
| 23 | Burkholderia mallei NCTC 10229 | 146.9007 | 2.167 | 15581 / 18362 | dbspec-15-Mar-2019-3-2-26.164 | UP000002283 |
| 24 | Hel cobacter pylori | 137.1433 | 2.1372 | 15581 / 6069 | dbspec-15-Mar-2019-3-4-25.333 | UP000011898 |
| 25 | Hel cobacter pylori | 135.7072 | 2.1326 | 15581 / 6078 | dbspec-15-Mar-2019-3-2-38.067 | UP000002997 |
| 26 | Hel cobacter pylori | 134.8268 | 2.1298 | 15581 / 6104 | dbspec-15-Mar-2019-3-4-27.314 | UP000012033 |
| 27 | Hel cobacter pylori | 134.7661 | 2.1296 | 15581 / 6107 | dbspec-15-Mar-2019-3-2-56.974 | UP000004225 |
| 28 | Hel cobacter pylori | 134.6858 | 2.1293 | 15581 / 6207 | dbspec-15-Mar-2019-3-3-21.981 | UP000005514 |
| 29 | Hel cobacter pylori | 134.6152 | 2.1291 | 15581 / 5871 | dbspec-15-Mar-2019-3-2-56.116 | UP000004153 |
| 30 | Burkholderia reimsis | 134.5186 | 2.1288 | 15581 / 30986 | dbspec-15-Mar-2019-3-10-32.495 | UP000252458 |

#### # 10 - CLASSIFICATION SUBREPORT FOR SPECTRUM *'190_2019_ZBS6_Peter-Burkholderia_E125'*

##### Actual test mass spectrum

| | |
|---|---|
| **genus / species / strain:** | Burkholderia thailandensis E125 |
| **file id:** | 190_2019_ZBS6_Peter-Burkholderia_E125 |
| **type:** | 190 |
| **NCBI id (species level):** | NaN |
| **NCBI id (strain level):** | NaN |
| **growth time:** | 24 h |
| **growth temperature:** | 37Â°C |
| **growth conditions:** | no CO2 |
| **growth medium:** | Caso (Oxo d) |
| **sample treatment:** | SPEED, operator: A. Schneider |
| **spores:** | no |
| **concentration:** | not available |
| **extra info:** | biological replicate #1, technical replicate #2 |
| **calibration standard:** | not available |
| **measurement method:** | LC-MS1 spectrum, method not available |
| **customer:** | RKI/ZBS 6 |
| **measurement date/time:** | 2019-01_11:11:32.906+02:00 |
| **path to MS file:** | E:\Matlab |

##### List of best matches with test spectrum

| No | Genus/Species/Strain | Score | Log Score | Peak Numbers | Spectral ID | Proteome ID |
|---|---|---|---|---|---|---|
| 1 | Burkholderia thailandensis ATCC 700388 / DSM 13276 / CIP 106301 / E264 | 198.8781 | 2.2986 | 15237 / 23226 | dbspec-15-Mar-2019-3-2-18.411 | UP000001930 |
| 2 | Burkholderia thailandensis | 191.8065 | 2.2829 | 15237 / 22799 | dbspec-15-Mar-2019-3-9-35.914 | UP000235972 |
| 3 | Burkholderia pseudomallei 1710b | 157.03 | 2.196 | 15237 / 24945 | dbspec-15-Mar-2019-3-2-33.574 | UP000002700 |
| 4 | Burkholderia pseudomallei | 156.1999 | 2.1937 | 15237 / 24032 | dbspec-15-Mar-2019-3-3-23.479 | UP000005700 |
| 5 | Burkholderia pseudomallei | 155.8203 | 2.1926 | 15237 / 24475 | dbspec-15-Mar-2019-3-3-10.094 | UP000004875 |
| 6 | Burkholderia pseudomallei K96243 | 155.6467 | 2.1921 | 15237 / 23961 | dbspec-15-Mar-2019-3-1-58.302 | UP000000605 |
| 7 | Burkholderia pseudomallei 1026b | 155.427 | 2.1915 | 15237 / 23916 | dbspec-15-Mar-2019-3-4-15.005 | UP000010087 |
| 8 | Burkholderia pseudomallei 668 | 155.3196 | 2.1912 | 15237 / 24129 | dbspec-15-Mar-2019-3-2-23.122 | UP000002153 |
| 9 | Burkholderia pseudomallei | 155.1664 | 2.1908 | 15237 / 24630 | dbspec-15-Mar-2019-3-6-4.424 | UP000032603 |
| 10 | Burkholderia pseudomallei 1106a | 155.1046 | 2.1906 | 15237 / 24105 | dbspec-15-Mar-2019-3-3-42.854 | UP000006738 |
| 11 | Burkholderia pseudomallei | 154.7512 | 2.1896 | 15237 / 24836 | dbspec-15-Mar-2019-3-2-21.734 | UP000002031 |
| 12 | Burkholderia pseudomallei | 154.5115 | 2.189 | 15237 / 24544 | dbspec-15-Mar-2019-3-2-15.743 | UP000001812 |
| 13 | Burkholderia pseudomallei | 154.1351 | 2.1879 | 15237 / 24832 | dbspec-15-Mar-2019-3-5-56.499 | UP000030449 |
| 14 | Burkholderia pseudomallei | 153.89 | 2.1872 | 15237 / 23842 | dbspec-15-Mar-2019-3-3-0.858 | UP000004448 |
| 15 | Burkholderia pseudomallei Pseudomonas pseudomallei | 153.8522 | 2.1871 | 15237 / 24887 | dbspec-15-Mar-2019-3-5-52.272 | UP000029527 |
| 16 | Burkholderia thailandensis | 152.2027 | 2.1824 | 15237 / 23446 | dbspec-15-Mar-2019-3-8-58.458 | UP000218714 |
| 17 | Burkholderia pseudomallei | 152.0272 | 2.1819 | 15237 / 22928 | dbspec-15-Mar-2019-3-2-35.82 | UP000002781 |
| 18 | Burkholderia pseudomallei | 148.9571 | 2.1731 | 15237 / 21894 | dbspec-15-Mar-2019-3-3-7.02 | UP000004700 |
| 19 | Burkholderia mallei | 144.6994 | 2.1605 | 15237 / 19432 | dbspec-15-Mar-2019-3-2-54.15 | UP000003995 |
| 20 | Burkholderia mallei Pseudomonas mallei | 144.4735 | 2.1598 | 15237 / 19381 | dbspec-15-Mar-2019-3-10-40.934 | UP000254505 |
| 21 | Burkholderia oklahomensis | 143.9648 | 2.1583 | 15237 / 24424 | dbspec-15-Mar-2019-3-10-40.31 | UP000254484 |
| 22 | Burkholderia mallei ATCC 23344 | 143.338 | 2.1564 | 15237 / 18442 | dbspec-15-Mar-2019-3-3-41.84 | UP000006693 |
| 23 | Burkholderia mallei NCTC 10229 | 141.4793 | 2.1507 | 15237 / 18362 | dbspec-15-Mar-2019-3-2-26.164 | UP000002283 |
| 24 | Hel cobacter pylori | 139.9822 | 2.1461 | 15237 / 6069 | dbspec-15-Mar-2019-3-4-25.333 | UP000011898 |
| 25 | Hel cobacter pylori | 137.0739 | 2.137 | 15237 / 6197 | dbspec-15-Mar-2019-3-4-30.995 | UP000012243 |
| 26 | Hel cobacter pylori | 136.3811 | 2.1348 | 15237 / 6231 | dbspec-15-Mar-2019-3-4-26.908 | UP000011951 |
| 27 | Hel cobacter pylori | 136.2264 | 2.1343 | 15237 / 6078 | dbspec-15-Mar-2019-3-2-38.067 | UP000002997 |
| 28 | Hel cobacter pylori | 135.2853 | 2.1313 | 15237 / 6107 | dbspec-15-Mar-2019-3-2-56.974 | UP000004225 |
| 29 | Hel cobacter pylori | 135.2722 | 2.1312 | 15237 / 5958 | dbspec-15-Mar-2019-3-4-26.924 | UP000011953 |
| 30 | Hel cobacter pylori | 134.8287 | 2.1298 | 15237 / 6108 | dbspec-15-Mar-2019-3-3-23.073 | UP000005601 |

#### # 11 - CLASSIFICATION SUBREPORT FOR SPECTRUM *'191_2019_ZBS6_Peter-Burkholderia_E125'*

##### Actual test mass spectrum

| | |
|---|---|
| **genus / species / strain:** | Burkholderia thailandensis E125 |
| **file id:** | 191_2019_ZBS6_Peter-Burkholderia_E125 |
| **type:** | 191 |
| **NCBI id (species level):** | NaN |

| | |
|---|---|
| **NCBI id (strain level):** | NaN |
| **growth time:** | 24 h |
| **growth temperature:** | 37Â°C |
| **growth conditions:** | no CO2 |
| **growth medium:** | Caso (Oxo d) |
| **sample treatment:** | SPEED, operator: A. Schneider |
| **spores:** | no |
| **concentration:** | not available |
| **extra info:** | biological replicate #1, technical replicate #3 |
| **calibration standard:** | not available |
| **measurement method:** | LC-MS1 spectrum, method not available |
| **customer:** | RKI/ZBS 6 |
| **measurement date/time:** | 2019-01_11:11:32.906+02:00 |
| **path to MS file:** | E:\Matlab |

#### List of best matches with test spectrum

| No | Genus/Species/Strain | Score | Log Score | Peak Numbers | Spectral ID | Proteome ID |
|---|---|---|---|---|---|---|
| 1 | Burkholderia thailandensis ATCC 700388 / DSM 13276 / CIP 106301 / E264 | 202.6297 | 2.3067 | 15429 / 23226 | dbspec-15-Mar-2019-3-2-18.411 | UP000001930 |
| 2 | Burkholderia thailandensis | 194.8451 | 2.2897 | 15429 / 22799 | dbspec-15-Mar-2019-3-9-35.914 | UP000235972 |
| 3 | Burkholderia pseudomallei 1710b | 164.1784 | 2.2153 | 15429 / 24945 | dbspec-15-Mar-2019-3-2-33.574 | UP000002700 |
| 4 | Burkholderia pseudomallei 668 | 162.4315 | 2.2107 | 15429 / 24129 | dbspec-15-Mar-2019-3-2-23.122 | UP000002153 |
| 5 | Burkholderia pseudomallei | 162.2452 | 2.2102 | 15429 / 24630 | dbspec-15-Mar-2019-3-6-4.424 | UP000032603 |
| 6 | Burkholderia pseudomallei | 162.1954 | 2.21 | 15429 / 24475 | dbspec-15-Mar-2019-3-3-10.094 | UP000004875 |
| 7 | Burkholderia pseudomallei | 161.915 | 2.2093 | 15429 / 24544 | dbspec-15-Mar-2019-3-2-15.743 | UP000001812 |
| 8 | Burkholderia pseudomallei Pseudomonas pseudomallei | 161.9097 | 2.2093 | 15429 / 24887 | dbspec-15-Mar-2019-3-5-52.272 | UP000029527 |
| 9 | Burkholderia pseudomallei K96243 | 161.7447 | 2.2088 | 15429 / 23961 | dbspec-15-Mar-2019-3-1-58.302 | UP000000605 |
| 10 | Burkholderia pseudomallei | 161.6516 | 2.2086 | 15429 / 24832 | dbspec-15-Mar-2019-3-5-56.499 | UP000030449 |
| 11 | Burkholderia pseudomallei 1026b | 161.6512 | 2.2086 | 15429 / 23916 | dbspec-15-Mar-2019-3-4-15.005 | UP000010087 |
| 12 | Burkholderia pseudomallei | 161.5763 | 2.2084 | 15429 / 24032 | dbspec-15-Mar-2019-3-3-23.479 | UP000005700 |
| 13 | Burkholderia pseudomallei | 161.502 | 2.2082 | 15429 / 24836 | dbspec-15-Mar-2019-3-2-21.734 | UP000002031 |
| 14 | Burkholderia pseudomallei 1106a | 161.1211 | 2.2072 | 15429 / 24105 | dbspec-15-Mar-2019-3-3-42.854 | UP000006738 |
| 15 | Burkholderia pseudomallei | 158.8786 | 2.2011 | 15429 / 22928 | dbspec-15-Mar-2019-3-2-35.82 | UP000002781 |
| 16 | Burkholderia pseudomallei | 158.591 | 2.2003 | 15429 / 23842 | dbspec-15-Mar-2019-3-3-0.858 | UP000004448 |
| 17 | Burkholderia thailandensis | 156.7974 | 2.1953 | 15429 / 23446 | dbspec-15-Mar-2019-3-8-58.458 | UP000218714 |
| 18 | Burkholderia pseudomallei | 154.8093 | 2.1898 | 15429 / 21894 | dbspec-15-Mar-2019-3-3-7.02 | UP000004700 |
| 19 | Burkholderia mallei | 149.4214 | 2.1744 | 15429 / 19432 | dbspec-15-Mar-2019-3-2-54.15 | UP000003995 |
| 20 | Burkholderia mallei Pseudomonas mallei | 149.1393 | 2.1736 | 15429 / 19381 | dbspec-15-Mar-2019-3-10-40.934 | UP000254505 |
| 21 | Burkholderia mallei ATCC 23344 | 148.3364 | 2.1712 | 15429 / 18442 | dbspec-15-Mar-2019-3-3-41.84 | UP000006693 |
| 22 | Burkholderia mallei NCTC 10229 | 147.6802 | 2.1693 | 15429 / 18362 | dbspec-15-Mar-2019-3-2-26.164 | UP000002283 |
| 23 | Burkholderia oklahomensis | 146.1737 | 2.1649 | 15429 / 24424 | dbspec-15-Mar-2019-3-10-40.31 | UP000254484 |
| 24 | Hel cobacter pylori | 138.6356 | 2.1419 | 15429 / 6069 | dbspec-15-Mar-2019-3-4-25.333 | UP000011898 |
| 25 | Hel cobacter pylori | 135.3376 | 2.1314 | 15429 / 6071 | dbspec-15-Mar-2019-3-3-24.165 | UP000005724 |
| 26 | Hel cobacter pylori ATCC 700392 / 26695 | 134.9372 | 2.1301 | 15429 / 6120 | dbspec-15-Mar-2019-3-1-54.87 | UP000000429 |
| 27 | Hel cobacter pylori | 134.7394 | 2.1295 | 15429 / 5873 | dbspec-15-Mar-2019-3-4-51.104 | UP000015602 |
| 28 | Hel cobacter pylori | 134.652 | 2.1292 | 15429 / 5754 | dbspec-15-Mar-2019-3-4-51.135 | UP000015663 |
| 29 | Hel cobacter pylori | 134.5515 | 2.1289 | 15429 / 5871 | dbspec-15-Mar-2019-3-2-56.116 | UP000004153 |
| 30 | Hel cobacter pylori | 134.4706 | 2.1286 | 15429 / 6046 | dbspec-15-Mar-2019-3-4-21.261 | UP000011225 |

### # 12 - CLASSIFICATION SUBREPORT FOR SPECTRUM *'246_2019_ZBS6_Peter-Burkholderia_E125'*

#### Actual test mass spectrum

| | |
|---|---|
| **genus / species / strain:** | Burkholderia thailandensis E125 |
| **file id:** | 246_2019_ZBS6_Peter-Burkholderia_E125 |
| **type:** | 246 |
| **NCBI id (species level):** | NaN |
| **NCBI id (strain level):** | NaN |
| **growth time:** | 24 h |
| **growth temperature:** | 37Â°C |
| **growth conditions:** | no CO2 |
| **growth medium:** | Caso (Oxo d) |
| **sample treatment:** | SPEED, operator: A. Schneider |
| **spores:** | no |
| **concentration:** | not available |
| **extra info:** | biological replicate #2, technical replicate #1 |
| **calibration standard:** | not available |
| **measurement method:** | LC-MS1 spectrum, method not available |
| **customer:** | RKI/ZBS 6 |
| **measurement date/time:** | 2019-01_11:11:32.906+02:00 |
| **path to MS file:** | E:\Matlab |

#### List of best matches with test spectrum

| No | Genus/Species/Strain | Score | Log Score | Peak Numbers | Spectral ID | Proteome ID |
|---|---|---|---|---|---|---|

| | | | | | | |
|---|---|---|---|---|---|---|
| 1 | Burkholderia thailandensis ATCC 700388 / DSM 13276 / CIP 106301 / E264 | 190.1885 | 2.2792 | 14738 / 23226 | dbspec-15-Mar-2019-3-2-18.411 | UP000001930 |
| 2 | Burkholderia thailandensis | 185.4785 | 2.2683 | 14738 / 22799 | dbspec-15-Mar-2019-3-9-35.914 | UP000235972 |
| 3 | Burkholderia pseudomallei 1710b | 154.1379 | 2.1879 | 14738 / 24945 | dbspec-15-Mar-2019-3-2-33.574 | UP000002700 |
| 4 | Burkholderia pseudomallei | 153.8421 | 2.1871 | 14738 / 24630 | dbspec-15-Mar-2019-3-6-4.424 | UP000032603 |
| 5 | Burkholderia pseudomallei Pseudomonas pseudomallei | 153.6732 | 2.1866 | 14738 / 24887 | dbspec-15-Mar-2019-3-5-52.272 | UP000029527 |
| 6 | Burkholderia pseudomallei | 153.6088 | 2.1864 | 14738 / 24475 | dbspec-15-Mar-2019-3-3-10.094 | UP000004875 |
| 7 | Burkholderia pseudomallei | 153.4526 | 2.186 | 14738 / 24032 | dbspec-15-Mar-2019-3-3-23.479 | UP000005700 |
| 8 | Burkholderia pseudomallei 668 | 153.2924 | 2.1855 | 14738 / 24129 | dbspec-15-Mar-2019-3-2-23.122 | UP000002153 |
| 9 | Burkholderia pseudomallei | 153.1774 | 2.1852 | 14738 / 24544 | dbspec-15-Mar-2019-3-2-15.743 | UP000001812 |
| 10 | Burkholderia pseudomallei K96243 | 152.971 | 2.1846 | 14738 / 23961 | dbspec-15-Mar-2019-3-1-58.302 | UP000000605 |
| 11 | Burkholderia pseudomallei | 152.6443 | 2.1837 | 14738 / 24832 | dbspec-15-Mar-2019-3-5-56.499 | UP000030449 |
| 12 | Burkholderia pseudomallei 1106a | 152.1276 | 2.1822 | 14738 / 24105 | dbspec-15-Mar-2019-3-3-42.854 | UP000006738 |
| 13 | Burkholderia pseudomallei | 152.0932 | 2.1821 | 14738 / 24836 | dbspec-15-Mar-2019-3-2-21.734 | UP000002031 |
| 14 | Burkholderia pseudomallei 1026b | 151.8721 | 2.1815 | 14738 / 23916 | dbspec-15-Mar-2019-3-4-15.005 | UP000010087 |
| 15 | Burkholderia pseudomallei | 151.275 | 2.1798 | 14738 / 23842 | dbspec-15-Mar-2019-3-3-0.858 | UP000004448 |
| 16 | Burkholderia pseudomallei | 148.6701 | 2.1722 | 14738 / 22928 | dbspec-15-Mar-2019-3-2-35.82 | UP000002781 |
| 17 | Burkholderia thailandensis | 148.2085 | 2.1709 | 14738 / 23446 | dbspec-15-Mar-2019-3-8-58.458 | UP000218714 |
| 18 | Burkholderia pseudomallei | 148.1891 | 2.1708 | 14738 / 21894 | dbspec-15-Mar-2019-3-3-7.02 | UP000004700 |
| 19 | Burkholderia mallei | 144.975 | 2.1613 | 14738 / 19432 | dbspec-15-Mar-2019-3-2-54.15 | UP000003995 |
| 20 | Burkholderia mallei Pseudomonas mallei | 144.2123 | 2.159 | 14738 / 19381 | dbspec-15-Mar-2019-3-10-40.934 | UP000254505 |
| 21 | Burkholderia mallei ATCC 23344 | 143.2958 | 2.1562 | 14738 / 18442 | dbspec-15-Mar-2019-3-3-41.84 | UP000006693 |
| 22 | Burkholderia mallei NCTC 10229 | 142.6723 | 2.1543 | 14738 / 18362 | dbspec-15-Mar-2019-3-2-26.164 | UP000002283 |
| 23 | Burkholderia oklahomensis | 141.9838 | 2.1522 | 14738 / 24424 | dbspec-15-Mar-2019-3-10-40.31 | UP000254484 |
| 24 | Arcobacter cryaerophilus | 132.3803 | 2.1218 | 14738 / 6493 | dbspec-15-Mar-2019-3-9-48.44 | UP000239151 |
| 25 | Burkholderia vietnamiensis G4 / LMG 22486 | 132.0052 | 2.1206 | 14738 / 27924 | dbspec-15-Mar-2019-3-2-26.18 | UP000002287 |
| 26 | Leptotr chia wadei F0279 | 130.7318 | 2.1164 | 14738 / 7438 | dbspec-15-Mar-2019-3-4-56.907 | UP000016626 |
| 27 | Burkholderia contaminans | 130.4382 | 2.1154 | 14738 / 28983 | dbspec-15-Mar-2019-3-6-11.491 | UP000035664 |
| 28 | Burkholderia cepacia Pseudomonas cepacia | 130.3425 | 2.1151 | 14738 / 29003 | dbspec-15-Mar-2019-3-5-52.038 | UP000029423 |
| 29 | Burkholderia cenocepacia | 129.5662 | 2.1125 | 14738 / 28696 | dbspec-15-Mar-2019-3-5-52.022 | UP000029413 |
| 30 | Burkholderia puraquae | 128.9945 | 2.1106 | 14738 / 28790 | dbspec-15-Mar-2019-3-8-4.997 | UP000193146 |

---

#### # 13 - CLASSIFICATION SUBREPORT FOR SPECTRUM *'167_2019_ZBS6_Peter-Burkholderia_20219'*

##### Actual test mass spectrum

| | |
|---|---|
| **genus / species / strain:** | Burkholderia thailandensis LMG 20219 |
| **file id:** | 167_2019_ZBS6_Peter-Burkholderia_20219 |
| **type:** | 167 |
| **NCBI id (species level):** | NaN |
| **NCBI id (strain level):** | NaN |
| **growth time:** | 24 h |
| **growth temperature:** | 37Â°C |
| **growth conditions:** | no CO2 |
| **growth medium:** | Caso (Oxo d) |
| **sample treatment:** | SPEED, operator: A. Schneider |
| **spores:** | no |
| **concentration:** | not available |
| **extra info:** | biological replicate #1, technical replicate #1 |
| **calibration standard:** | not available |
| **measurement method:** | LC-MS1 spectrum, method not available |
| **customer:** | RKI/ZBS 6 |
| **measurement date/time:** | 2019-01_11:11:32.906+02:00 |
| **path to MS file:** | E:\Matlab |

##### List of best matches with test spectrum

| No | Genus/Species/Strain | Score | Log Score | Peak Numbers | Spectral ID | Proteome ID |
|---|---|---|---|---|---|---|
| 1 | Burkholderia thailandensis ATCC 700388 / DSM 13276 / CIP 106301 / E264 | 220.4719 | 2.3434 | 16484 / 23226 | dbspec-15-Mar-2019-3-2-18.411 | UP000001930 |
| 2 | Burkholderia thailandensis | 207.1905 | 2.3164 | 16484 / 22799 | dbspec-15-Mar-2019-3-9-35.914 | UP000235972 |
| 3 | Burkholderia pseudomallei 1710b | 167.539 | 2.2241 | 16484 / 24945 | dbspec-15-Mar-2019-3-2-33.574 | UP000002700 |
| 4 | Burkholderia pseudomallei | 167.2872 | 2.2235 | 16484 / 24630 | dbspec-15-Mar-2019-3-6-4.424 | UP000032603 |
| 5 | Burkholderia pseudomallei 668 | 167.1392 | 2.2231 | 16484 / 24129 | dbspec-15-Mar-2019-3-2-23.122 | UP000002153 |
| 6 | Burkholderia pseudomallei 1106a | 167.0862 | 2.2229 | 16484 / 24105 | dbspec-15-Mar-2019-3-3-42.854 | UP000006738 |
| 7 | Burkholderia pseudomallei | 167.0802 | 2.2229 | 16484 / 24475 | dbspec-15-Mar-2019-3-3-10.094 | UP000004875 |
| 8 | Burkholderia pseudomallei | 167.0763 | 2.2229 | 16484 / 24032 | dbspec-15-Mar-2019-3-3-23.479 | UP000005700 |
| 9 | Burkholderia pseudomallei | 166.9944 | 2.2227 | 16484 / 24836 | dbspec-15-Mar-2019-3-2-21.734 | UP000002031 |
| 10 | Burkholderia pseudomallei K96243 | 166.7313 | 2.222 | 16484 / 23961 | dbspec-15-Mar-2019-3-1-58.302 | UP000000605 |
| 11 | Burkholderia pseudomallei Pseudomonas pseudomallei | 166.3988 | 2.2212 | 16484 / 24887 | dbspec-15-Mar-2019-3-5-52.272 | UP000029527 |
| 12 | Burkholderia pseudomallei | 166.2037 | 2.2206 | 16484 / 24544 | dbspec-15-Mar-2019-3-2-15.743 | UP000001812 |
| 13 | Burkholderia pseudomallei | 166.203 | 2.2206 | 16484 / 24832 | dbspec-15-Mar-2019-3-5-56.499 | UP000030449 |
| 14 | Burkholderia pseudomallei | 165.6135 | 2.2191 | 16484 / 23842 | dbspec-15-Mar-2019-3-3-0.858 | UP000004448 |
| 15 | Burkholderia pseudomallei 1026b | 165.0258 | 2.2176 | 16484 / 23916 | dbspec-15-Mar-2019-3-4-15.005 | UP000010087 |
| 16 | Burkholderia pseudomallei | 163.8974 | 2.2146 | 16484 / 22928 | dbspec-15-Mar-2019-3-2-35.82 | UP000002781 |
| 17 | Burkholderia thailandensis | 161.9119 | 2.2093 | 16484 / 23446 | dbspec-15-Mar-2019-3-8-58.458 | UP000218714 |
| 18 | Burkholderia pseudomallei | 160.4608 | 2.2054 | 16484 / 21894 | dbspec-15-Mar-2019-3-3-7.02 | UP000004700 |
| 19 | Burkholderia mallei Pseudomonas mallei | 159.3626 | 2.2024 | 16484 / 19381 | dbspec-15-Mar-2019-3-10-40.934 | UP000254505 |
| 20 | Burkholderia mallei | 158.2673 | 2.1994 | 16484 / 19432 | dbspec-15-Mar-2019-3-2-54.15 | UP000003995 |
| 21 | Burkholderia mallei ATCC 23344 | 156.4969 | 2.1945 | 16484 / 18442 | dbspec-15-Mar-2019-3-3-41.84 | UP000006693 |

| | | | | | | |
|---|---|---|---|---|---|---|
| 22 | Burkholderia mallei NCTC 10229 | 156.403 | 2.1942 | 16484 / 18362 | dbspec-15-Mar-2019-3-2-26.164 | UP000002283 |
| 23 | Burkholderia oklahomensis | 150.112 | 2.1764 | 16484 / 24424 | dbspec-15-Mar-2019-3-10-40.31 | UP000254484 |
| 24 | Riemerella anatipestifer ATCC 11845 / DSM 15868 / JCM 9532 / NCTC 11014 | 138.9752 | 2.1429 | 16484 / 8046 | dbspec-15-Mar-2019-3-4-15.021 | UP000010093 |
| 25 | Hel cobacter pylori | 138.1879 | 2.1405 | 16484 / 6100 | dbspec-15-Mar-2019-3-2-42.185 | UP000003215 |
| 26 | Hel cobacter fennelliae | 136.4647 | 2.135 | 16484 / 8395 | dbspec-15-Mar-2019-3-10-24.773 | UP000250166 |
| 27 | Arcobacter cryaerophilus | 136.2234 | 2.1343 | 16484 / 6493 | dbspec-15-Mar-2019-3-9-48.44 | UP000239151 |
| 28 | Sanguibacteroides justesenii | 136.0382 | 2.1337 | 16484 / 6412 | dbspec-15-Mar-2019-3-10-13.151 | UP000248308 |
| 29 | Streptobacillus moniliformis | 136.0051 | 2.1336 | 16484 / 5765 | dbspec-15-Mar-2019-3-10-29.921 | UP000251650 |
| 30 | Burkholderia dolosa | 135.7047 | 2.1326 | 16484 / 9075 | dbspec-15-Mar-2019-3-6-29.977 | UP000053482 |

#### # 14 - CLASSIFICATION SUBREPORT FOR SPECTRUM *'171_2019_ZBS6_Peter-Burkholderia_20219'*

##### Actual test mass spectrum

| | |
|---|---|
| **genus / species / strain:** | Burkholderia thailandensis LMG 20219 |
| **file id:** | 171_2019_ZBS6_Peter-Burkholderia_20219 |
| **type:** | 171 |
| **NCBI id (species level):** | NaN |
| **NCBI id (strain level):** | NaN |
| **growth time:** | 24 h |
| **growth temperature:** | 37Â°C |
| **growth conditions:** | no CO2 |
| **growth medium:** | Caso (Oxo d) |
| **sample treatment:** | SPEED, operator: A. Schneider |
| **spores:** | no |
| **concentration:** | not available |
| **extra info:** | biological replicate #1, technical replicate #2 |
| **calibration standard:** | not available |
| **measurement method:** | LC-MS1 spectrum, method not available |
| **customer:** | RKI/ZBS 6 |
| **measurement date/time:** | 2019-01_11:11:32.906+02:00 |
| **path to MS file:** | E:\Matlab |

##### List of best matches with test spectrum

| No | Genus/Species/Strain | Score | Log Score | Peak Numbers | Spectral ID | Proteome ID |
|---|---|---|---|---|---|---|
| 1 | Burkholderia thailandensis ATCC 700388 / DSM 13276 / CIP 106301 / E264 | 212.2679 | 2.3269 | 16166 / 23226 | dbspec-15-Mar-2019-3-2-18.411 | UP000001930 |
| 2 | Burkholderia thailandensis | 199.9794 | 2.301 | 16166 / 22799 | dbspec-15-Mar-2019-3-9-35.914 | UP000235972 |
| 3 | Burkholderia pseudomallei 1710b | 164.3614 | 2.2158 | 16166 / 24945 | dbspec-15-Mar-2019-3-2-33.574 | UP000002700 |
| 4 | Burkholderia pseudomallei K96243 | 163.9198 | 2.2146 | 16166 / 23961 | dbspec-15-Mar-2019-3-1-58.302 | UP000000605 |
| 5 | Burkholderia pseudomallei | 163.8575 | 2.2145 | 16166 / 24475 | dbspec-15-Mar-2019-3-3-10.094 | UP000004875 |
| 6 | Burkholderia pseudomallei 668 | 163.713 | 2.2141 | 16166 / 24129 | dbspec-15-Mar-2019-3-2-23.122 | UP000002153 |
| 7 | Burkholderia pseudomallei Pseudomonas pseudomallei | 163.3984 | 2.2132 | 16166 / 24887 | dbspec-15-Mar-2019-3-5-52.272 | UP000029527 |
| 8 | Burkholderia pseudomallei | 163.2068 | 2.2127 | 16166 / 24544 | dbspec-15-Mar-2019-3-2-15.743 | UP000001812 |
| 9 | Burkholderia pseudomallei 1026b | 162.9153 | 2.212 | 16166 / 23916 | dbspec-15-Mar-2019-3-4-15.005 | UP000010087 |
| 10 | Burkholderia pseudomallei | 162.4642 | 2.2108 | 16166 / 24832 | dbspec-15-Mar-2019-3-5-56.499 | UP000030449 |
| 11 | Burkholderia pseudomallei | 162.361 | 2.2105 | 16166 / 24630 | dbspec-15-Mar-2019-3-6-4.424 | UP000032603 |
| 12 | Burkholderia pseudomallei | 162.14 | 2.2099 | 16166 / 24032 | dbspec-15-Mar-2019-3-3-23.479 | UP000005700 |
| 13 | Burkholderia pseudomallei 1106a | 161.8597 | 2.2091 | 16166 / 24105 | dbspec-15-Mar-2019-3-3-42.854 | UP000006738 |
| 14 | Burkholderia pseudomallei | 161.8292 | 2.2091 | 16166 / 23842 | dbspec-15-Mar-2019-3-3-0.858 | UP000004448 |
| 15 | Burkholderia pseudomallei | 161.7899 | 2.209 | 16166 / 24836 | dbspec-15-Mar-2019-3-2-21.734 | UP000002031 |
| 16 | Burkholderia pseudomallei | 160.9084 | 2.2066 | 16166 / 22928 | dbspec-15-Mar-2019-3-2-35.82 | UP000002781 |
| 17 | Burkholderia pseudomallei | 157.8493 | 2.1982 | 16166 / 21894 | dbspec-15-Mar-2019-3-3-7.02 | UP000004700 |
| 18 | Burkholderia thailandensis | 156.6625 | 2.195 | 16166 / 23446 | dbspec-15-Mar-2019-3-8-58.458 | UP000218714 |
| 19 | Burkholderia mallei | 153.4354 | 2.1859 | 16166 / 19432 | dbspec-15-Mar-2019-3-2-54.15 | UP000003995 |
| 20 | Burkholderia mallei Pseudomonas mallei | 153.3505 | 2.1857 | 16166 / 19381 | dbspec-15-Mar-2019-3-10-40.934 | UP000254505 |
| 21 | Burkholderia mallei ATCC 23344 | 151.5613 | 2.1806 | 16166 / 18442 | dbspec-15-Mar-2019-3-3-41.84 | UP000006693 |
| 22 | Burkholderia mallei NCTC 10229 | 151.1978 | 2.1795 | 16166 / 18362 | dbspec-15-Mar-2019-3-2-26.164 | UP000002283 |
| 23 | Burkholderia oklahomensis | 150.0692 | 2.1763 | 16166 / 24424 | dbspec-15-Mar-2019-3-10-40.31 | UP000254484 |
| 24 | Riemerella anatipestifer ATCC 11845 / DSM 15868 / JCM 9532 / NCTC 11014 | 137.1683 | 2.1373 | 16166 / 8046 | dbspec-15-Mar-2019-3-4-15.021 | UP000010093 |
| 25 | Arcobacter cryaerophilus | 135.2826 | 2.1312 | 16166 / 6493 | dbspec-15-Mar-2019-3-9-48.44 | UP000239151 |
| 26 | Burkholderia ubonensis | 134.0125 | 2.1271 | 16166 / 27094 | dbspec-15-Mar-2019-3-6-35.016 | UP000058930 |
| 27 | Fimbriiglobus ruber | 133.9486 | 2.1269 | 16166 / 32923 | dbspec-15-Mar-2019-3-8-48.037 | UP000214646 |
| 28 | Hel cobacter pylori | 133.8357 | 2.1266 | 16166 / 6083 | dbspec-15-Mar-2019-3-3-11.919 | UP000004974 |
| 29 | Arcobacter cryaerophilus | 133.6917 | 2.1261 | 16166 / 7684 | dbspec-15-Mar-2019-3-9-46.459 | UP000238649 |
| 30 | Arcobacter cryaerophilus | 133.674 | 2.126 | 16166 / 7372 | dbspec-15-Mar-2019-3-9-46.88 | UP000238811 |

#### # 15 - CLASSIFICATION SUBREPORT FOR SPECTRUM *'175_2019_ZBS6_Peter-Burkholderia_20219'*

##### Actual test mass spectrum

| | |
|---|---|
| **genus / species / strain:** | Burkholderia thailandensis LMG 20219 |
| **file id:** | 175_2019_ZBS6_Peter-Burkholderia_20219 |

| | |
|---|---|
| **type:** | 175 |
| **NCBI id (species level):** | NaN |
| **NCBI id (strain level):** | NaN |
| **growth time:** | 24 h |
| **growth temperature:** | 37Â°C |
| **growth conditions:** | no CO2 |
| **growth medium:** | Caso (Oxo d) |
| **sample treatment:** | SPEED, operator: A. Schneider |
| **spores:** | no |
| **concentration:** | not available |
| **extra info:** | biological replicate #1, technical replicate #3 |
| **calibration standard:** | not available |
| **measurement method:** | LC-MS1 spectrum, method not available |
| **customer:** | RKI/ZBS 6 |
| **measurement date/time:** | 2019-01_11:11:32.906+02:00 |
| **path to MS file:** | E:\Matlab |

#### List of best matches with test spectrum

| No | Genus/Species/Strain | Score | Log Score | Peak Numbers | Spectral ID | Proteome ID |
|---|---|---|---|---|---|---|
| 1 | Burkholderia thailandensis ATCC 700388 / DSM 13276 / CIP 106301 / E264 | 221.6617 | 2.3457 | 16119 / 23226 | dbspec-15-Mar-2019-3-2-18.411 | UP000001930 |
| 2 | Burkholderia thailandensis | 206.7336 | 2.3154 | 16119 / 22799 | dbspec-15-Mar-2019-3-9-35.914 | UP000235972 |
| 3 | Burkholderia pseudomallei 1710b | 167.497 | 2.224 | 16119 / 24945 | dbspec-15-Mar-2019-3-2-33.574 | UP000002700 |
| 4 | Burkholderia pseudomallei K96243 | 166.0223 | 2.2202 | 16119 / 23961 | dbspec-15-Mar-2019-3-1-58.302 | UP000000605 |
| 5 | Burkholderia pseudomallei | 166.0008 | 2.2201 | 16119 / 24832 | dbspec-15-Mar-2019-3-5-56.499 | UP000030449 |
| 6 | Burkholderia pseudomallei 668 | 165.7513 | 2.2195 | 16119 / 24129 | dbspec-15-Mar-2019-3-2-23.122 | UP000002153 |
| 7 | Burkholderia pseudomallei Pseudomonas pseudomallei | 165.7481 | 2.2194 | 16119 / 24887 | dbspec-15-Mar-2019-3-5-52.272 | UP000029527 |
| 8 | Burkholderia pseudomallei | 165.7424 | 2.2194 | 16119 / 24544 | dbspec-15-Mar-2019-3-2-15.743 | UP000001812 |
| 9 | Burkholderia pseudomallei | 165.3683 | 2.2185 | 16119 / 24630 | dbspec-15-Mar-2019-3-6-4.424 | UP000032603 |
| 10 | Burkholderia pseudomallei 1106a | 165.2733 | 2.2182 | 16119 / 24105 | dbspec-15-Mar-2019-3-3-42.854 | UP000006738 |
| 11 | Burkholderia pseudomallei | 164.7635 | 2.2169 | 16119 / 24836 | dbspec-15-Mar-2019-3-2-21.734 | UP000002031 |
| 12 | Burkholderia pseudomallei 1026b | 164.7317 | 2.2168 | 16119 / 23916 | dbspec-15-Mar-2019-3-4-15.005 | UP000010087 |
| 13 | Burkholderia pseudomallei | 164.6973 | 2.2167 | 16119 / 24032 | dbspec-15-Mar-2019-3-3-23.479 | UP000005700 |
| 14 | Burkholderia pseudomallei | 163.9476 | 2.2147 | 16119 / 24475 | dbspec-15-Mar-2019-3-3-10.094 | UP000004875 |
| 15 | Burkholderia pseudomallei | 161.6676 | 2.2086 | 16119 / 23842 | dbspec-15-Mar-2019-3-3-0.858 | UP000004448 |
| 16 | Burkholderia pseudomallei | 161.5387 | 2.2083 | 16119 / 22928 | dbspec-15-Mar-2019-3-2-35.82 | UP000002781 |
| 17 | Burkholderia pseudomallei | 159.8418 | 2.2037 | 16119 / 21894 | dbspec-15-Mar-2019-3-3-7.02 | UP000004700 |
| 18 | Burkholderia thailandensis | 159.5739 | 2.203 | 16119 / 23446 | dbspec-15-Mar-2019-3-8-58.458 | UP000218714 |
| 19 | Burkholderia mallei Pseudomonas mallei | 157.5733 | 2.1975 | 16119 / 19381 | dbspec-15-Mar-2019-3-10-40.934 | UP000254505 |
| 20 | Burkholderia mallei | 156.405 | 2.1943 | 16119 / 19432 | dbspec-15-Mar-2019-3-2-54.15 | UP000003995 |
| 21 | Burkholderia mallei ATCC 23344 | 155.492 | 2.1917 | 16119 / 18442 | dbspec-15-Mar-2019-3-3-41.84 | UP000006693 |
| 22 | Burkholderia mallei NCTC 10229 | 154.4311 | 2.1887 | 16119 / 18362 | dbspec-15-Mar-2019-3-2-26.164 | UP000002283 |
| 23 | Burkholderia oklahomensis | 150.8957 | 2.1787 | 16119 / 24424 | dbspec-15-Mar-2019-3-10-40.31 | UP000254484 |
| 24 | Arcobacter cryaerophilus | 136.6654 | 2.1357 | 16119 / 7620 | dbspec-15-Mar-2019-3-9-50.437 | UP000239646 |
| 25 | Bartonella taylorii | 135.8603 | 2.1331 | 16119 / 6736 | dbspec-15-Mar-2019-3-2-32.81 | UP000002648 |
| 26 | Campylobacter lanienae | 135.2038 | 2.131 | 16119 / 6128 | dbspec-15-Mar-2019-3-8-46.571 | UP000202031 |
| 27 | Porphyromonas cangingivalis | 134.4175 | 2.1285 | 16119 / 8218 | dbspec-15-Mar-2019-3-7-52.345 | UP000189956 |
| 28 | Burkholderia reimsis | 134.0152 | 2.1272 | 16119 / 30986 | dbspec-15-Mar-2019-3-10-32.495 | UP000252458 |
| 29 | Myxococcus virescens | 133.9869 | 2.1271 | 16119 / 31646 | dbspec-15-Mar-2019-3-8-27.695 | UP000198717 |
| 30 | Burkholderia lata ATCC 17760 / DSM 23089 / LMG 22485 / NCIMB 9086 / R18194 / 383 | 133.9375 | 2.1269 | 16119 / 31172 | dbspec-15-Mar-2019-3-2-33.948 | UP000002705 |

---

### # 16 - CLASSIFICATION SUBREPORT FOR SPECTRUM *'247_2019_ZBS6_Peter-Burkholderia_LMG20219'*

#### Actual test mass spectrum

| | |
|---|---|
| **genus / species / strain:** | Burkholderia thailandensis LMG 20219 |
| **file id:** | 247_2019_ZBS6_Peter-Burkholderia_LMG20219 |
| **type:** | 247 |
| **NCBI id (species level):** | NaN |
| **NCBI id (strain level):** | NaN |
| **growth time:** | 24 h |
| **growth temperature:** | 37Â°C |
| **growth conditions:** | no CO2 |
| **growth medium:** | Caso (Oxo d) |
| **sample treatment:** | SPEED, operator: A. Schneider |
| **spores:** | no |
| **concentration:** | not available |
| **extra info:** | biological replicate #2, technical replicate #1 |
| **calibration standard:** | not available |
| **measurement method:** | LC-MS1 spectrum, method not available |
| **customer:** | RKI/ZBS 6 |
| **measurement date/time:** | 2019-01_11:11:32.906+02:00 |
| **path to MS file:** | E:\Matlab |

#### List of best matches with test spectrum

| No | Genus/Species/Strain | Score | Log Score | Peak Numbers | Spectral ID | Proteome ID |
|---|---|---|---|---|---|---|
| 1 | Burkholderia thailandensis ATCC 700388 / DSM 13276 / CIP 106301 / E264 | 203.342 | 2.3082 | 14898 / 23226 | dbspec-15-Mar-2019-3-2-18.411 | UP000001930 |
| 2 | Burkholderia thailandensis | 192.4561 | 2.2843 | 14898 / 22799 | dbspec-15-Mar-2019-3-9-35.914 | UP000235972 |
| 3 | Burkholderia pseudomallei 1710b | 161.0222 | 2.2069 | 14898 / 24945 | dbspec-15-Mar-2019-3-2-33.574 | UP000002700 |
| 4 | Burkholderia pseudomallei | 159.6542 | 2.2032 | 14898 / 24836 | dbspec-15-Mar-2019-3-2-21.734 | UP000002031 |
| 5 | Burkholderia pseudomallei | 159.5914 | 2.203 | 14898 / 24544 | dbspec-15-Mar-2019-3-2-15.743 | UP000001812 |
| 6 | Burkholderia pseudomallei | 159.4658 | 2.2027 | 14898 / 24630 | dbspec-15-Mar-2019-3-6-4.424 | UP000032603 |
| 7 | Burkholderia pseudomallei 668 | 159.4604 | 2.2027 | 14898 / 24129 | dbspec-15-Mar-2019-3-2-23.122 | UP000002153 |
| 8 | Burkholderia pseudomallei K96243 | 159.2715 | 2.2021 | 14898 / 23961 | dbspec-15-Mar-2019-3-1-58.302 | UP000000605 |
| 9 | Burkholderia pseudomallei | 159.1382 | 2.2018 | 14898 / 24475 | dbspec-15-Mar-2019-3-3-10.094 | UP000004875 |
| 10 | Burkholderia pseudomallei | 159.112 | 2.2017 | 14898 / 24032 | dbspec-15-Mar-2019-3-3-23.479 | UP000005700 |
| 11 | Burkholderia pseudomallei | 158.6596 | 2.2005 | 14898 / 24832 | dbspec-15-Mar-2019-3-5-56.499 | UP000030449 |
| 12 | Burkholderia pseudomallei Pseudomonas pseudomallei | 158.6267 | 2.2004 | 14898 / 24887 | dbspec-15-Mar-2019-3-5-52.272 | UP000029527 |
| 13 | Burkholderia pseudomallei 1106a | 158.5998 | 2.2003 | 14898 / 24105 | dbspec-15-Mar-2019-3-3-42.854 | UP000006738 |
| 14 | Burkholderia pseudomallei 1026b | 158.0287 | 2.1987 | 14898 / 23916 | dbspec-15-Mar-2019-3-4-15.005 | UP000010087 |
| 15 | Burkholderia pseudomallei | 156.7534 | 2.1952 | 14898 / 23842 | dbspec-15-Mar-2019-3-3-0.858 | UP000004448 |
| 16 | Burkholderia pseudomallei | 155.5339 | 2.1918 | 14898 / 22928 | dbspec-15-Mar-2019-3-2-35.82 | UP000002781 |
| 17 | Burkholderia thailandensis | 155.518 | 2.1918 | 14898 / 23446 | dbspec-15-Mar-2019-3-8-58.458 | UP000218714 |
| 18 | Burkholderia pseudomallei | 153.0814 | 2.1849 | 14898 / 21894 | dbspec-15-Mar-2019-3-3-7.02 | UP000004700 |
| 19 | Burkholderia mallei Pseudomonas mallei | 150.0248 | 2.1762 | 14898 / 19381 | dbspec-15-Mar-2019-3-10-40.934 | UP000254505 |
| 20 | Burkholderia mallei | 149.9324 | 2.1759 | 14898 / 19432 | dbspec-15-Mar-2019-3-2-54.15 | UP000003995 |
| 21 | Burkholderia mallei ATCC 23344 | 148.7288 | 2.1724 | 14898 / 18442 | dbspec-15-Mar-2019-3-3-41.84 | UP000006693 |
| 22 | Burkholderia oklahomensis | 147.6645 | 2.1693 | 14898 / 24424 | dbspec-15-Mar-2019-3-10-40.31 | UP000254484 |
| 23 | Burkholderia mallei NCTC 10229 | 147.3197 | 2.1683 | 14898 / 18362 | dbspec-15-Mar-2019-3-2-26.164 | UP000002283 |
| 24 | Burkholderia vietnamiensis G4 / LMG 22486 | 134.8929 | 2.13 | 14898 / 27924 | dbspec-15-Mar-2019-3-2-26.18 | UP000002287 |
| 25 | Arcobacter cryaerophilus | 134.6963 | 2.1294 | 14898 / 6493 | dbspec-15-Mar-2019-3-9-48.44 | UP000239151 |
| 26 | Peptoanaerobacter stomatis | 134.6317 | 2.1291 | 14898 / 6301 | dbspec-15-Mar-2019-3-5-3.755 | UP000017818 |
| 27 | Burkholderia ubonensis | 134.1045 | 2.1274 | 14898 / 27094 | dbspec-15-Mar-2019-3-6-35.016 | UP000058930 |
| 28 | Burkholderia ambifaria | 133.929 | 2.1269 | 14898 / 26412 | dbspec-15-Mar-2019-3-3-20.234 | UP000005463 |
| 29 | Hel cobacter pylori | 133.3524 | 2.125 | 14898 / 6096 | dbspec-15-Mar-2019-3-2-45.352 | UP000003358 |
| 30 | Burkholderia cepacia | 133.0896 | 2.1241 | 14898 / 29972 | dbspec-15-Mar-2019-3-10-52.322 | UP000263019 |

---

#### # 17 - CLASSIFICATION SUBREPORT FOR SPECTRUM *'192_2019_ZBS6_Peter-Burkholderia_DSM13277'*

##### Actual test mass spectrum

| | |
|---|---|
| **genus / species / strain:** | Burkholderia thailandensis DSM 13277 |
| **file id:** | 192_2019_ZBS6_Peter-Burkholderia_DSM13277 |
| **type:** | 192 |
| **NCBI id (species level):** | NaN |
| **NCBI id (strain level):** | NaN |
| **growth time:** | 24 h |
| **growth temperature:** | 37Â°C |
| **growth conditions:** | no CO2 |
| **growth medium:** | Caso (Oxo d) |
| **sample treatment:** | SPEED, operator: A. Schneider |
| **spores:** | no |
| **concentration:** | not available |
| **extra info:** | biological replicate #1, technical replicate #1 |
| **calibration standard:** | not available |
| **measurement method:** | LC-MS1 spectrum, method not available |
| **customer:** | RKI/ZBS 6 |
| **measurement date/time:** | 2019-01_11:11:32.906+02:00 |
| **path to MS file:** | E:\Matlab |

##### List of best matches with test spectrum

| No | Genus/Species/Strain | Score | Log Score | Peak Numbers | Spectral ID | Proteome ID |
|---|---|---|---|---|---|---|
| 1 | Burkholderia thailandensis ATCC 700388 / DSM 13276 / CIP 106301 / E264 | 204.233 | 2.3101 | 16083 / 23226 | dbspec-15-Mar-2019-3-2-18.411 | UP000001930 |
| 2 | Burkholderia thailandensis | 194.5373 | 2.289 | 16083 / 22799 | dbspec-15-Mar-2019-3-9-35.914 | UP000235972 |
| 3 | Burkholderia pseudomallei 668 | 159.5722 | 2.203 | 16083 / 24129 | dbspec-15-Mar-2019-3-2-23.122 | UP000002153 |
| 4 | Burkholderia pseudomallei | 157.7677 | 2.198 | 16083 / 24032 | dbspec-15-Mar-2019-3-3-23.479 | UP000005700 |
| 5 | Burkholderia pseudomallei | 157.4177 | 2.1971 | 16083 / 24475 | dbspec-15-Mar-2019-3-3-10.094 | UP000004875 |
| 6 | Burkholderia pseudomallei Pseudomonas pseudomallei | 157.3655 | 2.1969 | 16083 / 24887 | dbspec-15-Mar-2019-3-5-52.272 | UP000029527 |
| 7 | Burkholderia pseudomallei 1106a | 157.0878 | 2.1961 | 16083 / 24105 | dbspec-15-Mar-2019-3-3-42.854 | UP000006738 |
| 8 | Burkholderia pseudomallei K96243 | 156.9611 | 2.1958 | 16083 / 23961 | dbspec-15-Mar-2019-3-1-58.302 | UP000000605 |
| 9 | Burkholderia pseudomallei | 156.9447 | 2.1957 | 16083 / 24836 | dbspec-15-Mar-2019-3-2-21.734 | UP000002031 |
| 10 | Burkholderia pseudomallei | 156.8812 | 2.1956 | 16083 / 23842 | dbspec-15-Mar-2019-3-3-0.858 | UP000004448 |
| 11 | Burkholderia pseudomallei | 156.6824 | 2.195 | 16083 / 24630 | dbspec-15-Mar-2019-3-6-4.424 | UP000032603 |
| 12 | Burkholderia pseudomallei | 156.6462 | 2.1949 | 16083 / 24544 | dbspec-15-Mar-2019-3-2-15.743 | UP000001812 |
| 13 | Burkholderia pseudomallei 1710b | 156.6437 | 2.1949 | 16083 / 24945 | dbspec-15-Mar-2019-3-2-33.574 | UP000002700 |
| 14 | Burkholderia pseudomallei | 156.5564 | 2.1947 | 16083 / 24832 | dbspec-15-Mar-2019-3-5-56.499 | UP000030449 |
| 15 | Burkholderia pseudomallei 1026b | 155.435 | 2.1915 | 16083 / 23916 | dbspec-15-Mar-2019-3-4-15.005 | UP000010087 |
| 16 | Burkholderia thailandensis | 154.251 | 2.1882 | 16083 / 23446 | dbspec-15-Mar-2019-3-8-58.458 | UP000218714 |
| 17 | Burkholderia pseudomallei | 152.8786 | 2.1843 | 16083 / 22928 | dbspec-15-Mar-2019-3-2-35.82 | UP000002781 |
| 18 | Burkholderia pseudomallei | 151.3633 | 2.18 | 16083 / 21894 | dbspec-15-Mar-2019-3-3-7.02 | UP000004700 |
| 19 | Burkholderia mallei Pseudomonas mallei | 147.0846 | 2.1676 | 16083 / 19381 | dbspec-15-Mar-2019-3-10-40.934 | UP000254505 |

| | | | | | | |
|---|---|---|---|---|---|---|
| 20 | Burkholderia mallei | 146.7675 | 2.1666 | 16083 / 19432 | dbspec-15-Mar-2019-3-2-54.15 | UP000003995 |
| 21 | Burkholderia mallei ATCC 23344 | 145.956 | 2.1642 | 16083 / 18442 | dbspec-15-Mar-2019-3-3-41.84 | UP000006693 |
| 22 | Burkholderia mallei NCTC 10229 | 145.0683 | 2.1616 | 16083 / 18362 | dbspec-15-Mar-2019-3-2-26.164 | UP000002283 |
| 23 | Burkholderia oklahomensis | 143.5785 | 2.1571 | 16083 / 24424 | dbspec-15-Mar-2019-3-10-40.31 | UP000254484 |
| 24 | Hel cobacter cetorum ATCC BAA-540 / MIT 99-5656 | 137.8563 | 2.1394 | 16083 / 6477 | dbspec-15-Mar-2019-3-3-12.153 | UP000005013 |
| 25 | Sanguibacteroides justesenii | 136.6089 | 2.1355 | 16083 / 6412 | dbspec-15-Mar-2019-3-10-13.151 | UP000248308 |
| 26 | Hel cobacter pylori | 136.5444 | 2.1353 | 16083 / 6207 | dbspec-15-Mar-2019-3-3-21.981 | UP000005514 |
| 27 | Streptobacillus moniliformis ATCC 14647 / DSM 12112 / NCTC 10651 / 9901 | 136.0882 | 2.1338 | 16083 / 5488 | dbspec-15-Mar-2019-3-2-22.685 | UP000002072 |
| 28 | Streptobacillus moniliformis | 136.0094 | 2.1336 | 16083 / 5765 | dbspec-15-Mar-2019-3-10-29.921 | UP000251650 |
| 29 | Anaeroglobus geminatus | 135.6606 | 2.1325 | 16083 / 7132 | dbspec-15-Mar-2019-3-3-21.139 | UP000005481 |
| 30 | Campylobacter lanienae | 135.2749 | 2.1312 | 16083 / 6128 | dbspec-15-Mar-2019-3-8-46.571 | UP000202031 |

---

#### # 18 - CLASSIFICATION SUBREPORT FOR SPECTRUM '*193_2019_ZBS6_Peter-Burkholderia_DSM13277*'

##### Actual test mass spectrum

| | |
|---|---|
| **genus / species / strain:** | Burkholderia thailandensis DSM 13277 |
| **file id:** | 193_2019_ZBS6_Peter-Burkholderia_DSM13277 |
| **type:** | 193 |
| **NCBI id (species level):** | NaN |
| **NCBI id (strain level):** | NaN |
| **growth time:** | 24 h |
| **growth temperature:** | 37Â°C |
| **growth conditions:** | no CO2 |
| **growth medium:** | Caso (Oxo d) |
| **sample treatment:** | SPEED, operator: A. Schneider |
| **spores:** | no |
| **concentration:** | not available |
| **extra info:** | biological replicate #1, technical replicate #2 |
| **calibration standard:** | not available |
| **measurement method:** | LC-MS1 spectrum, method not available |
| **customer:** | RKI/ZBS 6 |
| **measurement date/time:** | 2019-01_11:11:32.906+02:00 |
| **path to MS file:** | E:\Matlab |

##### List of best matches with test spectrum

| No | Genus/Species/Strain | Score | Log Score | Peak Numbers | Spectral ID | Proteome ID |
|---|---|---|---|---|---|---|
| 1 | Burkholderia thailandensis ATCC 700388 / DSM 13276 / CIP 106301 / E264 | 206.5838 | 2.3151 | 16252 / 23226 | dbspec-15-Mar-2019-3-2-18.411 | UP000001930 |
| 2 | Burkholderia thailandensis | 197.0168 | 2.2945 | 16252 / 22799 | dbspec-15-Mar-2019-3-9-35.914 | UP000235972 |
| 3 | Burkholderia pseudomallei 668 | 162.9761 | 2.2121 | 16252 / 24129 | dbspec-15-Mar-2019-3-2-23.122 | UP000002153 |
| 4 | Burkholderia pseudomallei 1710b | 162.9752 | 2.2121 | 16252 / 24945 | dbspec-15-Mar-2019-3-2-33.574 | UP000002700 |
| 5 | Burkholderia pseudomallei | 162.7613 | 2.2116 | 16252 / 24630 | dbspec-15-Mar-2019-3-6-4.424 | UP000032603 |
| 6 | Burkholderia pseudomallei | 162.4392 | 2.2107 | 16252 / 24032 | dbspec-15-Mar-2019-3-3-23.479 | UP000005700 |
| 7 | Burkholderia pseudomallei | 162.1925 | 2.21 | 16252 / 24475 | dbspec-15-Mar-2019-3-3-10.094 | UP000004875 |
| 8 | Burkholderia pseudomallei | 162.107 | 2.2098 | 16252 / 24836 | dbspec-15-Mar-2019-3-2-21.734 | UP000002031 |
| 9 | Burkholderia pseudomallei 1106a | 162.0435 | 2.2096 | 16252 / 24105 | dbspec-15-Mar-2019-3-3-42.854 | UP000006738 |
| 10 | Burkholderia pseudomallei Pseudomonas pseudomallei | 161.5658 | 2.2083 | 16252 / 24887 | dbspec-15-Mar-2019-3-5-52.272 | UP000029527 |
| 11 | Burkholderia pseudomallei | 161.4889 | 2.2081 | 16252 / 24544 | dbspec-15-Mar-2019-3-2-15.743 | UP000001812 |
| 12 | Burkholderia pseudomallei | 161.1993 | 2.2074 | 16252 / 24832 | dbspec-15-Mar-2019-3-5-56.499 | UP000030449 |
| 13 | Burkholderia pseudomallei K96243 | 161.183 | 2.2073 | 16252 / 23961 | dbspec-15-Mar-2019-3-1-58.302 | UP000000605 |
| 14 | Burkholderia pseudomallei 1026b | 160.6969 | 2.206 | 16252 / 23916 | dbspec-15-Mar-2019-3-4-15.005 | UP000010087 |
| 15 | Burkholderia pseudomallei | 158.9846 | 2.2014 | 16252 / 23842 | dbspec-15-Mar-2019-3-3-0.858 | UP000004448 |
| 16 | Burkholderia pseudomallei | 157.2971 | 2.1967 | 16252 / 22928 | dbspec-15-Mar-2019-3-2-35.82 | UP000002781 |
| 17 | Burkholderia thailandensis | 156.3602 | 2.1941 | 16252 / 23446 | dbspec-15-Mar-2019-3-8-58.458 | UP000218714 |
| 18 | Burkholderia pseudomallei | 155.0607 | 2.1905 | 16252 / 21894 | dbspec-15-Mar-2019-3-3-7.02 | UP000004700 |
| 19 | Burkholderia mallei Pseudomonas mallei | 151.5592 | 2.1806 | 16252 / 19381 | dbspec-15-Mar-2019-3-10-40.934 | UP000254505 |
| 20 | Burkholderia mallei | 151.4574 | 2.1803 | 16252 / 19432 | dbspec-15-Mar-2019-3-2-54.15 | UP000003995 |
| 21 | Burkholderia mallei ATCC 23344 | 149.9455 | 2.1759 | 16252 / 18442 | dbspec-15-Mar-2019-3-3-41.84 | UP000006693 |
| 22 | Burkholderia mallei NCTC 10229 | 148.693 | 2.1723 | 16252 / 18362 | dbspec-15-Mar-2019-3-2-26.164 | UP000002283 |
| 23 | Burkholderia oklahomensis | 145.2426 | 2.1621 | 16252 / 24424 | dbspec-15-Mar-2019-3-10-40.31 | UP000254484 |
| 24 | Hel cobacter pylori | 139.5674 | 2.1448 | 16252 / 6019 | dbspec-15-Mar-2019-3-4-25.301 | UP000011889 |
| 25 | Hel cobacter pylori | 138.8283 | 2.1425 | 16252 / 6069 | dbspec-15-Mar-2019-3-4-25.333 | UP000011898 |
| 26 | Hel cobacter pylori | 136.9397 | 2.1365 | 16252 / 5958 | dbspec-15-Mar-2019-3-4-26.924 | UP000011953 |
| 27 | Hel cobacter pylori | 136.6475 | 2.1356 | 16252 / 6029 | dbspec-15-Mar-2019-3-3-8.081 | UP000004741 |
| 28 | Hel cobacter pylori | 136.6182 | 2.1355 | 16252 / 5948 | dbspec-15-Mar-2019-3-4-27.205 | UP000012023 |
| 29 | Hel cobacter pylori | 136.5528 | 2.1353 | 16252 / 6207 | dbspec-15-Mar-2019-3-3-21.981 | UP000005514 |
| 30 | Hel cobacter pylori | 136.545 | 2.1353 | 16252 / 6092 | dbspec-15-Mar-2019-3-2-47.942 | UP000003606 |

---

#### # 19 - CLASSIFICATION SUBREPORT FOR SPECTRUM '*194_2019_ZBS6_Peter-Burkholderia_DSM13277*'

##### Actual test mass spectrum

| | |
|---|---|
| **genus / species / strain:** | Burkholderia thailandensis DSM 13277 |

| | |
|---|---|
| **file id:** | 194_2019_ZBS6_Peter-Burkholderia_DSM13277 |
| **type:** | 194 |
| **NCBI id (species level):** | NaN |
| **NCBI id (strain level):** | NaN |
| **growth time:** | 24 h |
| **growth temperature:** | 37Â°C |
| **growth conditions:** | no CO2 |
| **growth medium:** | Caso (Oxo d) |
| **sample treatment:** | SPEED, operator: A. Schneider |
| **spores:** | no |
| **concentration:** | not available |
| **extra info:** | biological replicate #1, technical replicate #3 |
| **calibration standard:** | not available |
| **measurement method:** | LC-MS1 spectrum, method not available |
| **customer:** | RKI/ZBS 6 |
| **measurement date/time:** | 2019-01_11:11:32.906+02:00 |
| **path to MS file:** | E:\Matlab |

#### List of best matches with test spectrum

| No | Genus/Species/Strain | Score | Log Score | Peak Numbers | Spectral ID | Proteome ID |
|---|---|---|---|---|---|---|
| 1 | Burkholderia thailandensis ATCC 700388 / DSM 13276 / CIP 106301 / E264 | 208.7549 | 2.3196 | 16228 / 23226 | dbspec-15-Mar-2019-3-2-18.411 | UP000001930 |
| 2 | Burkholderia thailandensis | 196.7919 | 2.294 | 16228 / 22799 | dbspec-15-Mar-2019-3-9-35.914 | UP000235972 |
| 3 | Burkholderia pseudomallei 1710b | 162.134 | 2.2099 | 16228 / 24945 | dbspec-15-Mar-2019-3-2-33.574 | UP000002700 |
| 4 | Burkholderia pseudomallei | 161.5117 | 2.2082 | 16228 / 24630 | dbspec-15-Mar-2019-3-6-4.424 | UP000032603 |
| 5 | Burkholderia pseudomallei | 161.4378 | 2.208 | 16228 / 24544 | dbspec-15-Mar-2019-3-2-15.743 | UP000001812 |
| 6 | Burkholderia pseudomallei | 161.3301 | 2.2077 | 16228 / 24475 | dbspec-15-Mar-2019-3-3-10.094 | UP000004875 |
| 7 | Burkholderia pseudomallei | 161.1196 | 2.2071 | 16228 / 24032 | dbspec-15-Mar-2019-3-3-23.479 | UP000005700 |
| 8 | Burkholderia pseudomallei 668 | 161.0983 | 2.2071 | 16228 / 24129 | dbspec-15-Mar-2019-3-2-23.122 | UP000002153 |
| 9 | Burkholderia pseudomallei | 161.0847 | 2.2071 | 16228 / 24836 | dbspec-15-Mar-2019-3-2-21.734 | UP000002031 |
| 10 | Burkholderia pseudomallei 1106a | 161.0011 | 2.2068 | 16228 / 24105 | dbspec-15-Mar-2019-3-3-42.854 | UP000006738 |
| 11 | Burkholderia pseudomallei | 160.5606 | 2.2056 | 16228 / 24832 | dbspec-15-Mar-2019-3-5-56.499 | UP000030449 |
| 12 | Burkholderia pseudomallei K96243 | 160.3106 | 2.205 | 16228 / 23961 | dbspec-15-Mar-2019-3-1-58.302 | UP000000605 |
| 13 | Burkholderia pseudomallei Pseudomonas pseudomallei | 159.6884 | 2.2033 | 16228 / 24887 | dbspec-15-Mar-2019-3-5-52.272 | UP000029527 |
| 14 | Burkholderia pseudomallei | 159.4953 | 2.2027 | 16228 / 23842 | dbspec-15-Mar-2019-3-3-0.858 | UP000004448 |
| 15 | Burkholderia pseudomallei 1026b | 159.1369 | 2.2018 | 16228 / 23916 | dbspec-15-Mar-2019-3-4-15.005 | UP000010087 |
| 16 | Burkholderia pseudomallei | 157.2971 | 2.1967 | 16228 / 22928 | dbspec-15-Mar-2019-3-2-35.82 | UP000002781 |
| 17 | Burkholderia thailandensis | 155.6825 | 2.1922 | 16228 / 23446 | dbspec-15-Mar-2019-3-8-58.458 | UP000218714 |
| 18 | Burkholderia pseudomallei | 153.6592 | 2.1866 | 16228 / 21894 | dbspec-15-Mar-2019-3-3-7.02 | UP000004700 |
| 19 | Burkholderia mallei | 151.2884 | 2.1798 | 16228 / 19432 | dbspec-15-Mar-2019-3-2-54.15 | UP000003995 |
| 20 | Burkholderia mallei Pseudomonas mallei | 151.2038 | 2.1796 | 16228 / 19381 | dbspec-15-Mar-2019-3-10-40.934 | UP000254505 |
| 21 | Burkholderia mallei ATCC 23344 | 150.1283 | 2.1765 | 16228 / 18442 | dbspec-15-Mar-2019-3-3-41.84 | UP000006693 |
| 22 | Burkholderia mallei NCTC 10229 | 148.1723 | 2.1708 | 16228 / 18362 | dbspec-15-Mar-2019-3-2-26.164 | UP000002283 |
| 23 | Burkholderia oklahomensis | 142.8783 | 2.155 | 16228 / 24424 | dbspec-15-Mar-2019-3-10-40.31 | UP000254484 |
| 24 | Prevotella intermedia | 136.5025 | 2.1351 | 16228 / 7894 | dbspec-15-Mar-2019-3-9-23.777 | UP000231201 |
| 25 | Riemerella anatipestifer ATCC 11845 / DSM 15868 / JCM 9532 / NCTC 11014 | 135.78 | 2.1328 | 16228 / 8046 | dbspec-15-Mar-2019-3-4-15.021 | UP000010093 |
| 26 | Peptoniphilus lacrimalis | 135.6934 | 2.1326 | 16228 / 5889 | dbspec-15-Mar-2019-3-3-23.915 | UP000005711 |
| 27 | Porphyromonas crevioricanis | 135.6393 | 2.1324 | 16228 / 8115 | dbspec-15-Mar-2019-3-5-5.315 | UP000018031 |
| 28 | Arcobacter skirrowii | 135.3084 | 2.1313 | 16228 / 7091 | dbspec-15-Mar-2019-3-10-3.931 | UP000245014 |
| 29 | Hel cobacter pylori ATCC 700392 / 26695 | 134.5769 | 2.129 | 16228 / 6120 | dbspec-15-Mar-2019-3-1-54.87 | UP000000429 |
| 30 | Prevotella intermedia | 134.4708 | 2.1286 | 16228 / 8092 | dbspec-15-Mar-2019-3-9-23.09 | UP000230919 |

---

#### # 20 - CLASSIFICATION SUBREPORT FOR SPECTRUM *'248_2019_ZBS6_Peter-Burkholderia_DSM13277'*

##### Actual test mass spectrum

| | |
|---|---|
| **genus / species / strain:** | Burkholderia thailandensis DSM 13277 |
| **file id:** | 248_2019_ZBS6_Peter-Burkholderia_DSM13277 |
| **type:** | 248 |
| **NCBI id (species level):** | NaN |
| **NCBI id (strain level):** | NaN |
| **growth time:** | 24 h |
| **growth temperature:** | 37Â°C |
| **growth conditions:** | no CO2 |
| **growth medium:** | Caso (Oxo d) |
| **sample treatment:** | SPEED, operator: A. Schneider |
| **spores:** | no |
| **concentration:** | not available |
| **extra info:** | biological replicate #2, technical replicate #1 |
| **calibration standard:** | not available |
| **measurement method:** | LC-MS1 spectrum, method not available |
| **customer:** | RKI/ZBS 6 |
| **measurement date/time:** | 2019-01_11:11:32.906+02:00 |
| **path to MS file:** | E:\Matlab |

#### List of best matches with test spectrum

| No | Genus/Species/Strain | Score | Log Score | Peak Numbers | Spectral ID | Proteome ID |
|---|---|---|---|---|---|---|
| 1 | Burkholderia thailandensis ATCC 700388 / DSM 13276 / CIP 106301 / E264 | 192.1228 | 2.2836 | 15144 / 23226 | dbspec-15-Mar-2019-3-2-18.411 | UP000001930 |
| 2 | Burkholderia thailandensis | 183.0681 | 2.2626 | 15144 / 22799 | dbspec-15-Mar-2019-3-9-35.914 | UP000235972 |
| 3 | Burkholderia pseudomallei Pseudomonas pseudomallei | 153.6698 | 2.1866 | 15144 / 24887 | dbspec-15-Mar-2019-3-5-52.272 | UP000029527 |
| 4 | Burkholderia pseudomallei 668 | 152.2834 | 2.1827 | 15144 / 24129 | dbspec-15-Mar-2019-3-2-23.122 | UP000002153 |
| 5 | Burkholderia pseudomallei | 152.0424 | 2.182 | 15144 / 24630 | dbspec-15-Mar-2019-3-6-4.424 | UP000032603 |
| 6 | Burkholderia pseudomallei 1710b | 151.9586 | 2.1817 | 15144 / 24945 | dbspec-15-Mar-2019-3-2-33.574 | UP000002700 |
| 7 | Burkholderia pseudomallei | 151.1369 | 2.1794 | 15144 / 24832 | dbspec-15-Mar-2019-3-5-56.499 | UP000030449 |
| 8 | Burkholderia pseudomallei 1026b | 151.0658 | 2.1792 | 15144 / 23916 | dbspec-15-Mar-2019-3-4-15.005 | UP000010087 |
| 9 | Burkholderia pseudomallei | 151.0434 | 2.1791 | 15144 / 24032 | dbspec-15-Mar-2019-3-3-23.479 | UP000005700 |
| 10 | Burkholderia pseudomallei | 151.0416 | 2.1791 | 15144 / 24544 | dbspec-15-Mar-2019-3-2-15.743 | UP000001812 |
| 11 | Burkholderia pseudomallei K96243 | 150.9247 | 2.1788 | 15144 / 23961 | dbspec-15-Mar-2019-3-1-58.302 | UP000000605 |
| 12 | Burkholderia pseudomallei | 150.8875 | 2.1787 | 15144 / 24836 | dbspec-15-Mar-2019-3-2-21.734 | UP000002031 |
| 13 | Burkholderia pseudomallei 1106a | 150.6078 | 2.1778 | 15144 / 24105 | dbspec-15-Mar-2019-3-3-42.854 | UP000006738 |
| 14 | Burkholderia pseudomallei | 149.6493 | 2.1751 | 15144 / 24475 | dbspec-15-Mar-2019-3-3-10.094 | UP000004875 |
| 15 | Burkholderia thailandensis | 149.0666 | 2.1734 | 15144 / 23446 | dbspec-15-Mar-2019-3-8-58.458 | UP000218714 |
| 16 | Burkholderia pseudomallei | 148.622 | 2.1721 | 15144 / 22928 | dbspec-15-Mar-2019-3-2-35.82 | UP000002781 |
| 17 | Burkholderia pseudomallei | 148.5115 | 2.1718 | 15144 / 23842 | dbspec-15-Mar-2019-3-3-0.858 | UP000004448 |
| 18 | Burkholderia pseudomallei | 147.3077 | 2.1682 | 15144 / 21894 | dbspec-15-Mar-2019-3-3-7.02 | UP000004700 |
| 19 | Burkholderia mallei | 144.5132 | 2.1599 | 15144 / 19432 | dbspec-15-Mar-2019-3-2-54.15 | UP000003995 |
| 20 | Burkholderia mallei Pseudomonas mallei | 144.1572 | 2.1588 | 15144 / 19381 | dbspec-15-Mar-2019-3-10-40.934 | UP000254505 |
| 21 | Burkholderia oklahomensis | 142.6922 | 2.1544 | 15144 / 24424 | dbspec-15-Mar-2019-3-10-40.31 | UP000254484 |
| 22 | Burkholderia mallei ATCC 23344 | 142.569 | 2.154 | 15144 / 18442 | dbspec-15-Mar-2019-3-3-41.84 | UP000006693 |
| 23 | Burkholderia mallei NCTC 10229 | 142.4639 | 2.1537 | 15144 / 18362 | dbspec-15-Mar-2019-3-2-26.164 | UP000002283 |
| 24 | Burkholderia vietnamiensis G4 / LMG 22486 | 132.5274 | 2.1223 | 15144 / 27924 | dbspec-15-Mar-2019-3-2-26.18 | UP000002287 |
| 25 | Arcobacter cryaerophilus | 132.5178 | 2.1223 | 15144 / 6493 | dbspec-15-Mar-2019-3-9-48.44 | UP000239151 |
| 26 | Leptotr chia wadei F0279 | 131.485 | 2.1189 | 15144 / 7438 | dbspec-15-Mar-2019-3-4-56.907 | UP000016626 |
| 27 | Burkholderia cepacia Pseudomonas cepacia | 130.3843 | 2.1152 | 15144 / 29003 | dbspec-15-Mar-2019-3-5-52.038 | UP000029423 |
| 28 | Burkholderia ambifaria | 129.5231 | 2.1123 | 15144 / 26412 | dbspec-15-Mar-2019-3-3-20.234 | UP000005463 |
| 29 | Burkholderia reimsis | 129.1765 | 2.1112 | 15144 / 30986 | dbspec-15-Mar-2019-3-10-32.495 | UP000252458 |
| 30 | Neisseria mening tidis | 128.824 | 2.11 | 15144 / 7747 | dbspec-15-Mar-2019-3-7-26.433 | UP000182715 |

### # 21 - CLASSIFICATION SUBREPORT FOR SPECTRUM *'168_2019_ZBS6_Peter-Burkholderia_21774'*

#### Actual test mass spectrum

| | |
|---|---|
| **genus / species / strain:** | Burkholderia oklahomensis DSM 21774 |
| **file id:** | 168_2019_ZBS6_Peter-Burkholderia_21774 |
| **type:** | 168 |
| **NCBI id (species level):** | NaN |
| **NCBI id (strain level):** | NaN |
| **growth time:** | 24 h |
| **growth temperature:** | 37Â°C |
| **growth conditions:** | no CO2 |
| **growth medium:** | Caso (Oxo d) |
| **sample treatment:** | SPEED, operator: A. Schneider |
| **spores:** | no |
| **concentration:** | not available |
| **extra info:** | biological replicate #1, technical replicate #1 |
| **calibration standard:** | not available |
| **measurement method:** | LC-MS1 spectrum, method not available |
| **customer:** | RKI/ZBS 6 |
| **measurement date/time:** | 2019-01_11:11:32.906+02:00 |
| **path to MS file:** | E:\Matlab |

#### List of best matches with test spectrum

| No | Genus/Species/Strain | Score | Log Score | Peak Numbers | Spectral ID | Proteome ID |
|---|---|---|---|---|---|---|
| 1 | Burkholderia oklahomensis | 217.199 | 2.3369 | 16485 / 24424 | dbspec-15-Mar-2019-3-10-40.31 | UP000254484 |
| 2 | Burkholderia pseudomallei | 149.3277 | 2.1741 | 16485 / 24836 | dbspec-15-Mar-2019-3-2-21.734 | UP000002031 |
| 3 | Burkholderia pseudomallei 1710b | 149.2699 | 2.174 | 16485 / 24945 | dbspec-15-Mar-2019-3-2-33.574 | UP000002700 |
| 4 | Burkholderia pseudomallei Pseudomonas pseudomallei | 148.7986 | 2.1726 | 16485 / 24887 | dbspec-15-Mar-2019-3-5-52.272 | UP000029527 |
| 5 | Burkholderia pseudomallei | 148.5615 | 2.1719 | 16485 / 24630 | dbspec-15-Mar-2019-3-6-4.424 | UP000032603 |
| 6 | Burkholderia pseudomallei | 148.409 | 2.1715 | 16485 / 24544 | dbspec-15-Mar-2019-3-2-15.743 | UP000001812 |
| 7 | Burkholderia pseudomallei K96243 | 147.7105 | 2.1694 | 16485 / 23961 | dbspec-15-Mar-2019-3-1-58.302 | UP000000605 |
| 8 | Burkholderia pseudomallei 668 | 147.4779 | 2.1687 | 16485 / 24129 | dbspec-15-Mar-2019-3-2-23.122 | UP000002153 |
| 9 | Burkholderia pseudomallei | 147.1982 | 2.1679 | 16485 / 24475 | dbspec-15-Mar-2019-3-3-10.094 | UP000004875 |
| 10 | Burkholderia thailandensis | 146.9722 | 2.1672 | 16485 / 23446 | dbspec-15-Mar-2019-3-8-58.458 | UP000218714 |
| 11 | Burkholderia pseudomallei 1026b | 146.6508 | 2.1663 | 16485 / 23916 | dbspec-15-Mar-2019-3-4-15.005 | UP000010087 |
| 12 | Burkholderia pseudomallei | 146.5943 | 2.1661 | 16485 / 24832 | dbspec-15-Mar-2019-3-5-56.499 | UP000030449 |
| 13 | Burkholderia pseudomallei | 146.5322 | 2.1659 | 16485 / 24032 | dbspec-15-Mar-2019-3-3-23.479 | UP000005700 |
| 14 | Burkholderia pseudomallei 1106a | 146.098 | 2.1646 | 16485 / 24105 | dbspec-15-Mar-2019-3-3-42.854 | UP000006738 |
| 15 | Burkholderia pseudomallei | 145.5062 | 2.1629 | 16485 / 23842 | dbspec-15-Mar-2019-3-3-0.858 | UP000004448 |
| 16 | Burkholderia thailandensis | 145.2809 | 2.1622 | 16485 / 22799 | dbspec-15-Mar-2019-3-9-35.914 | UP000235972 |
| 17 | Burkholderia thailandensis ATCC 700388 / DSM 13276 / CIP 106301 / E264 | 145.1429 | 2.1618 | 16485 / 23226 | dbspec-15-Mar-2019-3-2-18.411 | UP000001930 |

| | | | | | | |
|---|---|---|---|---|---|---|
| 18 | Burkholderia pseudomallei | 144.8626 | 2.161 | 16485 / 22928 | dbspec-15-Mar-2019-3-2-35.82 | UP000002781 |
| 19 | Burkholderia mallei Pseudomonas mallei | 140.6895 | 2.1483 | 16485 / 19381 | dbspec-15-Mar-2019-3-10-40.934 | UP000254505 |
| 20 | Burkholderia mallei | 140.3507 | 2.1472 | 16485 / 19432 | dbspec-15-Mar-2019-3-2-54.15 | UP000003995 |
| 21 | Burkholderia mallei ATCC 23344 | 139.6644 | 2.1451 | 16485 / 18442 | dbspec-15-Mar-2019-3-3-41.84 | UP000006693 |
| 22 | Burkholderia mallei NCTC 10229 | 139.6644 | 2.1451 | 16485 / 18362 | dbspec-15-Mar-2019-3-2-26.164 | UP000002283 |
| 23 | Burkholderia pseudomallei | 139.6511 | 2.145 | 16485 / 21894 | dbspec-15-Mar-2019-3-3-7.02 | UP000004700 |
| 24 | Hel cobacter saguini | 137.6687 | 2.1388 | 16485 / 7111 | dbspec-15-Mar-2019-3-5-54.284 | UP000029714 |
| 25 | Riemerella anatipestifer ATCC 11845 / DSM 15868 / JCM 9532 / NCTC 11014 | 137.6288 | 2.1387 | 16485 / 8046 | dbspec-15-Mar-2019-3-4-15.021 | UP000010093 |
| 26 | Sanguibacteroides justesenii | 137.1762 | 2.1373 | 16485 / 6412 | dbspec-15-Mar-2019-3-10-13.151 | UP000248308 |
| 27 | Prevotella intermedia | 136.7565 | 2.1359 | 16485 / 8428 | dbspec-15-Mar-2019-3-9-19.955 | UP000229323 |
| 28 | Hel cobacter pylori | 135.6116 | 2.1323 | 16485 / 6086 | dbspec-15-Mar-2019-3-3-11.919 | UP000004970 |
| 29 | Prevotella intermedia | 135.489 | 2.1319 | 16485 / 8585 | dbspec-15-Mar-2019-3-9-18.535 | UP000228641 |
| 30 | Tenacibaculum dicentrarchi | 135.4754 | 2.1319 | 16485 / 8112 | dbspec-15-Mar-2019-3-9-29.97 | UP000234582 |

#### # 22 - CLASSIFICATION SUBREPORT FOR SPECTRUM '*172_2019_ZBS6_Peter-Burkholderia_21774*'

##### Actual test mass spectrum

| | |
|---|---|
| **genus / species / strain:** | Burkholderia oklahomensis DSM 21774 |
| **file id:** | 172_2019_ZBS6_Peter-Burkholderia_21774 |
| **type:** | 172 |
| **NCBI id (species level):** | NaN |
| **NCBI id (strain level):** | NaN |
| **growth time:** | 24 h |
| **growth temperature:** | 37Â°C |
| **growth conditions:** | no CO2 |
| **growth medium:** | Caso (Oxo d) |
| **sample treatment:** | SPEED, operator: A. Schneider |
| **spores:** | no |
| **concentration:** | not available |
| **extra info:** | biological replicate #1, technical replicate #2 |
| **calibration standard:** | not available |
| **measurement method:** | LC-MS1 spectrum, method not available |
| **customer:** | RKI/ZBS 6 |
| **measurement date/time:** | 2019-01_11:11:32.906+02:00 |
| **path to MS file:** | E:\Matlab |

##### List of best matches with test spectrum

| No | Genus/Species/Strain | Score | Log Score | Peak Numbers | Spectral ID | Proteome ID |
|---|---|---|---|---|---|---|
| 1 | Burkholderia oklahomensis | 211.9611 | 2.3263 | 15944 / 24424 | dbspec-15-Mar-2019-3-10-40.31 | UP000254484 |
| 2 | Burkholderia pseudomallei 1710b | 146.1807 | 2.1649 | 15944 / 24945 | dbspec-15-Mar-2019-3-2-33.574 | UP000002700 |
| 3 | Burkholderia pseudomallei | 145.7201 | 2.1635 | 15944 / 24836 | dbspec-15-Mar-2019-3-2-21.734 | UP000002031 |
| 4 | Burkholderia pseudomallei | 145.6309 | 2.1633 | 15944 / 24544 | dbspec-15-Mar-2019-3-2-15.743 | UP000001812 |
| 5 | Burkholderia pseudomallei K96243 | 145.4409 | 2.1627 | 15944 / 23961 | dbspec-15-Mar-2019-3-1-58.302 | UP000000605 |
| 6 | Burkholderia thailandensis ATCC 700388 / DSM 13276 / CIP 106301 / E264 | 145.1695 | 2.1619 | 15944 / 23226 | dbspec-15-Mar-2019-3-2-18.411 | UP000001930 |
| 7 | Burkholderia pseudomallei Pseudomonas pseudomallei | 144.9381 | 2.1612 | 15944 / 24887 | dbspec-15-Mar-2019-3-5-52.272 | UP000029527 |
| 8 | Burkholderia pseudomallei | 144.7885 | 2.1607 | 15944 / 24475 | dbspec-15-Mar-2019-3-3-10.094 | UP000004875 |
| 9 | Burkholderia pseudomallei | 144.7511 | 2.1606 | 15944 / 24832 | dbspec-15-Mar-2019-3-5-56.499 | UP000030449 |
| 10 | Burkholderia pseudomallei | 144.717 | 2.1605 | 15944 / 24630 | dbspec-15-Mar-2019-3-6-4.424 | UP000032603 |
| 11 | Burkholderia pseudomallei 668 | 144.65 | 2.1603 | 15944 / 24129 | dbspec-15-Mar-2019-3-2-23.122 | UP000002153 |
| 12 | Burkholderia thailandensis | 144.5546 | 2.16 | 15944 / 23446 | dbspec-15-Mar-2019-3-8-58.458 | UP000218714 |
| 13 | Burkholderia thailandensis | 144.2107 | 2.159 | 15944 / 22799 | dbspec-15-Mar-2019-3-9-35.914 | UP000235972 |
| 14 | Burkholderia pseudomallei | 143.9774 | 2.1583 | 15944 / 24032 | dbspec-15-Mar-2019-3-3-23.479 | UP000005700 |
| 15 | Burkholderia pseudomallei 1026b | 142.9271 | 2.1551 | 15944 / 23916 | dbspec-15-Mar-2019-3-4-15.005 | UP000010087 |
| 16 | Burkholderia pseudomallei 1106a | 142.8955 | 2.155 | 15944 / 24105 | dbspec-15-Mar-2019-3-3-42.854 | UP000006738 |
| 17 | Burkholderia pseudomallei | 142.7957 | 2.1547 | 15944 / 23842 | dbspec-15-Mar-2019-3-3-0.858 | UP000004448 |
| 18 | Burkholderia pseudomallei | 141.3192 | 2.1502 | 15944 / 22928 | dbspec-15-Mar-2019-3-2-35.82 | UP000002781 |
| 19 | Burkholderia pseudomallei | 137.224 | 2.1374 | 15944 / 21894 | dbspec-15-Mar-2019-3-3-7.02 | UP000004700 |
| 20 | Burkholderia mallei Pseudomonas mallei | 135.9329 | 2.1333 | 15944 / 19381 | dbspec-15-Mar-2019-3-10-40.934 | UP000254505 |
| 21 | Burkholderia mallei | 135.8463 | 2.133 | 15944 / 19432 | dbspec-15-Mar-2019-3-2-54.15 | UP000003995 |
| 22 | Burkholderia mallei NCTC 10229 | 135.7731 | 2.1328 | 15944 / 18362 | dbspec-15-Mar-2019-3-2-26.164 | UP000002283 |
| 23 | Burkholderia mallei ATCC 23344 | 135.4712 | 2.1318 | 15944 / 18442 | dbspec-15-Mar-2019-3-3-41.84 | UP000006693 |
| 24 | Capnocytophaga canis | 133.9701 | 2.127 | 15944 / 7708 | dbspec-15-Mar-2019-3-6-21.35 | UP000045051 |
| 25 | Sanguibacteroides justesenii | 133.4142 | 2.1252 | 15944 / 6412 | dbspec-15-Mar-2019-3-10-13.151 | UP000248308 |
| 26 | Leptotr chia wadei F0279 | 132.571 | 2.1224 | 15944 / 7438 | dbspec-15-Mar-2019-3-4-56.907 | UP000016626 |
| 27 | Dyadobacter jiangsuensis | 132.1852 | 2.1212 | 15944 / 31028 | dbspec-15-Mar-2019-3-9-56.131 | UP000241964 |
| 28 | Campylobacter upsaliensis | 131.7728 | 2.1198 | 15944 / 6179 | dbspec-15-Mar-2019-3-3-26.443 | UP000005813 |
| 29 | Burkholderia reimsis | 131.408 | 2.1186 | 15944 / 30986 | dbspec-15-Mar-2019-3-10-32.495 | UP000252458 |
| 30 | Desulfurobacterium atlanticum | 130.6981 | 2.1163 | 15944 / 6084 | dbspec-15-Mar-2019-3-8-23.451 | UP000198405 |

#### # 23 - CLASSIFICATION SUBREPORT FOR SPECTRUM '*176_2019_ZBS6_Peter-Burkholderia_21774*'

##### Actual test mass spectrum

| | |
|---|---|
| **genus / species / strain:** | Burkholderia oklahomensis DSM 21774 |
| **file id:** | 176_2019_ZBS6_Peter-Burkholderia_21774 |
| **type:** | 176 |
| **NCBI id (species level):** | NaN |
| **NCBI id (strain level):** | NaN |
| **growth time:** | 24 h |
| **growth temperature:** | 37Â°C |
| **growth conditions:** | no CO2 |
| **growth medium:** | Caso (Oxo d) |
| **sample treatment:** | SPEED, operator: A. Schneider |
| **spores:** | no |
| **concentration:** | not available |
| **extra info:** | biological replicate #1, technical replicate #3 |
| **calibration standard:** | not available |
| **measurement method:** | LC-MS1 spectrum, method not available |
| **customer:** | RKI/ZBS 6 |
| **measurement date/time:** | 2019-01_11:11:32.906+02:00 |
| **path to MS file:** | E:\Matlab |

#### List of best matches with test spectrum

| No | Genus/Species/Strain | Score | Log Score | Peak Numbers | Spectral ID | Proteome ID |
|---|---|---|---|---|---|---|
| 1 | Burkholderia oklahomensis | 207.9029 | 2.3179 | 15640 / 24424 | dbspec-15-Mar-2019-3-10-40.31 | UP000254484 |
| 2 | Burkholderia pseudomallei | 147.1922 | 2.1679 | 15640 / 24836 | dbspec-15-Mar-2019-3-2-21.734 | UP000002031 |
| 3 | Burkholderia pseudomallei 1710b | 147.169 | 2.1678 | 15640 / 24945 | dbspec-15-Mar-2019-3-2-33.574 | UP000002700 |
| 4 | Burkholderia pseudomallei 668 | 147.102 | 2.1676 | 15640 / 24129 | dbspec-15-Mar-2019-3-2-23.122 | UP000002153 |
| 5 | Burkholderia pseudomallei Pseudomonas pseudomallei | 145.6726 | 2.1634 | 15640 / 24887 | dbspec-15-Mar-2019-3-5-52.272 | UP000029527 |
| 6 | Burkholderia pseudomallei | 145.6538 | 2.1633 | 15640 / 24544 | dbspec-15-Mar-2019-3-2-15.743 | UP000001812 |
| 7 | Burkholderia pseudomallei K96243 | 145.5045 | 2.1629 | 15640 / 23961 | dbspec-15-Mar-2019-3-1-58.302 | UP000000605 |
| 8 | Burkholderia pseudomallei | 145.4357 | 2.1627 | 15640 / 24630 | dbspec-15-Mar-2019-3-6-4.424 | UP000032603 |
| 9 | Burkholderia pseudomallei | 145.0273 | 2.1614 | 15640 / 24832 | dbspec-15-Mar-2019-3-5-56.499 | UP000030449 |
| 10 | Burkholderia pseudomallei 1026b | 144.9303 | 2.1612 | 15640 / 23916 | dbspec-15-Mar-2019-3-4-15.005 | UP000010087 |
| 11 | Burkholderia pseudomallei | 144.5154 | 2.1599 | 15640 / 24475 | dbspec-15-Mar-2019-3-3-10.094 | UP000004875 |
| 12 | Burkholderia pseudomallei | 144.3639 | 2.1595 | 15640 / 24032 | dbspec-15-Mar-2019-3-3-23.479 | UP000005700 |
| 13 | Burkholderia pseudomallei 1106a | 144.2215 | 2.159 | 15640 / 24105 | dbspec-15-Mar-2019-3-3-42.854 | UP000006738 |
| 14 | Burkholderia thailandensis | 143.0219 | 2.1554 | 15640 / 22799 | dbspec-15-Mar-2019-3-9-35.914 | UP000235972 |
| 15 | Burkholderia pseudomallei | 142.3921 | 2.1535 | 15640 / 23842 | dbspec-15-Mar-2019-3-3-0.858 | UP000004448 |
| 16 | Burkholderia thailandensis ATCC 700388 / DSM 13276 / CIP 106301 / E264 | 141.2481 | 2.15 | 15640 / 23226 | dbspec-15-Mar-2019-3-2-18.411 | UP000001930 |
| 17 | Burkholderia thailandensis | 141.066 | 2.1494 | 15640 / 23446 | dbspec-15-Mar-2019-3-8-58.458 | UP000218714 |
| 18 | Burkholderia pseudomallei | 140.8678 | 2.1488 | 15640 / 22928 | dbspec-15-Mar-2019-3-2-35.82 | UP000002781 |
| 19 | Burkholderia pseudomallei | 138.058 | 2.1401 | 15640 / 21894 | dbspec-15-Mar-2019-3-3-7.02 | UP000004700 |
| 20 | Burkholderia mallei Pseudomonas mallei | 137.6132 | 2.1387 | 15640 / 19381 | dbspec-15-Mar-2019-3-10-40.934 | UP000254505 |
| 21 | Burkholderia mallei ATCC 23344 | 137.5372 | 2.1384 | 15640 / 18442 | dbspec-15-Mar-2019-3-3-41.84 | UP000006693 |
| 22 | Burkholderia mallei | 137.2077 | 2.1374 | 15640 / 19432 | dbspec-15-Mar-2019-3-2-54.15 | UP000003995 |
| 23 | Burkholderia mallei NCTC 10229 | 136.7949 | 2.1361 | 15640 / 18362 | dbspec-15-Mar-2019-3-2-26.164 | UP000002283 |
| 24 | Riemerella anatipestifer ATCC 11845 / DSM 15868 / JCM 9532 / NCTC 11014 | 130.8535 | 2.1168 | 15640 / 8046 | dbspec-15-Mar-2019-3-4-15.021 | UP000010093 |
| 25 | Burkholderia pyrrocinia Pseudomonas pyrrocinia | 129.7318 | 2.113 | 15640 / 28460 | dbspec-15-Mar-2019-3-6-38.026 | UP000064483 |
| 26 | Burkholderia reimsis | 129.4645 | 2.1122 | 15640 / 30986 | dbspec-15-Mar-2019-3-10-32.495 | UP000252458 |
| 27 | Chitinophaga ginsengisoli | 129.3023 | 2.1116 | 15640 / 31964 | dbspec-15-Mar-2019-3-9-53.947 | UP000240978 |
| 28 | Burkholderia ubonensis | 128.9963 | 2.1106 | 15640 / 27094 | dbspec-15-Mar-2019-3-6-35.016 | UP000058930 |
| 29 | Burkholderia cepacia | 128.9767 | 2.1105 | 15640 / 29972 | dbspec-15-Mar-2019-3-10-52.322 | UP000263019 |
| 30 | Burkholderia cepacia Pseudomonas cepacia | 128.0182 | 2.1073 | 15640 / 30555 | dbspec-15-Mar-2019-3-6-40.382 | UP000068482 |

---

### # 24 - CLASSIFICATION SUBREPORT FOR SPECTRUM *'243_2019_ZBS6_Peter-Burkholderia_DSM21774'*

#### Actual test mass spectrum

| | |
|---|---|
| **genus / species / strain:** | Burkholderia oklahomensis DSM 21774 |
| **file id:** | 243_2019_ZBS6_Peter-Burkholderia_DSM21774 |
| **type:** | 243 |
| **NCBI id (species level):** | NaN |
| **NCBI id (strain level):** | NaN |
| **growth time:** | 24 h |
| **growth temperature:** | 37Â°C |
| **growth conditions:** | no CO2 |
| **growth medium:** | Caso (Oxo d) |
| **sample treatment:** | SPEED, operator: A. Schneider |
| **spores:** | no |
| **concentration:** | not available |
| **extra info:** | biological replicate #2, technical replicate #1 |
| **calibration standard:** | not available |
| **measurement method:** | LC-MS1 spectrum, method not available |
| **customer:** | RKI/ZBS 6 |
| **measurement date/time:** | 2019-01_11:11:32.906+02:00 |
| **path to MS file:** | E:\Matlab |

#### List of best matches with test spectrum

| No | Genus/Species/Strain | Score | Log Score | Peak Numbers | Spectral ID | Proteome ID |
|---|---|---|---|---|---|---|
| 1 | Burkholderia oklahomensis | 202.1129 | 2.3056 | 14928 / 24424 | dbspec-15-Mar-2019-3-10-40.31 | UP000254484 |
| 2 | Burkholderia pseudomallei 1710b | 148.8684 | 2.1728 | 14928 / 24945 | dbspec-15-Mar-2019-3-2-33.574 | UP000002700 |
| 3 | Burkholderia pseudomallei 668 | 147.8435 | 2.1698 | 14928 / 24129 | dbspec-15-Mar-2019-3-2-23.122 | UP000002153 |
| 4 | Burkholderia pseudomallei | 147.2967 | 2.1682 | 14928 / 24544 | dbspec-15-Mar-2019-3-2-15.743 | UP000001812 |
| 5 | Burkholderia pseudomallei | 146.6417 | 2.1663 | 14928 / 24836 | dbspec-15-Mar-2019-3-2-21.734 | UP000002031 |
| 6 | Burkholderia pseudomallei Pseudomonas pseudomallei | 145.7389 | 2.1636 | 14928 / 24887 | dbspec-15-Mar-2019-3-5-52.272 | UP000029527 |
| 7 | Burkholderia pseudomallei K96243 | 145.5927 | 2.1631 | 14928 / 23961 | dbspec-15-Mar-2019-3-1-58.302 | UP000000605 |
| 8 | Burkholderia pseudomallei | 145.5352 | 2.163 | 14928 / 24630 | dbspec-15-Mar-2019-3-6-4.424 | UP000032603 |
| 9 | Burkholderia pseudomallei | 145.4263 | 2.1626 | 14928 / 24832 | dbspec-15-Mar-2019-3-5-56.499 | UP000030449 |
| 10 | Burkholderia pseudomallei | 145.4149 | 2.1626 | 14928 / 24475 | dbspec-15-Mar-2019-3-3-10.094 | UP000004875 |
| 11 | Burkholderia pseudomallei 1026b | 144.5794 | 2.1601 | 14928 / 23916 | dbspec-15-Mar-2019-3-4-15.005 | UP000010087 |
| 12 | Burkholderia pseudomallei | 144.4314 | 2.1597 | 14928 / 24032 | dbspec-15-Mar-2019-3-3-23.479 | UP000005700 |
| 13 | Burkholderia pseudomallei 1106a | 144.413 | 2.1596 | 14928 / 24105 | dbspec-15-Mar-2019-3-3-42.854 | UP000006738 |
| 14 | Burkholderia thailandensis | 143.7268 | 2.1575 | 14928 / 23446 | dbspec-15-Mar-2019-3-8-58.458 | UP000218714 |
| 15 | Burkholderia pseudomallei | 143.6768 | 2.1574 | 14928 / 23842 | dbspec-15-Mar-2019-3-3-0.858 | UP000004448 |
| 16 | Burkholderia thailandensis ATCC 700388 / DSM 13276 / CIP 106301 / E264 | 143.4335 | 2.1567 | 14928 / 23226 | dbspec-15-Mar-2019-3-2-18.411 | UP000001930 |
| 17 | Burkholderia thailandensis | 142.7679 | 2.1546 | 14928 / 22799 | dbspec-15-Mar-2019-3-9-35.914 | UP000235972 |
| 18 | Burkholderia pseudomallei | 141.464 | 2.1506 | 14928 / 22928 | dbspec-15-Mar-2019-3-2-35.82 | UP000002781 |
| 19 | Burkholderia mallei | 138.555 | 2.1416 | 14928 / 19432 | dbspec-15-Mar-2019-3-2-54.15 | UP000003995 |
| 20 | Burkholderia mallei Pseudomonas mallei | 138.5064 | 2.1415 | 14928 / 19381 | dbspec-15-Mar-2019-3-10-40.934 | UP000254505 |
| 21 | Burkholderia mallei NCTC 10229 | 138.0028 | 2.1399 | 14928 / 18362 | dbspec-15-Mar-2019-3-2-26.164 | UP000002283 |
| 22 | Burkholderia pseudomallei | 137.6652 | 2.1388 | 14928 / 21894 | dbspec-15-Mar-2019-3-3-7.02 | UP000004700 |
| 23 | Burkholderia mallei ATCC 23344 | 137.6361 | 2.1387 | 14928 / 18442 | dbspec-15-Mar-2019-3-3-41.84 | UP000006693 |
| 24 | Burkholderia cepacia | 131.5169 | 2.119 | 14928 / 29972 | dbspec-15-Mar-2019-3-10-52.322 | UP000263019 |
| 25 | Burkholderia lata ATCC 17760 / DSM 23089 / LMG 22485 / NCIMB 9086 / R18194 / 383 | 130.3785 | 2.1152 | 14928 / 29677 | dbspec-15-Mar-2019-3-6-12.489 | UP000036081 |
| 26 | Pseudomonas mesoacidophila | 128.9454 | 2.1104 | 14928 / 27324 | dbspec-15-Mar-2019-3-8-56.913 | UP000217994 |
| 27 | Burkholderia puraquae | 128.9136 | 2.1103 | 14928 / 28790 | dbspec-15-Mar-2019-3-8-4.997 | UP000193146 |
| 28 | Burkholderia stagnalis | 128.8023 | 2.1099 | 14928 / 26280 | dbspec-15-Mar-2019-3-6-40.413 | UP000068603 |
| 29 | Burkholderia contaminans | 128.7292 | 2.1097 | 14928 / 28983 | dbspec-15-Mar-2019-3-6-11.491 | UP000035664 |
| 30 | Burkholderia ubonensis | 128.5981 | 2.1092 | 14928 / 27094 | dbspec-15-Mar-2019-3-6-35.016 | UP000058930 |
